# Next-generation plasmids for transgenesis in zebrafish and beyond

**DOI:** 10.1101/2022.12.13.520107

**Authors:** Cassie L. Kemmler, Hannah R. Moran, Brooke F. Murray, Aaron Scoresby, John R. Klem, Rachel L. Eckert, Elizabeth Lepovsky, Sylvain Bertho, Susan Nieuwenhuize, Sibylle Burger, Gianluca D’Agati, Charles Betz, Ann-Christin Puller, Anastasia Felker, Karolína Ditrychová, Seraina Bötschi, Markus Affolter, Nicolas Rohner, C. Ben Lovely, Kristen M. Kwan, Alexa Burger, Christian Mosimann

## Abstract

Transgenesis is an essential technique for any genetic model. Tol2-based transgenesis paired with Gateway-compatible vector collections has transformed zebrafish transgenesis with an accessible, modular system. Here, we established several next-generation transgenesis tools for zebrafish and other species to expand and enhance transgenic applications. To facilitate gene-regulatory element testing, we generated Gateway middle entry vectors harboring the small mouse *beta-globin* minimal promoter coupled to several fluorophores, CreERT2, and Gal4. To extend the color spectrum for transgenic applications, we established middle entry vectors encoding the bright, blue-fluorescent protein mCerulean and mApple as an alternative red fluorophore. We present a series of p2A peptide-based 3’ vectors with different fluorophores and subcellular localizations to co-label cells expressing proteins of interest. Lastly, we established Tol2 destination vectors carrying the zebrafish *exorh* promoter driving different fluorophores as a pineal gland-specific transgenesis marker active prior to hatching and through adulthood. e*xorh*-based reporters and transgenesis markers also drive specific pineal gland expression in the eye-less cavefish (*Astyanax*). Together, our vectors provide versatile reagents for transgenesis applications in zebrafish, cavefish, and other models.

## Introduction

Transgenesis is a central technique across model systems for investigating biological processes *in vivo*. Integration of a transgene into the genome of a model organism enables experiments including live imaging, lineage tracing, generating conditional mutants, and disease modelling. Transgenic reporters permit labelling of cells, lineages, and structures *in vivo* using fluorescent proteins, making transgenesis an imperative tool in the study of developmental processes. Transgenesis can be paired with spatio-temporally controlled or ectopic gene expression to investigate the role of a particular gene within a molecular pathway. The zebrafish is an ideal system for transgenic applications due to its rapid development, optically transparent embryos, and amenability to genetic manipulation. Numerous additions to the toolbox for creating transgenic lines in zebrafish and improving transferability of expression constructs between species have critically enabled the study of evolutionarily conserved genes and their role in developmental processes and disease progression (Carney and Mosimann, 2018; Fisher et al., 2006a; Kawakami, 2004; Kawakami, 2005; Kikuta and Kawakami, 2009; White et al., 2013).

In particular, Tol2-based transgenesis and accessible vectors for its application have transformed the zebrafish community by providing a simple, modular system for the efficient generation of transgene constructs (Kawakami, 2004; Kawakami, 2005). Tol2-based transgenesis requires the delivery of transposase mRNA and a transgenesis vector containing a desired transgene cargo flanked by Tol2 transposon repeats, resulting in efficient random integration into the genome (Kawakami, 2005; Kawakami et al., 1998). This workflow is particularly suitable for use in zebrafish, which are injected with Tol2 transposase-encoding mRNA and a transgenesis vector at the one-cell stage (Kawakami, 2005; Kawakami, 2007; Kikuta and Kawakami, 2009; Urasaki et al., 2008). Originally derived from medaka, Tol2 transposon activity for transgenesis is not restricted to zebrafish, as injection or electroporation-based delivery methods have been successfully used in several fish species and in other vertebrate models including *Amphioxus, Xenopus*, and chicken embryos (Hamlet et al., 2006; Kozmikova and Kozmik, 2015; Sato et al., 2007).

Despite its versatility, taking full advantage of Tol2 transgenesis hinges upon the broad availability, applicability, and functionality of its required components. The first iteration of the Tol2kit (Kwan et al., 2007) and other powerful plasmid collections (Don et al., 2017; Villefranc et al., 2007) provided several entry and destination vectors for Multisite Gateway cloning (Cheo et al., 2004; Hartley et al., 2000) or restriction enzyme-based assembly of *[regulatory element(s)]–[effector coding sequence]–[3′ trailer]* constructs into a *Tol2* transposon-flanked backbone. However, as transgenesis applications have become more widely applied and complex, an expanded set of transgenesis components has become essential.

First, transgenes based on isolated regulatory elements need a basic complement of validated parts to constitute a gene expression unit. Most prominent is the need for a minimal promoter sequence that is inert to position effects upon random Tol2-based integration and that features reproducible, robust interaction with numerous enhancer elements (Carney and Mosimann, 2018; Felker and Mosimann, 2016). Second, the widespread use of EGFP, mCherry, and other fluorophores that generate red and green fluorescent reporters has introduced a challenge in the generation of combinatorial transgenic lines (Don et al., 2017; Kwan et al., 2007; Villefranc et al., 2007). Third, marking the expression of a transgene carrying a gene of interest as a fusion protein is not always practical, and previous attempts to use internal ribosomal entry sites (IRES) sequences to generate bi-cistronic reporters have been generally unsuccessful in zebrafish (Kwan et al., 2007). Lastly, the limited collection of available destination backbones with validated transgenesis markers restricts combinatorial transgene use and at times limits observations in tissues dominated by common transgenesis markers such as the heart (promoter of *myl7*, formerly called *cmlc2*) and the eye lens (various *crystallin* gene promoters) (Hesselson et al., 2009; Huang et al., 2003; Kwan et al., 2007; Villefranc et al., 2007).

To expand on previously available transgenesis tool sets including the Tol2kit (Kwan et al., 2007), we embarked on a collaborative effort to augment, diversify, and increase the functionality of available plasmid sets for transgenesis. Building upon the work of many before us (including, but not restricted to, Amsterdam et al., 1995; Asaoka et al., 2002; Bessa et al., 2009; Distel et al., 2009; Fisher et al., 2006b; Hesselson et al., 2009; Huang et al., 2003; Kawakami et al., 2000; Koga et al., 1996; Mosimann et al., 2011; Tamplin et al., 2011; Woolfe et al., 2004) and with the goal of accessible documentation and distribution to the community, we here present several plasmid vectors compatible with MultiSite Gateway and traditional restriction enzyme cloning. These constructs are geared towards facilitating gene-regulatory element discovery, transgene quality control, reporter analysis, and an expanded fluorophore spectrum for imaging. While the here described new backbones are geared towards Tol2-based transgenesis in zebrafish, the individual modules are deployable for numerous applications in diverse species, such as cavefish. Together, our vectors (sequences in GenBank format and PDF maps provided as **Supplemental Data File 1**) expand the available modules for reproducible, quality-controlled Tol2 transgenesis and gene-regulatory element testing in zebrafish and other model systems.

## Results

### The mouse *beta-globin* (*Hbb-bt*) minimal promoter for universal enhancer testing

The discovery and isolation of gene-regulatory elements, and especially of distal enhancer elements, has been instrumental in decoding gene-regulatory input and in the creation of new transgenic tools to label and manipulate cell types of choice. Enhancers require a minimal promoter region associated with the transgene reporter to initiate RNA Pol II-based transcription (Haberle and Stark, 2018). Since reporter expression can be positively and negatively influenced by neighboring loci in the genome (Amsterdam et al., 1995; Jaenisch et al., 1981; Lalonde et al., 2022; Wilson et al., 1990), an expression construct designed to test an enhancer requires an inert minimal promoter that is not sensitive to other enhancers in the vicinity of the transgene insertion. Several minimal promoters have been applied for transgenesis in zebrafish, including *Ef1a, hsp70l, c-fos, krt4, TK, gata2a* promoter regions that are several hundred to several kilo-bases in length (Amsterdam et al., 1995; Bessa et al., 2009; Johnson and Krieg, 1995; Kondrychyn et al., 2009; Meng et al., 1997; Ogura et al., 2009; Scott et al., 2007; Shoji and Sato-Maeda, 2008; Weber and Köster, 2013).

The promoter activity of the mouse *beta-globin* minimal promoter (*Hbb-bt* gene, 129 base pair sequence including 54 bases of 5’ UTR) was initially characterized by mutagenizing the regulatory region and measuring the resulting transcripts in HeLa cells (Myers et al., 1986). Coupling of the mouse *beta-globin* minimal promoter to a promoter-less enhancer enables the generation of stable Tol2-based zebrafish transgenics with tissue-specific reporter activity, including fluorescent reporters and CreERT2 and KalTA4 driver lines (D’Agati et al., 2019; Kaufman et al., 2016; Prummel et al., 2019; Tamplin et al., 2011; Woolfe et al., 2004). Reporters including the mouse *beta-globin* minimal promoter show minimal to no background in transient injections and drive stable transgene activity in zebrafish, including when paired with enhancers from other species (Prummel et al., 2019; Tamplin et al., 2011; Tamplin et al., 2015; Woolfe et al., 2004). To enable testing and application of candidate enhancers in 5’ entry vectors, we have generated middle entry vectors with mouse *beta-globin* minimal promoter-coupled m*Cerulean, EGFP, mCherry*, and *mApple* ORFs, as well as *creERT2* and codon-optimized *Gal4* (*KalTA4*) (Distel et al., 2009) (**Fig. 1A**).

**Figure 1:**
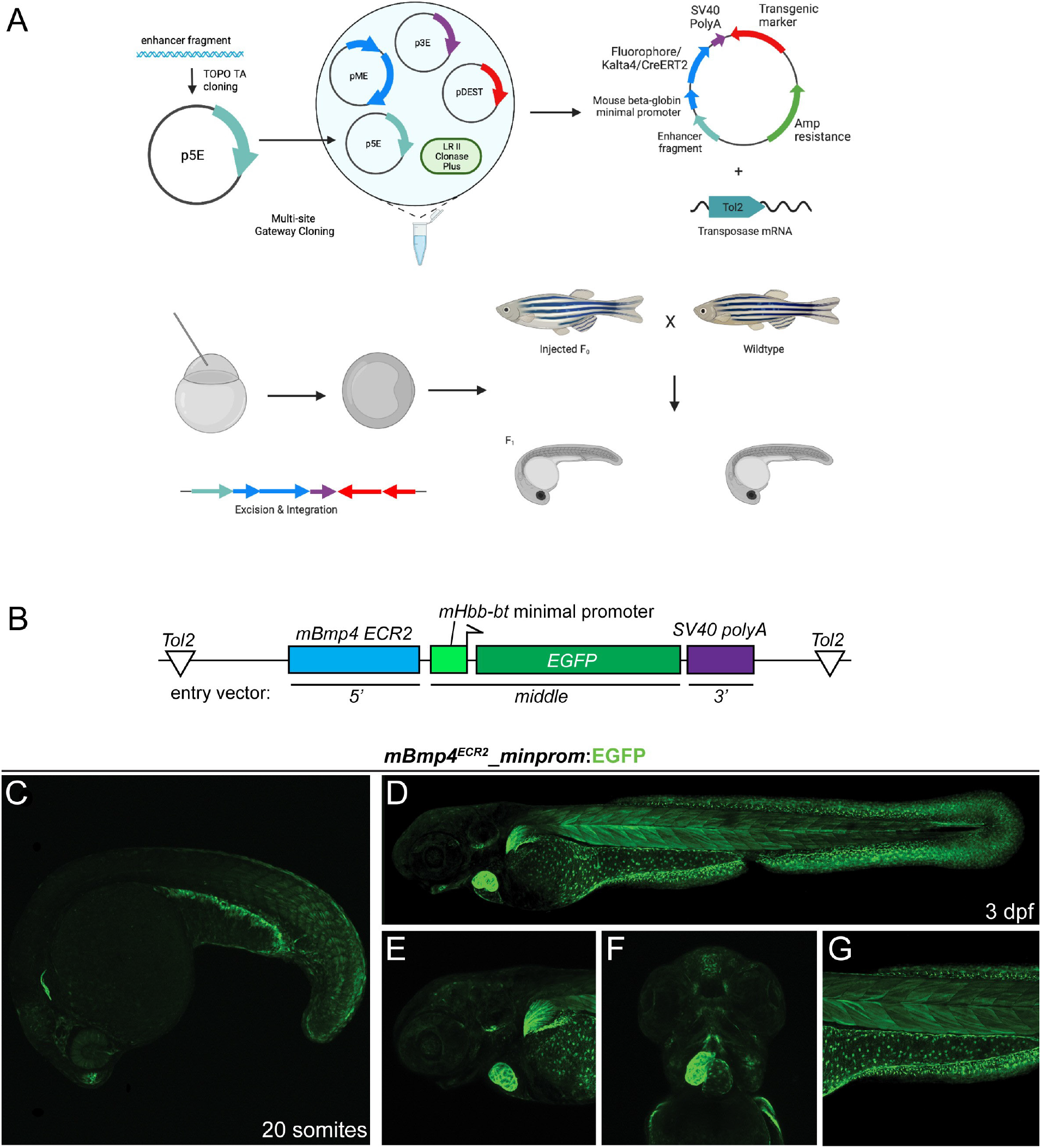
Incorporating the small mouse *beta-globin* minimal promoter into transgenic reporters. (**A**) Workflow of reporter generation or enhancer testing with Multisite Gateway recombination and Tol2 transgenesis. Enhancer fragment is cloned into a *p5E* vector and Gateway- or restriction enzyme-cloned into an expression construct composed of 5’ fragment, middle entry mouse *beta-globin minimal promoter*:*fluorophore/KalTa4/creERT2*, 3’ *SV40 polyA*, transgenesis marker (optional). The resulting Tol2 expression construct is injected along with Tol2 transposase-encoding RNA into one-cell stage zebrafish, randomly integrated into the genome, screened to establish transgenic lines. Schematic generated using BioRender. (**B**) Schematic of *mBmp4*^*ECR2*^*:minprom:EGFP* construct containing the distal 668 bp mouse *BMP4 ECR2* element, the 129 bp mouse *beta-globin* minimal promoter driving *EGFP* and the *SV40 polyA*. (**C-G**) the *mBmp4*^*ECR2*^ reporter expresses in a variety of zebrafish tissues linked to endogenous *bmp4* expression. Lateral view of stable transgenic line of *mBmp4*^*ECR2*^*:minprom:EGFP* at 20 somite stage (**C**). Lateral (**D,E,G**) and ventral (**F**) views of 3 dpf stable *mBmp4*^*ECR2*^*:minprom:EGFP* transgenic line (**D**) show expression in the heart and pectoral fin (**E**), parts of the head musculature (**F**), and in the mesothelium surrounding the yolk as well as trunk musculature and fin fibroblasts (**G**).

To further illustrate the utility of the mouse *beta-globin* minimal promoter, we documented regulatory element testing of a mouse enhancer. The 668 bp-spanning downstream enhancer *ECR2* in the mouse *Bmp4* locus has previously been discovered to convey posterior lateral plate mesoderm activity when tested as a reporter in mouse embryos (Chandler et al., 2009). In zebrafish embryos, the *ECR2* enhancer coupled with *beta-globin-minprom:EGFP* (*mBmp4*^*ECR2*^*_minprom:EGFP*) (**Fig. 1B**) as stable transgene resulted in EGFP reporter expression in a dynamic expression pattern in different cell types, including trunk and tail muscles, pectoral fins, the heart, and the yolk-surrounding mesothelium (**Fig. 1C**). Notably, several of these reporter expression domains matched with the reported expression of zebrafish *bmp4* as established by mRNA *in situ* hybridization (**Fig. 1D-G**) (Bisgrove et al., 2000; Lenhart et al., 2013; Nikaido et al., 1997; Stickney et al., 2007; Thisse and Thisse, 2005). Together with prior work applying this small promoter, these data illustrate the utility of the mouse *beta-globin* minimal promoter for zebrafish transgenes based on isolated regulatory elements.

### Components for fluorescent reporter generation

To expand available basic components for transgenic reporter generation, we revisited available fluorophores and *3’* UTR constructs. The prevalent use of green fluorescent avGFP derivatives as well as red-fluorescent proteins (Chalfie et al., 1994; Rodriguez et al., 2017; Tsien, 2003) in the zebrafish field has greatly expanded the number of available transgenic reporters, yet has led to a bottleneck in combining transgenic reporters for multi-color imaging. In addition, the application of fluorophores that have not been extensively used remains challenging, due to possible toxicity or other experimental issues when embarking on a lengthy transgenesis experiment. To extend the color spectrum available for transgenic applications with accessible tools, we have generated middle entry clones with cytoplasmic, membrane, and nuclear versions of mCerulean (**Fig. 2A-F**). mCerulean is a monomeric, blue-fluorescent protein derived from avGFP and subsequent CFP with an excitation wavelength of 433 nm and an emission wavelength of 475 nm (Rizzo and Piston, 2005; Rizzo et al., 2004). While other blue fluorophores are spectrally more distant to the commonly used green-fluorescent GFP derivatives, mCerulean can be detected using standard GFP dissecting scope setups. Transgenes using mCerulean as reporter are well-tolerated in zebrafish (**Fig. 2A**) (Hesselson et al., 2009; Trinh et al., 2011; Zuniga et al., 2011). Spectral separation of mCerulean from GFP is easily achievable on common confocal and light sheet setups, either by using a 445 nm laser or using features of typical confocal software platforms to minimize spectral overlap (such as in Zeiss’ ZEN software). We noted that depending on genetic background, mCerulean imaging on a scanning confocal microscope resulted in autofluorescence of skin or pigment cells, particularly in the head region of zebrafish embryos and larvae (48 hpf onwards) when visualized with an Argon laser (488 nm) (**Fig. 2A-F**). Use of the ideal excitation laser wavelengths greatly diminished this autofluorescence and we have not found any significant problems with mCerulean reporter imaging at these stages. To demonstrate the versatility of mCerulean to visualize different embryonic structures, we imaged the transgenic reporter line *Tg(Hs-HOPX*^*4q12N3*.*1*^*:minprom_mCerulean)* (subsequently abbreviated as *HOPX:mCerulean). HOPX:mCerulean* includes the upstream region of the human *HOPX* gene together with the mouse *beta-globin* minimal promoter and resulted in stable expression in the notochord at 48 hpf (**Fig. 2A**). This expression is in line with previous work that established the expression of *HOPX* in the developing primitive streak as well as expression in chordoma tumors (Loh et al., 2016; Nelson et al., 2012; Palpant et al., 2017). Notably, the human *HOPX* upstream region also drives expression in rhombomeres 3 and 5 in individual transgenic insertions of the reporter (Chen et al., 2002). As a blue fluorescent reporter, *HOPX:mCerulean* combines well with mCherry-based reporters such as *drl:mCherry* that marks lateral plate mesoderm before restricting expression to cardiovascular lineages (**Fig. 2A**) (Mosimann et al., 2015; Sánchez-Iranzo et al., 2018). Together, mCerulean provides a practical alternative fluorophore to expand the reporter spectrum in transgenic experiments.

**Figure 2:**
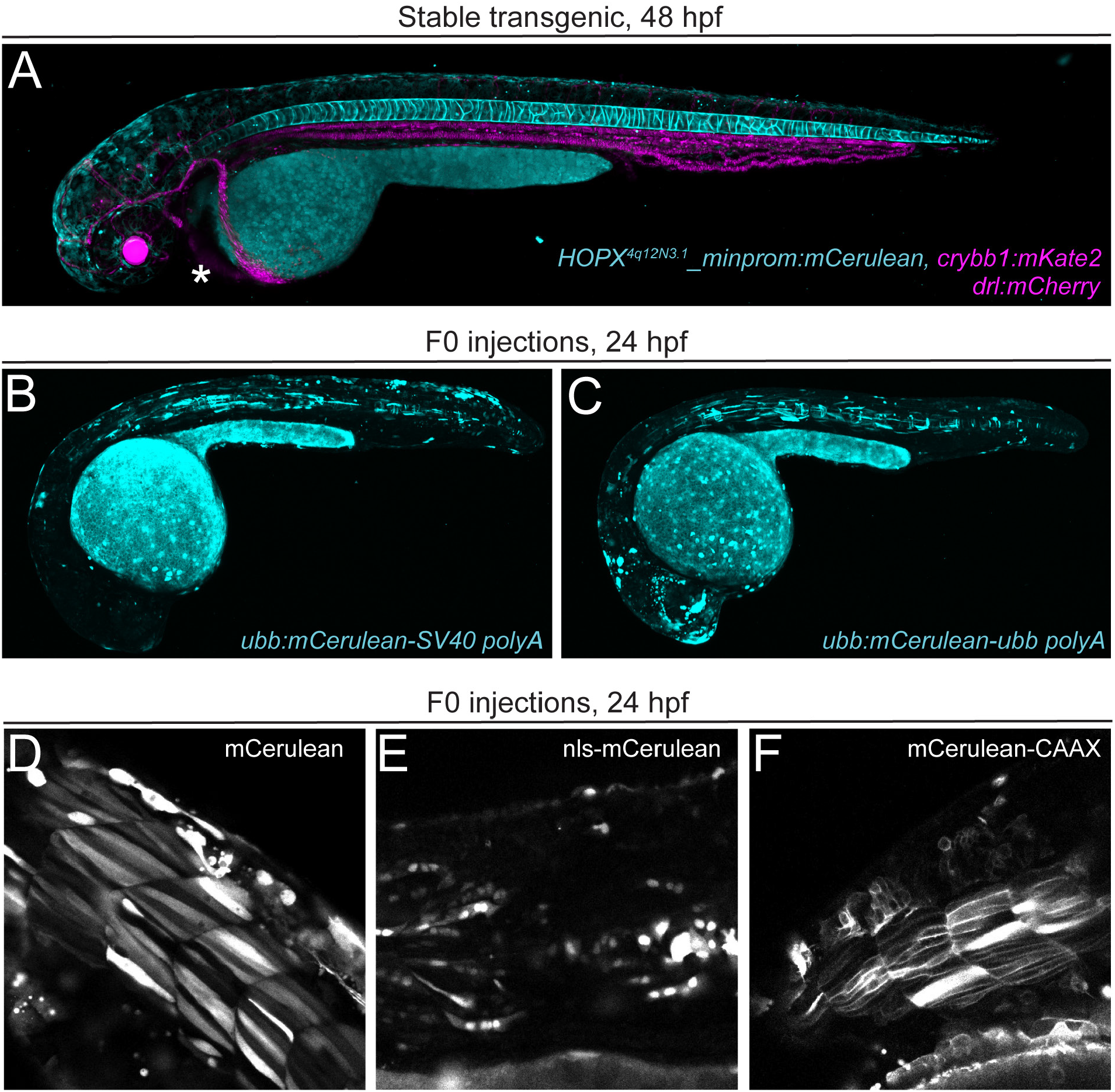
mCerulean as versatile blue fluorophore for zebrafish transgenesis. (**A**) Stable transgenic line for *HOPX4q12N3*.*1:minprom_mCerulean,crybb1:mKate2*, shows mCerulean expression in both the vacuolar and sheath cells of the zebrafish notochord at 2 days post-fertilization and eye lens-specific mKate2 expression (asterisk) when crossed with *drl:mCherry* transgenic, marking lateral plate mesoderm-derived lineages (cardiovascular system, blood at 2 dpf). (**B,C**) Transient injection of zebrafish *ubiquitin* (*ubb*) promoter-driven mCerulean with 3’ polyadenylation trailer (*polyA*) from *ubb* (**B**) or *SV40* (**C**). (**D-F**) mCerulean constructs for different cellular localization. Transient expression of cytoplasmic mCerulean (**D**), nuclear nls-mCerulean (**E**), and membrane-bound mCerulean-CAAX (**F**).

Transgene constructs require a 3’ UTR with polyadenylation signal (polyA) for productive mRNA transcription, stability, and translation (Felker and Mosimann, 2016; Fink et al., 2006; Matoulkova et al., 2012; Okayama and Berg, 1992). The most common 3’ UTR with polyA sequences used for zebrafish transgenes are the bi-directional *SV40 late polyA* sequence (abbreviated as *SV40 polyA*) (Okayama and Berg, 1992) that was included in the original Tol2kit collection (vector *#302*) (Kwan et al., 2007) and the *BGH* polyA sequence (Higashijima et al., 1997). Several available *Tol2* backbones for MultiSite Gateway assembly contain the *SV40 polyA* after the recombination cassette (Kwan et al., 2007), which can lead to assembly and sequencing issues when another *SV40 polyA* trailer is used in the 3’ position of the assembly. To expand the available 3’ UTRs for transgene assembly, we isolated a 516 bp fragment of the 3’ UTR of the zebrafish *ubiquitin B* (*ubb, ubi*) gene that we transferred into the *p3E* vector backbone, resulting in *p3E_ubb-polyA*. To test the *p3E_ubb-polyA* and compare it to the commonly used Tol2kit vector *#302*, we generated *pCK005 ubi:mCerulean-ubb-polyA* as well as *pCK004 ubi:mCerulean-SV40-polyA*, which provide convenient control vectors for co-injection along with newly generated reporter constructs of unknown activity (**Fig. 2B,C**). In addition to this alternative *SV40 polyA p3E* vector, we also generated an ‘empty’ *p3E* vector that contains the multiple cloning site from *pBluescript KSII(+)*, providing single-cutter restriction sites for cloning inserts into the final plasmid (see Methods for details).

mCherry and multimeric dsRED versions are commonly applied red fluorophores for zebrafish transgenesis (Davison et al., 2007; Kwan et al., 2007; Pisharath et al., 2007; Provost et al., 2007; Rodriguez et al., 2017; Traver et al., 2004). mApple (derived from *Discosoma sp*.) is a fast-maturing, monomeric fluorophore with an excitation maximum of 568 nm, therefore less red-shifted than mCherry and still excitable by standard lasers used with confocal microscopes (543 or 561 nm) (**Fig. 3A-G**) (Shaner et al., 2008). mApple has a better theoretical quantum yield and a fast-folding mechanism, rendering it a potentially suitable, additional red monomeric fluorophore for any transgenesis toolkit. mApple is well-tolerated in zebrafish and has been successfully pioneered for transgenic applications (Heim et al., 2014; Lou et al., 2015), yet remains less frequently used compared to mCherry or dsRED derivatives. Applied to label endoderm membranes, mApple expression in stable *Tg(sox17:mApple-CAAX)* showed faithful and well-detectable expression (**Fig. 3A,B**) akin to *Tg(sox17:EGFP-CAAX)* (**Fig. 3C,D**), complementing the existing and widely used cytoplasmic *sox17:EGFP* to label endoderm progenitors (Chung and Stainier, 2008; Sakaguchi et al., 2006). To expand the application of mApple in zebrafish, we further generated combinations of *mApple* with the mouse *beta-globin* minimal promoter as middle entry clones (see Methods for details) with nuclear as well as membrane-localized versions (**Fig. 3E-G**).

**Figure 3:**
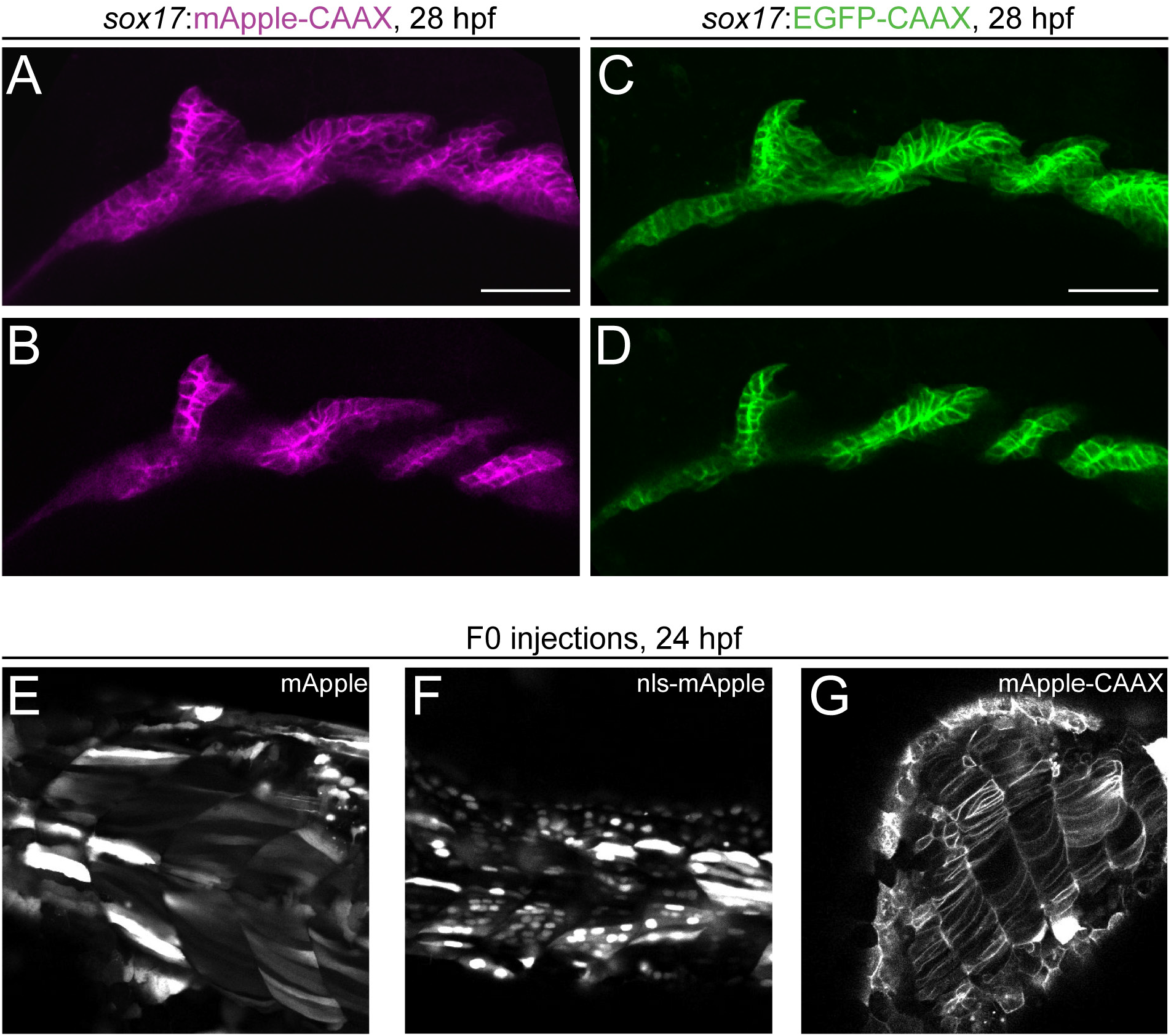
mApple as bright red fluorophore for zebrafish transgenesis. (**A-D**) Comparison of membrane-localized mApple-CAAX with EGFP-CAAX in stable s*ox17:mAppleCAAX and sox17:EGFPCAAX* lines show matching expression in the developing pharyngeal endoderm at 28 hpf. Pharyngeal pouches shown as close-up with anterior to the left in max projection (**A,C**) and single slice views (**B,D**); scale bars 50 μm. (**E-G**) mApple fusion proteins for different localizations. Transient expression of cytoplasmic mApple (**E**), nuclear nls-mApple (**F**), and membrane bound mApple-CAAX (**G**).

Together, our series of minimal promoter-coupled fluorophore vectors, multiple localization tags, plus *p3E polyA* with multiple cloning site vectors provide a basic set of reagents for versatile regulatory element discovery and simple fluorescent microscopy with standard equipment accessible in the field.

### 2A peptide clones for fusion protein reporters

A key technique in transgenesis is driving expression of a protein of interest in a specific cell population, an approach that greatly benefits from fluorescently labeling transgene-expressing cells throughout development. One method to directly label the expressed protein of interest is a fusion with a fluorescent tag; yet the impact of adding a large fluorescent protein to the protein of interest can result in mis-localization, sequestration, degradation, or inactivity of the fusion product (Giepmans et al., 2006). A widely deployed alternative method is the translation of the protein of interest and a fluorescent reporter via a bi-cistronic message (Chan et al., 2011; Trichas et al., 2008). We previously released a series of vectors including *IRES* (*internal ribosome entry site*) sequences (Jang et al., 1988; Pelletier and Sonenberg, 1988) for such applications. However, *IRES*-containing constructs have been repeatedly reported to feature variable expression of the ORF (open reading frame) downstream of the *IRES* sequence (Ibrahimi et al., 2009; Wong et al., 2002). With *IRES*-containing constructs used in zebrafish, the ORF following the IRES (often a fluorescent reporter) is commonly expressed at a much lower level than the first ORF. Individual users observed up to a 10-fold lower expression of the second ORF, while other data indicate that the presence of the *IRES* reporter can even dampen expression of the first ORF (Weber and Köster, 2013).

To provide an alternative strategy for fluorescent labeling of gene of interest-expressing cells, we have established 3’ entry (*p3E*) vectors that include the PTV-1 2A peptide sequence, a viral peptide that results in stoichiometric production of two proteins from the same message (Kim et al., 2011; Provost et al., 2007; Ryan et al., 1991). 2A-mediated “self-cleavage” takes advantage of the ribosome not forming a glycyl-prolyl peptide bond at the C-terminus of the 2A peptide, allowing the translational complex to release the first polypeptide and continue translation of the downstream product without independent ribosome recruitment or additional host factors (Donnelly et al., 2001a; Donnelly et al., 2001b) (**Fig. 4A**). The resulting equimolar protein amounts make the 2A peptide an ideal tool for the quantification of transgene expression or the controlled expression of multiple proteins from one transgene (Chi et al., 2008; Knopf et al., 2010; Knopf et al., 2011). We generated 3’ entry constructs containing the 2A peptide-encoding sequence followed by different fluorophores (EGFP, mCherry, and mCerulean) in cytoplasmic, membrane, and nuclear forms (**Fig. 4B-D**). These constructs are all designed in a Gateway-compatible reading frame (see Methods and Gateway manual for details on primer design) and are ready for use with middle entry clones that are in Gateway reading frame without a stop codon.

**Figure 4:**
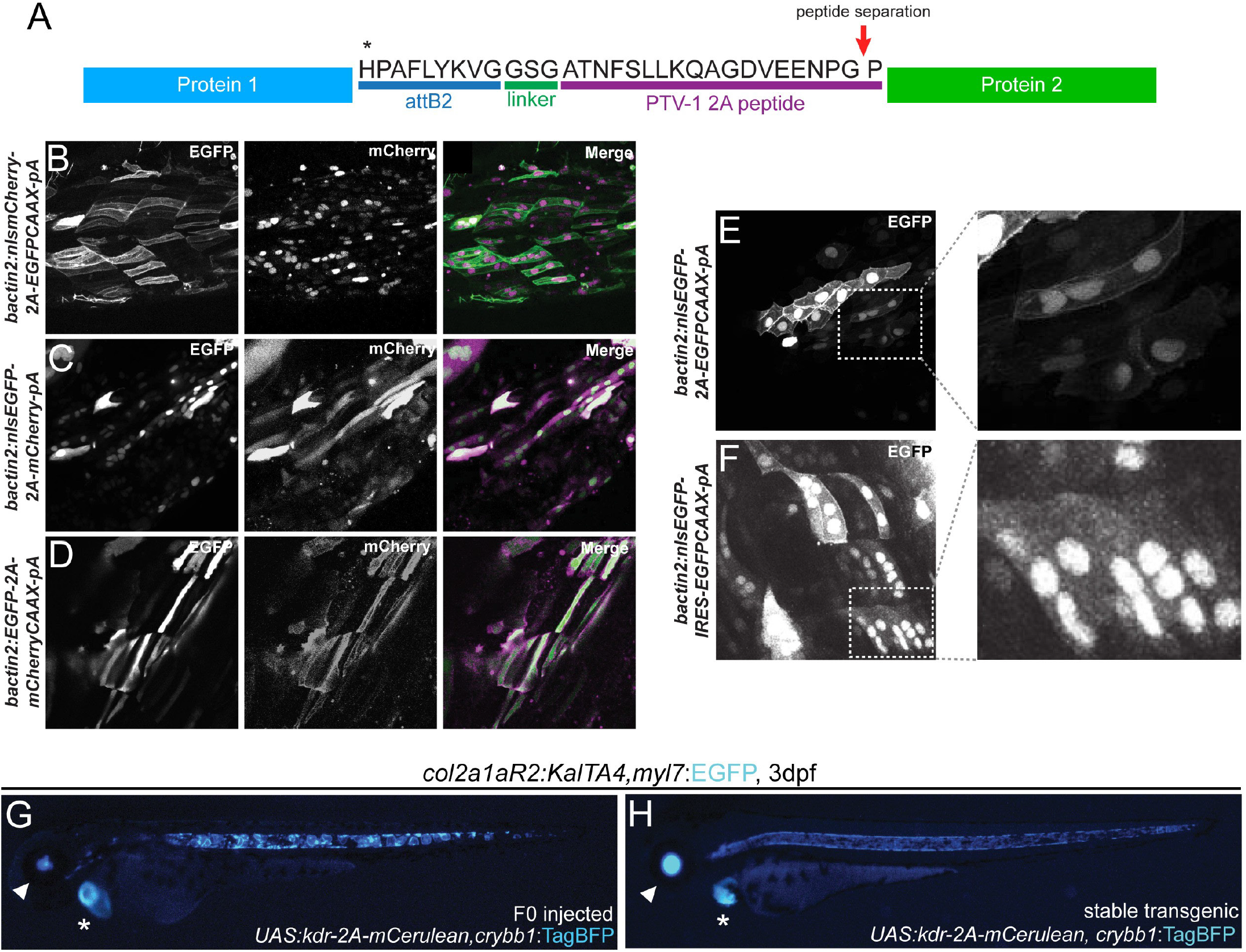
2A peptide linkers for fluorescent protein tagging. (**A**) Schematic of protein product resulting from a Tol2 construct containing two protein sequences separated by a 2A peptide following the *attB2* site of a Multisite Gateway expression construct. Translation of the first protein occurs, the first polypeptide is released at the site of peptide separation, and translation of the second ORF proceeds with the same ribosome resulting in equimolar protein amounts. (**B-F**) Transient injections of bactin2 promoter-driven fluorophore combinations separated by a 2A peptide tag; *nlsmCherry-2A-EGFPCAAX* (**B**), *nlsEGFP-2A-mCherry* (**C**), and *EGFP-2A-mCherryCAAX* (**D**). Transient injection of *bactin2:nlsEGFP-2A-EGFPCAAX* (**E**) demonstrates expression of EGFP in both the nucleus and membrane of cells whereas injection of *bactin2:nlsEGFP-IRES-EGFPCAAX* (**F**) results in poor EGFP-CAAX detection in cells positive for nuclear EGFP. (**G,H**) Use of 2A fusions in Gal4/*UAS* transgene experiments. Injection of *UAS:kdr-2A-mCerulean,crybb1:TagBFP* into stable transgenic *col2a1aR2:KalTA4,myl7:EGFP* that expresses codon-optimized Gal4 in the developing notochord and subsequent cartilage lineages shows mosaic mCerulean expression in the developing notochord (**G**), with the secondary *myl7:EGFP* transgenic marker noted with an asterisk and the *crybb1:TagBFP* marker noted with an arrow. Homogenous mCerulean notochord expression can be seen in stable F2 *UAS:kdr-2A-mCerulean,crybb1:TagBFP* crossed with *col2a1aR2:KalTA4, myl7:EGFP* (**H**).

To evaluate and compare stoichiometry of protein production using the 2A peptide and the IRES, we generated transgene expression constructs in which the same fluorophore (EGFP) was targeted to two different subcellular locations in transient injections: nuclear-localized EGFP (nls-EGFP) was in the first position and membrane-targeted EGFP (EGFP-CAAX) was in the second (**Fig. 4E**). When separated by the 2A peptide, EGFP was easily visualized in both nuclear and membrane locations, even in cells expressing low levels of the transgene (**Fig. 4E**). When separated by the IRES, in most cells, only nuclear-localized EGFP was detectable; EGFP-CAAX was only detected in the brightest cells, confirming that the 2A peptide yields effective production of two separate proteins targeted to distinct subcellular locations (**Fig. 4F**).

The utility of 2A-based fusions becomes apparent when applied with the Gal4/*UAS* system. Gal4-dependent expression of genes of interest as UAS-controlled transgenes is a powerful approach to test individual gene function and for disease modeling (Akitake et al., 2011; Distel et al., 2009; Halpern et al., 2008; Köster and Fraser, 2001; Provost et al., 2007). The ability to fluorescently label the *UAS* transgene-expressing cells provides an immediate phenotype readout; as 2A-based fusions harbor the fluorescent reporter C-terminally of the ORF, the fluorescence signal additionally provides an expression control of the entire coding sequence. Injection of *UAS:kdr-2A-Cerulean* into the cartilage-specific Gal4-driver transgenic line *col2a1aR2:KalTA4* (D’Agati et al., 2019; Dale and Topczewski, 2011) resulted in mosaic Cerulean expression in the developing notochord (**Fig. 4G**). Stable *UAS:kdr-2A-Cerulean* transgenic insertions (selected for with the *crybb1:TagBFP* transgenesis marker *in cis* on the transgene backbone *pCB24*) crossed to the same driver resulted in homogenous notochord expression (**Fig. 4H**).

A major technical consideration remains when using the 2A peptide in the Gateway system as it will result in the addition of a 28 aa “tag” to the C-terminal end of the first protein (containing the translated *att* site and the 2A peptide) (Rothwell et al., 2010) (**Fig. 4A**). This residual amino acid sequence may disturb the protein of interest in the first position if it is sensitive to C-terminal tagging. To reverse the relative positions of the fluorophore and ORF of interest cloning, we also cloned a 3’ “empty” 2A entry vector containing the *MCS* from *pCS2* for cloning of genes of interest after the 2A peptide sequence that can be used with middle entry fluorescent reporters (see Methods for details). Together, our 2A-based vectors enable versatile tagging of proteins of interest in zebrafish transgenes and beyond.

### A selective, inert transgenic marker labeling the pineal gland

Transgenesis requires rigorous quality control, which is facilitated by inclusion of an easily-to-screen, dominant reporter gene *in cis* with the intended transgene cargo. A reliable transgenesis marker further supports regulatory element testing by ensuring positive selection for transgene carriers. In practical terms, inclusion of an independent fluorescent transgene that is detectable for screening at a desired developmental stage enables identification of transgene-positive animals and tracking of the transgenic insertion across generations (Felker and Mosimann, 2016). The initial Tol2kit contained the destination vectors *pDestTol2pA2* as “empty” destination vector and *pDestTol2CG2* that included the *myl7* (formerly *cmlc2)* promoter driving cytoplasmic EGFP in the developing myocardium from late somitogenesis stages on, easily screenable by 24 hpf (Huang et al., 2003; Kwan et al., 2007). However, *myl7*-based transgenesis markers are not desirable for transgenes focused on heart development, a significant interest in the field (Bakkers, 2011; Cao and Poss, 2018; Kemmler et al., 2021; Potts et al., 2021; Smith et al., 2021; Yao et al., 2021). Further, the heart becomes increasingly obscured by skin pigmentation in standard zebrafish strains, rendering transgene detection in adults challenging (White et al., 2008).

Eye lens-specific promoters of different *crystallin* genes have been used as powerful transgenesis markers (Davidson et al., 2003; Emelyanov and Parinov, 2008; Hesselson et al., 2009; Kurita et al., 2003; Marques et al., 2008; Waxman et al., 2008). *Alpha-* or *beta-crystallin* promoter-based transgenes (*cryaa* or *crybb1*) are visible throughout adulthood and are easily detected via UV flashlights and filter glasses (Felker and Mosimann, 2016). We previously adopted *cryaa:Venus* (Hesselson et al., 2009) to continuously mark effector transgenics such as CreERT2 drivers with a yellow fluorescent transgenic marker (Mosimann et al., 2015). To complement existing reporter backbones, we generated *pCB59 crybb1:mKate2* to serve as a red eye transgenesis marker (**Fig. 2A**) and *pCB24 crybb1:TagBFP* as a blue eye transgenesis marker (**Fig. 4G,H**). The far-red mKate2 (Lin et al., 2012; Shcherbo et al., 2009) is still detectable using standard red fluorescence filters on dissecting scopes with excitation at 588 nm and emission at 633 nm, and is complementary to blue, green, or yellow fluorescent *crystallin* promoter-driven markers for combinatorial transgene experiments. TagBFP is a robust blue fluorophore with excitation at 402 nm and emission at 457 nm (Subach et al., 2008) that is well-tolerated in zebrafish as well (Singh et al., 2012). One drawback of *crystallin*-based reporters is their expression initiates around 36-48 hpf (Emelyanov and Parinov, 2008; Hesselson et al., 2009; Kurita et al., 2003; Marques et al., 2008; Waxman et al., 2008), which is later than *myl7*-based reporters and close to the start of swimming that can render sorting of transgene-positive embryos challenging. Additionally, eye lens expression of fluorophores can interfere with imaging of head regions and is not desirable for labs and experiments focused on eye development.

To establish a transgenesis marker with a) early developmental expression to enable efficient screening, b) persistent fluorescent reporter expression that is unobtrusive to tissues of interest, c) focused expression in a small region in the zebrafish, and d) activity throughout the zebrafish life cycle, we sought to generate *Tol2* backbones with pineal gland-specific reporter expression. The deeply conserved pineal gland, pineal organ, or conarium is a small photosensitive endocrine gland forming at the anterior dorsal midline on the head and is involved in circadian rhythm control (Erlich and Apuzzo, 1985; Gheban et al., 2019). Importantly, in zebrafish the pineal gland condenses during somitogenesis, remains in place for the lifespan of the animal, is small, and at a different anatomical location compared to commonly used transgenesis markers (Asaoka et al., 2002; Kazimi and Cahill, 1999; Ziv et al., 2005).

The 1055 bp upstream region of the *exo-rhodopsin* (*exorh*) gene harbors a *pineal expression-promoting element* (*PIPE*) that drives specific, persistent transgene expression in the pineal organ as detectable by stereo microscopy from 22 somites on (Asaoka et al., 2002). We generated three destination vectors containing the *exorh* promoter driving *EGFP, mCherry*, and *mCerulean*, respectively (*pCK017 pDEST-exorh:EGFP, pCK053 pDEST-exorh:mCherry, pCK054 pDEST-exorh:mCerulean*) and tested their performance as transgenesis backbones (**Fig. 5A**).

**Figure 5:**
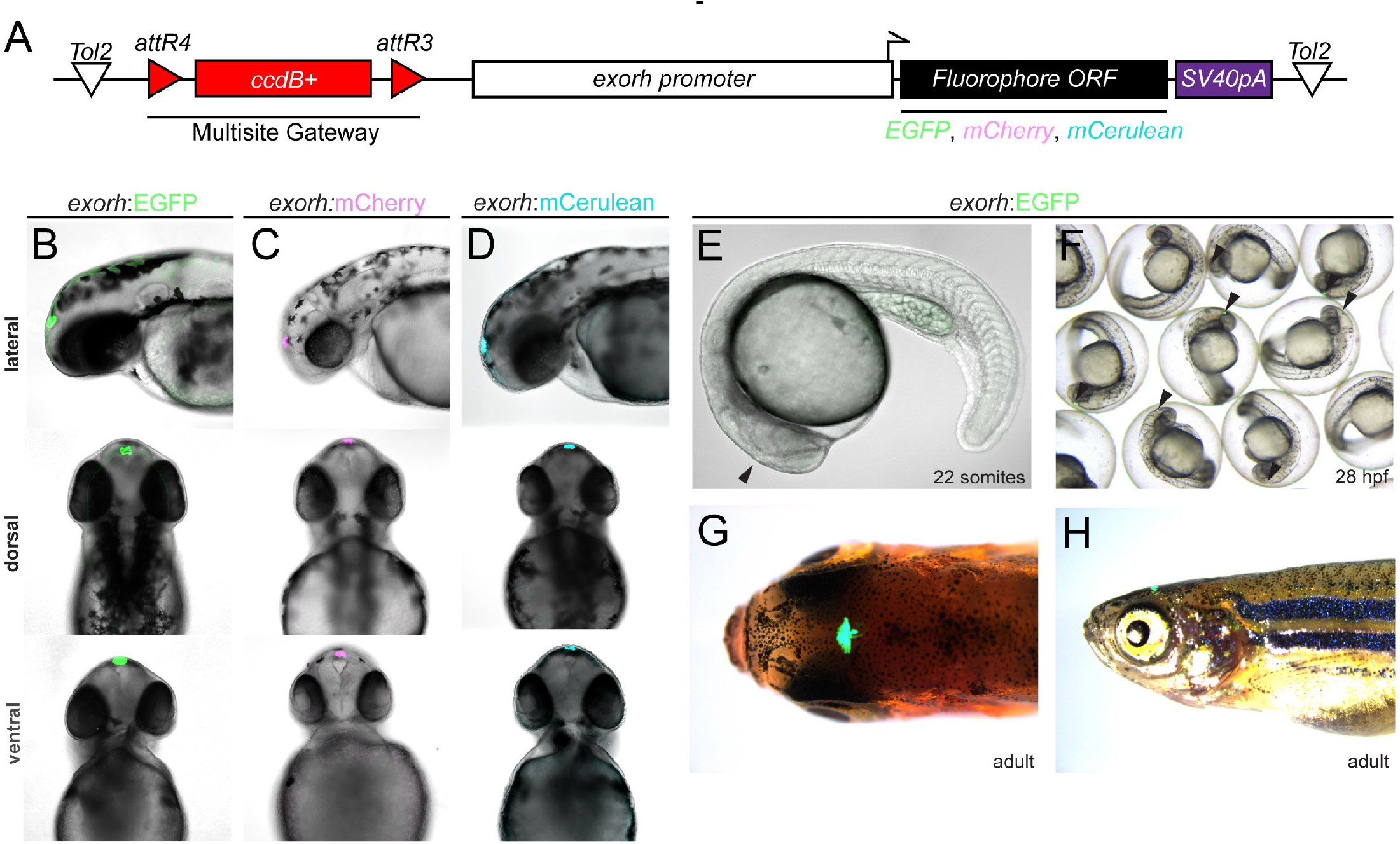
Labeling the zebrafish pineal gland with the *exorhodopsin* (*exorh*) regulatory region as transgenesis marker. (**A**) Schematic of the Tol2 secondary transgenic marker backbones *pCK017 pDEST-exorh:EGFP, pCK053 pDEST-exorh:mCherry-*, and *pCK054 pDEST-exorh:mCerulean* including the *attR4-* and *attR3*-flanked *ccdB* cassette followed by the1055 bp *exorh* promoter driving *EGFP, mCherry*, or *mCerulean*. (B-D) Stable transgenic lines for EGFP-, mCherry-, and mCerulean-based *exorh*-driven fluorophore reporters; lateral, dorsal, and ventral views at 2 dpf showing specific, easily detectable reporter activity in the pineal gland. (**E-H**) Stages and appearances of exorh-driven transgenesis reporters. Starting at 22 somite stage (**E**), *exorh:EGFP* expression (black arrowheads) is easily detectable in individual embryos and in groups under a standard dissecting scope (**F**) while embryos are still in their chorions. *exorh*-driven fluorophore expression persists into adulthood as seen in dorsal (**G**) and lateral views (**H**).

Upon injection with Tol2 transposase mRNA, all three reporter backbones resulted in prominent, robust, and reproducible pineal gland-restricted fluorophore expression starting from around 22 hpf, in line with the initial reports characterizing *exorh*-based reporters (**Fig. 5B-F**) (Asaoka et al., 2002). As stable transgenic insertions, *exorh:EGFP* recapitulated the transient expression (**Fig. 5B-D**), showing reporter expression that is easy to screen under a standard fluorescent dissecting scope from 24 hpf onwards while embryos remained in their chorions (**Fig. 5F**). We observed stable integration and expression of *exorh:EGFP* throughout zebrafish development and adulthood, when the pineal gland-specific fluorescence can be detected by UV filter glasses for simple, non-intrusive screening of transgene-positive adult zebrafish (**Fig. 5G,H, Supplemental Movie 1**). These data establish *exorh*-based transgenesis markers as a viable alternative to existing transgenesis markers in zebrafish.

A key application benefiting from transgene markers *in cis* is testing of gene-regulatory elements by transient injections and subsequent generation of stable transgenic lines (Felker and Mosimann, 2016; Fisher et al., 2006a; Fisher et al., 2006b; Kague et al., 2010). To facilitate the use of *exorh* reporter-based backbones for regulatory element testing beyond Multisite Gateway, we introduced the 2.4 kb upstream regulatory region of zebrafish *desmin a* (*desma*) together with *beta-globin*-coupled mApple and mCerulean into Tol2 backbones carrying 1) *exorh:EGFP* and 2) *cryaa:Venus*, respectively (**Fig. 6A**). These vectors enable traditional cloning for replacement of the *desma* regulatory sequences (flanked 5’ by *EcoRI, EcoRV* and *NheI*, 3’ by *SalI* and *EcoRI* sites) with other regulatory elements of interest in vectors that include a minimal promoter and a transgenesis marker *in cis*. Upon transient injections into wildtype zebrafish embryos, we observed robust somatic muscle expression of *desma:minprom-mApple* (**Fig. 6B**). Taken together, these experiments establish *exorh:fluorophore*-carrying Tol2 backbones as a new tool for marking transgenes with different cargo.

**Figure 6:**
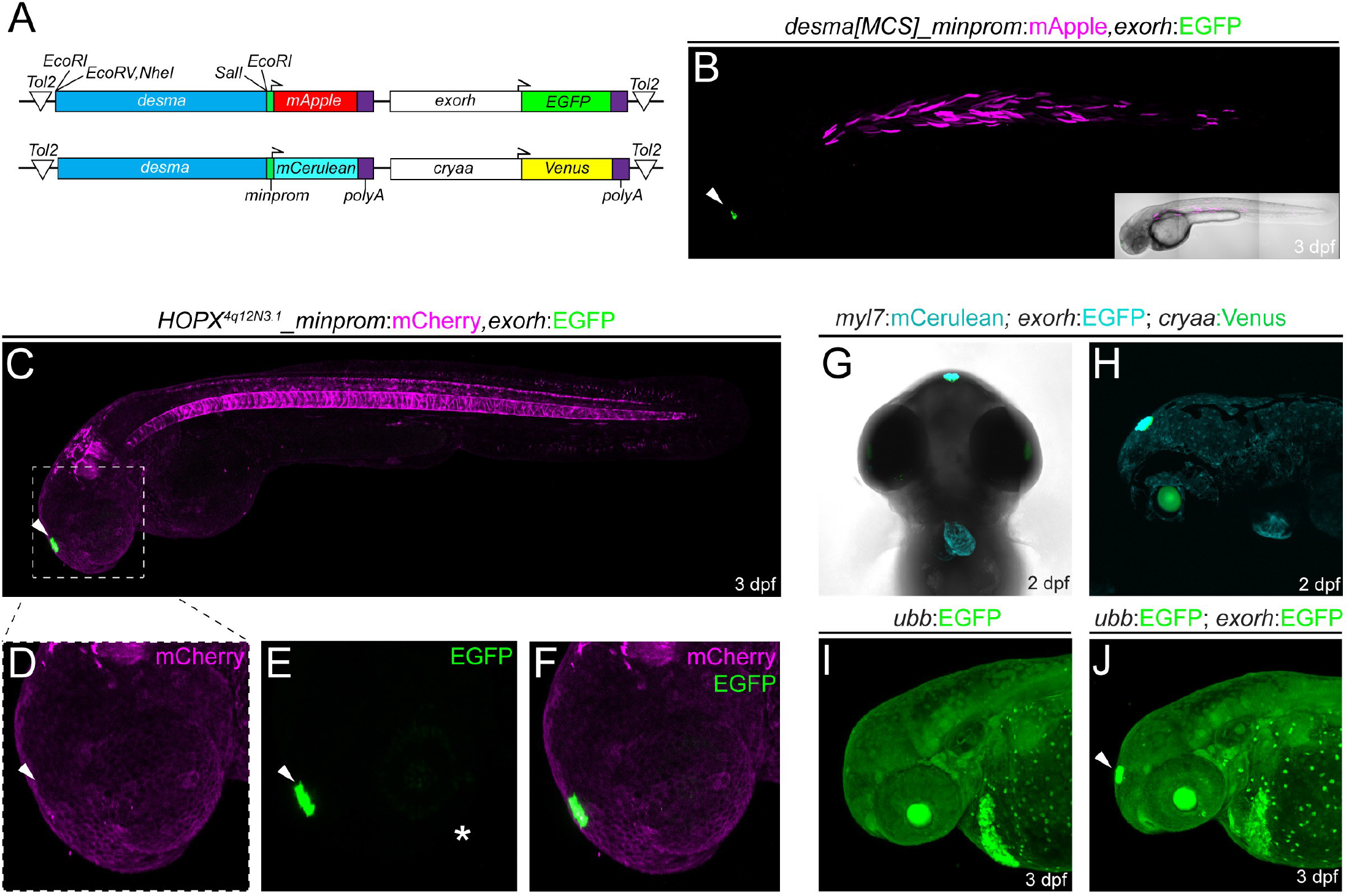
Application examples of an *exorh*-based, pineal gland-specific transgenesis reporter. (**A,B**) Injection-based, transient testing and stable transgenesis of regulatory elements with Tol2 vectors that harbor a transgenesis marker *in cis* for quality control. The *desmin a* (*desma*) upstream region paired with the mouse *beta-globin* minimal promoter driving mApple with *exorh:EGFP*, or driving mCerulean with *cryaa:Venus* (eye lens transgenesis marker) in Tol2 backbones (**A**). In both vectors, the *desma* regulatory region is flanked by restriction sites as simple Multiple Cloning Site (MCS), enabling restriction enzyme-based replacement of *desma* with a regulatory element of interest (**A**). Injection shows muscle-selective *desma* activity (mCherry) and pineal-specific transgenesis reporter expression (EGFP) (**B**). See Methods for details. (**C**-**F**) Regulatory element testing with *exorh:EGFP* as transgenesis marker *in cis*. Stable transgene integration of *HOPX*^*4Q12N3*.*1*^*:minprom-mCherry,exorh:EGFP* at 3 dpf shows mCherry expression in the notochord and rhombomeres, while EGFP expression is confined to the pineal gland (**C**, arrowhead). The transgenes do not cross-activate mCherry (**D,F**) or EGFP (**E,F**) (inserts, white arrowheads) despite being encoded *in cis* and in close proximity, suggesting the *exorh* regulatory region is inert. The white asterisk (**E**) indicates faint, patchy EGFP signal in the eye. (**G,H**) Combinatorial use of transgenesis markers. Ventral (**G**) and lateral (**H**) views of a stable *myl7:mCerulean,exorh:EGFP* transgenic crossed to *cryaa:Venus and* imaged at 2 dpf; note the distinct expression patterns of the secondary transgenic markers labeling the heart (*myl7*), pineal gland (*exorh*), and eye lens (*cryaa*), respectively. (**I,J**) *exorh:EGFP* fluorescence is discernible in ubiquitous green-fluorescent transgenics. Ubiquitous *ubb:EGFP* (*ubi:Switch*) expression (**I**) still allows detection of *exorh:EGFP* expression (white arrowhead) (**J**).

Enhancers can act at a distance on different promoters to activate transcription of target genes (Schaffner, 2015). This distant activation potential is detrimental when combining regulatory elements *in cis* on a confined transgenesis vector, such as a transgenesis reporter together with an enhancer-driven transgene. To assess the robustness and utility of the new *exorh*-based transgenesis backbones, we tested each *exorh:fluorophore* combination in assembled MultiSite Gateway constructs to determine if the *exorh* promoter cross-reacts with any reporters cloned *in cis*, in particular minimal promoter-containing reporters. In stable transgene integrations of *HOPX_4Q12N3*.*1:minprom-mCherry, exorh:EGFP*, we observed faithful mCherry activity in the notochord and rhombomeres and EGFP expression in the pineal gland, as anticipated (**Fig. 6C,E,F**); however, we did not observe any mCherry signal in the pineal (**Fig. 6D**) or notochord expression of EGFP (**Fig. 6C**). While not comprehensive, this basic test indicates that both regulatory elements in our observed transgenic insertion were inert to their reciprocal activities.

Crossing stable transgenics of *myl7:mCerulean,exorh:EGFP* with the *hsp70l:Switch* line that contains the *cryaa:Venus* lens marker in the backbone (Felker et al., 2018; Hesselson et al., 2009) further documented how the pineal gland marker can be used in combination with commonly applied transgene reporters (**Fig. 6G,H**). *exorh*-based fluorophore reporters such as *exorh:EGFP* are reproducibly strong and bright, enabling their use in ubiquitous fluorescent backgrounds, as documented with easily detectable *exorh:EGFP* expression and ubiquitous EGFP (**Fig. 6I,J**). Together, these observations establish *exorh:fluorophore* Tol2 backbones as new transgenesis vectors for a multitude of applications.

### Zebrafish *exorh:fluorophore* reporters are functional in *Astyanax*

Establishing a new model organism greatly benefits from access to cross-compatible transgenesis reagents. The Mexican cavefish (*Astyanax mexicanus*) is an increasingly applied model system to study evolutionary selection and metabolism (McGaugh et al., 2020; Rohner, 2018). While transgenesis has been successfully introduced in the system including the use of originally zebrafish-derived regulatory sequences (Elipot et al., 2014; Lloyd et al., 2022; Stahl et al., 2019), a more widespread use has been hampered by the lack of functional transgenesis markers: *myl7*-based cardiac reporters seem not functional when tested in *Astyanax* (Elipot et al., 2014) and the diverse *crystallin* gene-based eye lens transgenesis markers cannot be used due to the early eye degeneration phenotype in cavefish. We therefore sought to test the functionality of *exorh*-based reporters in the model.

Injection of a Tol2 expression construct carrying *exorh:EGFP* (**Fig. 7A**) into one-cell stage embryos of different populations of *Astyanax mexicanus* (derived from surface and Tinaja cave populations) resulted in easily detectable pineal gland expression beginning around hatching stage (1 dpf) in both Tinaja cave and surface morphs (**Fig. 7B-E**). At 5 dpf, we continued to observe robust expression in both morphs (**Fig 7C,E**), detectable from different imaging angles (lateral-dorsal-ventral) (**Fig. 7F-H**). Pineal expression of *exorh*-driven fluorophores continued throughout the larval stages of surface and cave morphs into adulthood (**Fig. 7I,J**). Tol2-based destination vectors engineered with the *exorh* promoter driving a fluorescent marker (tested here as mGreenLantern and mCherry) (Campbell et al., 2020) also recapitulated the pineal gland expression pattern in transient injections (**Fig. 7K-O**) and can be applied to generate transgenesis constructs using MultiSite Gateway. Of note, in our transient injections we observed weak, mosaic expression in retinal cells of surface morph animals (**Fig. 7M**). Taken together, these observations document the activity of zebrafish *exorh* promoter-based reporters in *Astyanax* as a new transgenesis marker as supported by the developmental retention of the pineal gland in cave morphs.

**Figure 7:**
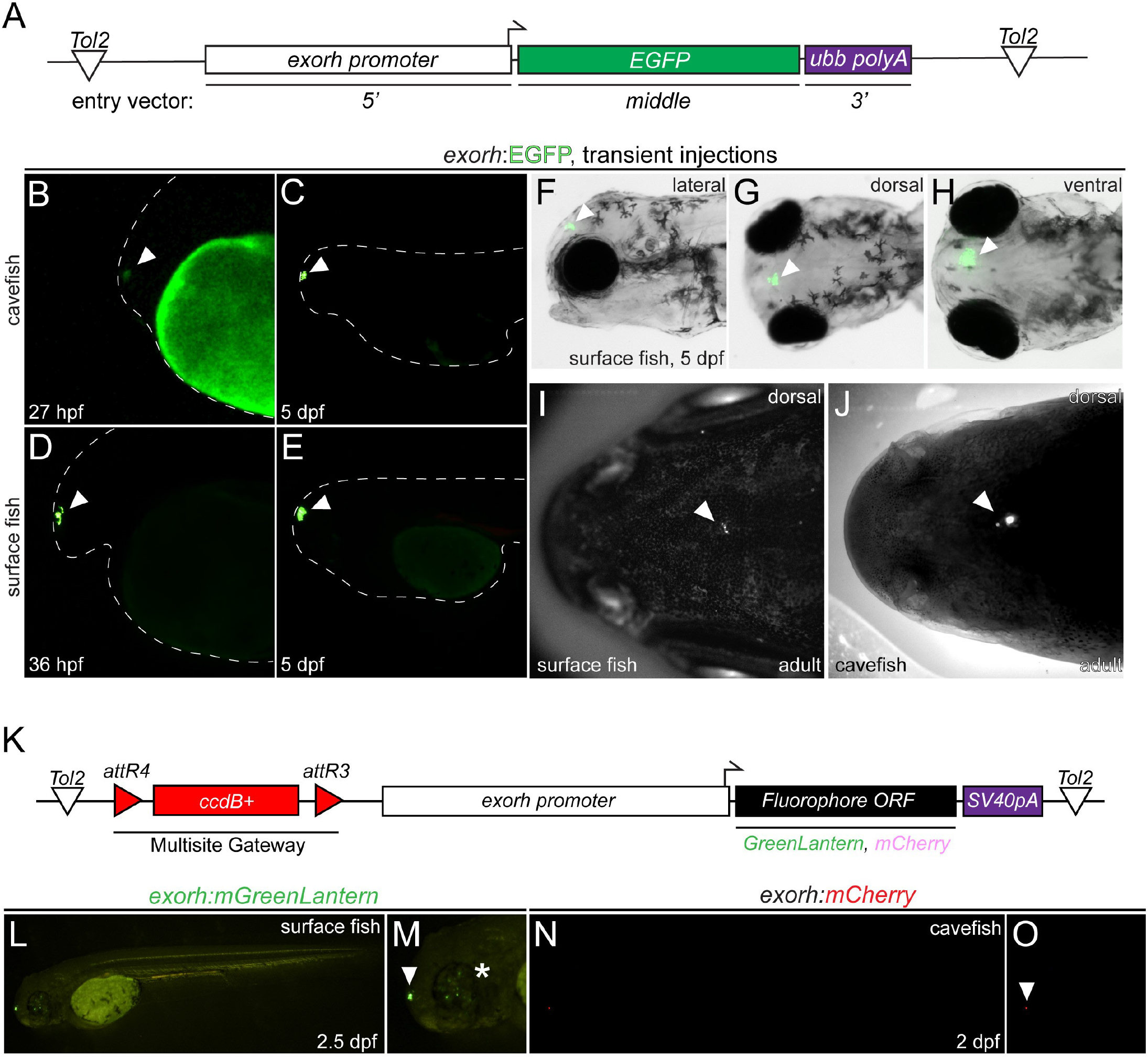
The *exorh*-based transgenesis marker functions in Mexican cavefish (*Astyanax mexicanus*). (**A**) Schematic of the Tol2 expression vector using the zebrafish *exorh* promoter driving the expression of EGFP; see also Figure 5. (**B-E**) Transient injection of *exorh:EGFP* shows reporter activity in the pineal gland of both Tinaja cavefish at 27 hpf (**B**) and 5 dpf (**C**), as well as in surface fish at 36 hpf (**D**) and 5 dpf (**E**). Pineal expression is indicated with an arrowhead and embryo morphology is traced with a dashed line. (**F-H**) Lateral, dorsal, and ventral views at 5 dpf showing specific, easily detectable EGFP reporter activity in the pineal gland (arrowheads). (**I,J**) Grayscale images of live *exorh:EGFP*-injected animals, detection of EGFP (white dots) in surface and cavefish adults (arrowheads). (**K**) Schematic of Tol2-based transgenesis backbones *pDEST-exorh:mGreenLantern* and *pDEST-exorh:mCherry* including the *attR4* and *attR3*-flanked *ccdB* cassette and the 1055 bp zebrafish *exorh* promoter driving *mGreenLantern* or *mCherry* and for Multisite Gateway cloning. (**L,M**) Injection of the *exorh:mGreenLantern*-containing Tol2 Destination vector shows pineal gland expression (arrowhead) as well as patchy retinal expression in developing surface fish (asterisk). (**N,O**) Injection of the *exorh:mCherry*-expressing Tol2 destination vector, showing pineal gland expression in developing cavefish (arrowheads).

## Discussion

Tol2-based transgenesis and the wide accessibility of the required basic tools has led to a revolution of transgenic reporter and effector strains in the zebrafish field (Kawakami, 2005; Kawakami, 2007). The industriousness and collegiality of the zebrafish community has facilitated expansion beyond the basic tools as well as extensive sharing and distribution. While by no means comprehensive, our data and reagents presented here augment, diversify, and increase the functionality of the transgenesis toolkit for zebrafish and other systems. All plasmid sequences and maps are available with this manuscript (**Supplemental Data File 1**) and all reagents are in the process of submission for distribution by Addgene.

The widespread use of green and red fluorophores, in particular EGFP, mCherry, and dsRED2, increasingly hinders the combinatorial use of reporter transgenes. We provide a set of middle and 3’ entry vectors containing mCerulean, a blue-fluorescent avGFP derivative, which provides a complementary fluorophore to common green and red fluorophores. mCerulean is detectable with standard GFP filters on fluorescent dissecting scopes as routinely used in the field, yet does benefit from using a dedicated blue filter set and matching excitation light source (**Fig. 2**). As an alternative to mCherry and dsRED2, mApple provides a complementary red fluorophore that is detectable with standard RFP filters (**Fig. 3**); we also found that antibodies against dsRED detect mApple to stain for its expression (data not shown). Combined with its better theoretical quantum yield and fast folding mechanism, mApple constructs provide a versatile alternative to mCherry and dsRED derivatives.

Regulatory element testing is a powerful approach in zebrafish; the discovery and testing of regulatory elements from different species can reveal fundamental insights into upstream regulation of gene expression as well as into the evolutionary and developmental ontogeny of cell types (Fisher et al., 2006a; Fisher et al., 2006b). As enhancer sequences need to act on a promoter region to support assembly of productive RNA Pol II complexes, transgene constructs for enhancer testing require a sensitive, yet position effect-inert minimal promoter region that is also ideally short to keep assembled vector sizes down to facilitate cloning and delivery (Felker and Mosimann, 2016). Several minimal promoter regions have been successfully deployed in zebrafish, including sequences from divergent species. The *Xenopus Ef1a* minimal promoter has been repeatedly used for basic transgenes, yet remains sensitive to silencing (Amsterdam et al., 1995; Tamplin et al., 2011). The zebrafish *gata2a* promoter region has been successfully applied in a variety of settings including enhancer trap vectors and reporters (Bessa et al., 2009; Meng et al., 1997); however, its comparatively large size (over 2 kb) challenges larger transgene vectors. Previous work has provided solid evidence for the potency of the mouse *beta-globin* minimal promoter to test various regulator elements including notochord and hematopoietic enhancers (Tamplin et al., 2011; Tamplin et al., 2015; Woolfe et al., 2004). We have previously incorporated this module in testing enhancers active in the lateral plate mesoderm and neural crest across species (Kaufman et al., 2016; Parker et al., 2019; Prummel et al., 2019). Here, we provide vectors pairing the mouse *beta-globin* minimal promoter with EGFP, mCherry, mApple, and mCerulean for versatile gene-regulatory element testing (**Fig. 1**). While required for enhancers, upstream regions that contain the transcription start of a gene of interest might not require an additional minimal promoter as the provided sequences might suffice to assemble a functional RNA Pol II pre-initiation complex. Incorporation of the mouse *beta-globin* minimal promoter is advisable when testing regulatory elements from divergent species or when establishing basic regulatory element activity (Bergman et al., 2022). Notably, we also present here the *pAP02 p3E MCS* 3’ entry vector that enables cloning of enhancer elements downstream of the reporter cassette, resulting in the enhancer getting incorporated into the resulting mRNA before being terminated and polyadenylated. This vector design will allow the potential application of methods to discover active regulatory elements by sequencing, such as successfully performed in cell culture systems as STARR-seq (Arnold et al., 2013; Neumayr et al., 2019).

Our collection of 2A peptide fusions with different fluorophores and localization tags is suitable to generate bi-protein constructs with minimal C-terminal tagging (**Fig. 4**). Previous attempts to broadly apply IRES sequences for bi-cistronic transgenes and fluorescent labeling have widely been unsuccessful for as-of-yet unclear reasons (Kwan et al., 2007). Viral p2A or similar peptides that break the emerging polypeptide chain during translation have been widely applied to express several genes of interests at equal protein levels in diverse cell culture systems, such as for cardiac reprogramming (Muraoka and Ieda, 2015; Song et al., 2012; Zhang et al., 2019). The direct fluorescent labeling of cells expressing a gene of interest for modifier studies, functional tests, and disease modeling provides a potent phenotype readout and imaging tool (**Fig. 4**). Despite the simple cloning that requires omission of the stop codon in the N-terminal ORF of interest, cloned in-frame with the following *attB2*Gateway repeat to connect with a 3’ p2A-fluorophore, the polypeptide breakage leaves a residual p2A sequence as well as the encoded Gateway repeat at the C-terminal end of the ORF of interest. Mechanistic tests are therefore warranted to ensure proper functionality of this tagging method in individual transgenes.

Versatile, easy-to-screen transgenesis markers greatly augment any model organism tool kit by enabling rigorous quality control and transgene identification. A frequently used transgenesis marker in zebrafish is based on the *myl7* promoter that drives fluorescence in cardiomyocytes starting from late somitogenesis stages, but routinely from 24 hpf onward (Huang et al., 2003). However, the heart is obscured by skin and pigment in the growing and adult zebrafish and the marker cannot be easily adapted for transgenes associated with cardiac biology. *crystallin* gene-based reporters drive fluorescence in the eye lens, providing an elegant and simple reporter property for screening in adults using UV filter goggles and handheld flashlights. We here add two *crybb1*-based reporters driving mKate2 and TagBFP in Tol2-based Multisite Backbones to the roster (**Fig. 2A, Fig. 4G,H**). A major drawback from *crystallin*-based reporters is their comparatively late onset of detectable fluorophore expression from around 36 hpf or later (Davidson et al., 2003; Kurita et al., 2003), which can render sorting of transgene-positive embryos challenging. Our *exorh*-based reporters documented here (**Fig. 5,6**) circumvent this issue and provide reporters that are reproducibly active from 22ss and stay active throughout adulthood (Asaoka et al., 2002); *exorh*-marked backbones are thereby screenable while embryos remain in their chorions as well as with UV filter goggles in adults (**Supplemental Movie 1**). The confined, small area of fluorescence dorsal and anterior in the head does not obscure major organs during early development and should still enable imaging of significant brain regions in fluorescent transgenic reporters or stained embryos. *exorh:fluorophore* cassettes are less than 2 kb in size, and we here documented several successful germline transgenes using such Tol2 backbones (**Fig. 5,6**). Notably, *exorh*-marked transgenes are easily combined with other transgenes with *myl7-, crystallin*-, or even ubiquitously expressed reporters (**Fig. 6E-H**). We here generated three distinct *exorh:fluorophore* combinations to facilitate planning and execution of such combinatorial experiments, and we provide versions compatible with restriction enzyme-based cloning and MultiSite Gateway (**Fig. 5A**). The provided backbones that harbor *desma*-driven fluorophores enable restriction site-based exchange of the *desma* regulatory region to test different regulatory elements (**Fig. 6A**). In addition, *desma*-driven reporters express well in somatic muscle and are simple to detect (**Fig. 6B**), providing potent reagents to troubleshoot injections, to test *Tol2* mRNA batches, or to train new researchers in transgenesis basics.

Use of our vector tools is not confined to zebrafish, as many can be applied in other models and Gateway-compatible cloning reactions. We have previously used mouse *beta-globin*-containing enhancer constructs used for zebrafish transgenes without modifications for electroporation in chick (Prummel et al., 2019), underscoring the utility of the mouse *beta-globin* minimal promoter for enhancer testing. Relevant to other aquatic models, *exorh*-based reporter utility expands to *Astyanax*, an emerging model for the study of evolutionary adaptation (**Fig. 7**) (Elipot et al., 2014; McGaugh et al., 2020; Rohner, 2018). While the cave forms of *Astyanax* have lost their eyes, they do form functional pinealocytes and a pineal gland, although their impact on behavior and circadian rhythms remains to be fully elucidated (Lloyd et al., 2022; Mack et al., 2021; Omura, 1975; Yoshizawa and Jeffery, 2008). *exorh*-based reporter transgenes have the potential to facilitate transgene quality control, visual genotyping, and combinatorial transgene experiments such as Cre/*lox* approaches in the model. As the pineal gland remains visible on the skull of fishes and amphibians, *exorh*-based reporters have the potential to support transgenesis experiments in a wide variety of aquatic models.

## Methods

### Zebrafish Husbandry and Procedures

Animal care and procedures were carried out in accordance with the veterinary office of the University of Zürich, Zürich, Switzerland; the IACUC of the University of Colorado School of Medicine (protocol #00979), Aurora, Colorado, USA; and the IACUC of the University of Utah (protocol #21-01007), Salt Lake City, Utah, USA. All zebrafish embryos were incubated at 28.5 °C in E3 medium unless indicated otherwise. Staging was performed as described by Kimmel et al. (Kimmel et al., 1995).

### Cavefish Husbandry and Procedures

Astyanax husbandry and care was conducted identical to reported procedures (Xiong et al., 2022) and as approved by the Institutional Animal Care and Use Committee (IACUC) of the Stowers Institute for Medical Research on protocol 2021-122. All methods described here are approved on protocol 2021-129. Housing conditions meet federal regulations and are accredited by AAALAC International.

### Plasmid Construction

All plasmids generated for this study are in the process of being deposited, along with maps and sequence files, with AddGene for non-profit distribution. First-time users of Tol2 technology are asked to connect with Dr. Koichi Kawakami to set up a material transfer agreement (MTA). Plasmid sequences and maps are available online accompanying this manuscript (**Supplemental Data File 1**). The full list of deposited vectors is:

| Plasmid Name | Plasmid Type | Resistance | Source Lab |
| --- | --- | --- | --- |
| *pCK033 p5E desma MCS* | *p5E* | Kanamycin | Mosimann |
| *pCK011 p5E exorh* | *p5E* | Kanamycin | Mosimann |
| *pCK072 p5E 147bp exorh:EGFP* | *p5E* | Kanamycin | Mosimann |
| *pCK074 p5E exorh:EGFP* | *p5E* | Kanamycin | Mosimann |
| *pCK051 p5E exorh:mCerulean* | *p5E* | Kanamycin | Mosimann |
| *pCK052 p5E exorh:mCherry* | *p5E* | Kanamycin | Mosimann |
| *pCK006 pME minprom_mCherry no stop* | *pME* | Kanamycin | Mosimann |
| *pKD001 pME minprom_EGFP* | *pME* | Kanamycin | Mosimann |
| *pCK002 pME mCerulean* | *pME* | Kanamycin | Burger |
| *769 pME mCerulean-CAAX* | *pME* | Kanamycin | Kwan |
| *765 pME nls-mCerulean* | *pME* | Kanamycin | Kwan |
| *pSN001 pME minprom_mCerulean* | *pME* | Kanamycin | Burger |
| *763 pME mApple* | *pME* | Kanamycin | Kwan |
| *768 pME mApple-CAAX* | *pME* | Kanamycin | Kwan |
| *764 pME nls-mApple* | *pME* | Kanamycin | Kwan |
| *pCK068 pME minprom_mApple* | *pME* | Kanamycin | Mosimann |
| *pME minprom_KalTa4* | *pME* | Kanamycin | Burger |
| *pCK007 pME KalTa4* | *pME* | Kanamycin | Burger |
| *pME minprom_creERT2* | *pME* | Kanamycin | Mosimann |
| *457 p3E-2A-EGFPpA* | *p3E* | Kanamycin | Kwan |
| *458 p3E-2A-EGFP-CAAXpA* | *p3E* | Kanamycin | Kwan |
| *459 p3E-2A-nlsEGFPpA* | *p3E* | Kanamycin | Kwan |
| *460 p3E-2A-mCherry pA* | *p3E* | Kanamycin | Kwan |
| *612 p3E-2A-mCherryCAAXpA* | *p3E* | Kanamycin | Kwan |
| *766 p3E-2A-nlsmCherry pA* | *p3E* | Kanamycin | Kwan |
| 791 *p3E-2A-pCS2MCSpA v2* | *p3E* | Kanamycin | Kwan |
| *pAF020 p3E-2A-mCerulean* | *p3E* | Kanamycin | Mosimann |
| *pAF018 p3E-2A-mCerulean-CAAX* | *p3E* | Kanamycin | Mosimann |
| *pAP02 p3E MCS* | *p3E* | Kanamycin | Mosimann |
| *p3E ubb polyA* | *p3E* | Kanamycin | Burger |
| *pCK004 ubi:mCerulean_SV40 (#302)* | Expression construct | Ampicillin | Burger |
| *pCK005 ubi:mCerulean_ubb-polyA* | Expression construct | Ampicillin | Burger |
| *pCK029 exorh:EGFP* | Expression construct | Ampicillin | Mosimann |
| *pCK036 desmaMCS:minCerulean,cryaa:Venus* | Expression construct | Ampicillin | Mosimann |
| *pCK069 desmaMCS:minprom mApple,exorh:EGFP* | Expression construct | Ampicillin | Mosimann |
| *pCK017 pDEST exorh:EGFP* | *pDEST* | Ampicillin, Chloramphenicol, ccdB | Mosimann |
| *pCK053 pDEST exorh:mCherry-* | *pDEST* | Ampicillin, Chloramphenicol, ccdB | Mosimann |
| *pCK054 pDEST exorh:mCerulean* | *pDEST* | Ampicillin, Chloramphenicol, ccdB | Mosimann |
| *pCB59 pDEST crybb1:mKate2* | *pDEST* | Ampicillin, Chloramphenicol, ccdB | Affolter |
| *pCB24 pDEST crybb1:TagBFP* | *pDEST* | Ampicillin, Chloramphenicol, ccdB | Affolter |
| *pDEST exorh:emGreenLantern* | *pDEST* | Ampicillin, Chloramphenicol, ccdB | Rohner |
| *pDEST exorh:mCherry* | *pDEST* | Ampicillin, Chloramphenicol, ccdB | Rohner |

### Entry Clones

All entry clones were verified by colony PCR and sequencing with *M13 fw*: *5’-GTAAAACGACGGCCAG-3’* and *M13 rv*: *5’-CAGGAAACAGCTATGAC-3’*.

### p5E

For *p5E* entry clones, all PCR products were amplified using Roche Expand High Fidelity PCR kit (Sigma 11732641001), which contains a 3’-adenine overhang-depositing Taq. PCR products were purified using the QIAquick PCR Purification Kit (Qiagen 28104) or the QIAquick Gel Extraction Kit (Qiagen 28706) using ddH2O elution as last step. PCR products were then TOPO-cloned into MultiSite Gateway entry vectors with the *pENTR5’-TOPO TA* Cloning Kit (Invitrogen K59120): 4 μL of purified PCR product was immediately combined with 1 μL *pENTR5’-TOPO TA* and 1 μL of salt solution and incubated at room temperature for a minimum of 2 hours to overnight to ligate. 3 μL of the TOPO reactions were then transformed with One Shot™ TOP10 Chemically Competent *E. coli* (Invitrogen C404010) and plated on Kanamycin-infused agar plates. Half reactions can also be used. Primer sequences for *pE5’* enhancer/promoter fragments are as follows:

*pCK079 p5E MmBMP4ECR2*: *oBuS014 Mm BMP4 ECR2 fw*:

*5’-GGGGATGAAAGTAGCATCCTG-3’ oBuS015 Mm BMP4 ECR2 rv*:

*5’-TTCCACTTTGCTTCCCAAACTGG-3’ pAB019 pE5 Hs HOPX_4q12N3*.*1*:

*oAB019 Hs HOPX fw*: *5’-GTGTTGGGTTAGTTTGAGC-3’*

*oAB019 Hs HOPX rv: 5’-GTTCTGTTGGGGATATGTCC-3’*

*pCK033 p5E desma MCS*:

*oCK024 Dr desma MCS fw*:

*5’-gatatcACTGATgctagcTCCTTGAGGCACTTTCGG-3’*

*oCK025 Dr desma MCS rv*:

*5’-gtcgacTACGCTGTGTGAATGCTGG-3’*

*pME, pENTR/D, mouse beta-globin minimal promoter, and fluorophore ORFs*

For middle entry vectors, blunt-end PCR products were generated using Phusion High Fidelity DNA Polymerase (NEB M0530S) and forward primers were designed including *5’-CACC-3’* on the *5’* end for directional cloning and then ligated into MultiSite Gateway™-compatible *pME* entry vectors using *pENTR/D-TOPO* (Invitrogen K240020). 4 μL of purified PCR product was combined with 1 μL *pENTR/D-TOPO* and 1 μL of salt solution and incubated at room temperature for a minimum of 2 hours to overnight to ligate. 3 μL of the TOPO reactions were then transformed with One Shot TOP10 Chemically Competent *E. coli* and plated Kanamycin-containing agarose plates. Half reactions can also be used.

The mouse *beta-globin* (*Hbb-bt*) minimal promoter sequence spans 129 base pairs: *5’-CCAATCTGCTCAGAGAGGACAGAGTGGGCAGGAGCCAGCATTGGGTATATAAAGCTGAGCAGGGTCAGTTGCT TC<u>TTACGTTTGCTTCTGAGTCTGTTGTGTTGACTTGCA ACCTCAGAAACAGACATC</u>-3’*, as annotated in the reference genome *NC_000073*.*7:103463077-103463205 Mus musculus* strain *C57BL/6J* chromosome 7, *GRCm39*; underlined sequence stretch represents the 5’ UTR.

The mCerulean ORF sequence can be amplified using *mCerulean fw*: *5’-ATGGTGAGCAAGGGCGAGGAGC-3’* and *mCerulean rv*: *5’-CAGCTCGTCCATGCCGAGAGTG-3’*. The mApple ORF sequence can be amplified using *mApple fw*: *5’-ATGGTGAGCAAGGGCGAGG-3’* and *mApple rv*: *5’-CTTGTACAGCTCGTCCATGC-3’*.

### p3E

*pAP02_p3E-MCS* was generated by amplifying the multiple cloning site from *pBluescript KSII(+)* with the primers *AP13fw*:

*5’-GGGGACAGCTTTCTTGTACAAAGTGGGCGGCCGCTC TAGAA-3’* and *AP14rv*:

*5’-GGGGACAACTTTGTATAATAAAGTTGGGTACCGGGC CCCCC-3’* and inserted into *pDONRP2R-P3* via BP reaction.

*P3E_ubb-polyA* was cloned by amplifying a 516 bp fragment of the zebrafish *ubiquitinB* (*ubb, ubi*) locus using the primers *ubb polyA fw*: *5’-TAGAACCGACAGTCTTAGGGATGG-3’* and *ubb polyA rv*: *5’-GAATTCATTGCCATCAAGTGTTAGC-3’* with Phusion High Fidelity DNA Polymerase (NEB M0530S), subcloned into the *Zero Blunt TOPO PCR* Cloning vector (Invitrogen K283020), digested out with BglII and ligated into the single cutter BamHI site in *pAP02_p3E-MCS*.

Of note, frequent users of both the *#302 p3E-polyA* from the original Tol2kit (Kwan et al., 2007) and *p3E_ubb-polyA* introduced here have experienced decreased Multisite Gateway reaction efficiencies after multiple freeze-thaw cycles of these *p3E* vectors. For optimal colony numbers and reaction efficiency, we recommend freshly preparing minipreps, facilitated by frequent retransformations of the vector and storage of the plasmid in small aliquots. The reason for this reduction in cloning efficiency with prolonged storage of these p3E vectors is currently unknown, but could possibly be due to structural factors such as secondary structure formation detrimental to Multisite Gateway assembly or bacteria-based propagation. 2A constructs were built via a two-step process. First, the 2A peptide-encoding sequence was added to the N-terminus of fluorescence protein ORFs using PCR and inclusion of the 2A peptide sequence in the 5’ upstream primer. The fusion product was conventionally cloned into *pCS2FA* using the *FseI* and *AscI* sites (enzymes from NEB). This placed the fusion product upstream of the *SV40 polyA* signal sequence. Ligation reactions were transformed using Subcloning Efficiency *DH5α* chemically competent *E. coli* (Invitrogen/ThermoFisher) and plated onto Ampicillin- or Carbenicillin-containing agar plates. Cloning was confirmed by restriction digest and subsequent sequencing. This *2A-FP-polyA* cassette was then cloned into the 3’ entry vector using common *att* PCR primers (*2A-attB2F* and *CS2pA-attB3R*). For *p3E-2A-CS2MCS-pA, pCS2P+* was used as a template for a PCR reaction in which the 2A peptide-encoding sequence was included in the *att2F* forward primer. Care was taken to ensure that all primers used “Gateway reading frame”, in which bases listed in the manual as “*N*” in the primer were included as “*G*”. This ensures maintenance of reading frame between the middle entry clone (which must not include a stop codon) and the 3’ clone containing the 2A peptide sequence and reporter. PCR products were gel-purified (Qiagen Gel Extraction Kit) and quantified. For the recombination, equimolar amounts (usually 50 fmoles each) of purified PCR product were combined with *pDONR-P2R-P3* donor vector and Gateway BP Clonase II Enzyme Mix (Invitrogen/ThermoFisher). Reactions were incubated at room temperature for several hours and transformed using Subcloning Efficiency *DH5α* chemically competent *E. coli* (Invitrogen/ThermoFisher) and plated onto Kanamycin-containing agar plates. Cloning was confirmed by restriction digest and subsequent sequencing.

For *2A-FP* cloning into *pCS2FA* (restriction sites in bold, sequence complementary to template <u>underlined</u>): *2AEGFP5Fse*: *5’-gaac****GGCCGGCC*** *GGATCCGGAGCCACGAACTTCTCTCTGTTAAAGCAAGCAGGAGACGTGGAAGAAAACCCCGGTCCT <u>ATGGTGAGCAAGGGCGAGGAGCTG</u>-3’*

*2AmCherry5Fse*: *5’-gaac****GGCCGGCC*** *GGATCCGGAGCCACGAACTTCTCTCTGTTAAAGCAAGCAGGAGACGTGGAAGAAAACCCCGGTCCT <u>ATGGTGAGCAAGGGCGAGGAGGAC</u>-3’*

*2Anls5Fse*: *5’-gaac****GGCCGGCC*** *GGATCCGGAGCCACGAACTTCTCTCTGTTAAAGCAAGCAGGAGACGTGGAAGAAAACCCCGGTCCT <u>ATGGCTCCAAAGAAGAAGCGTAAGG</u>-3’*

*EGFP3Asc (rev)*:

*5’- gaac****GGCGCGCC****<u>TCACTATAGGGCTGCAGAATCTAGA GG</u>-3’*

*mCherry3Asc (rev)*:

*5’- gaac****GGCGCGCC****<u>TTACTTGTACAGCTCGTCCATGCCG C</u>-3’*

*CAAX3Asc (rev)*:

*5’- gaac****GGCGCGCC****<u>TCAGGAGAGCACACACTTGCAGCTC</u> -3’*

For *p3E* cloning from *pCS2FA*:

*2A-attB2F*:

*5’- ggggACAGCTTTCTTGTACAAAGTGGGG<u>GGATCCGGA GCCACGAACTTCTCTC</u>-3’*

*2A-CS2PMCS-attB2F*:

*5’-ggggACAGCTTTCTTGTACAAAGTGGGG GGATCCGGAGCCACGAACTTCTCTCTGTTAAAGCAAG CAGGAGACGTGGAAGAAAACCCCGGTCCT AG<u>GGATCCCATCGATTCGAATTCAAGGCCTC</u>-3’*

*CS2pA-attB3R (rev)*: *5’- ggggACAACTTTGTATAATAAAGTTGG<u>AAAAAACCTCCC ACACCTCCCCCTG</u>-3’*

### Expression Constructs

Multisite Gateway recombination reactions were principally performed as described in the Invitrogen Multisite Gateway Manual, with minor modifications. 10 fmol of *pE5’, pME*, and *pE3’* vectors were combined with 20 fmol of destination vector, 1 μL of LR Clonase II Plus Enzyme Mix (Invitrogen 12538120, vortexed 2x for two seconds prior to use), and ddH2O for a final reaction volume of 5 μL. Vector calculations for molarity were performed using the Multisite Gateway Excel spreadsheet (Mosimann, 2022). Reactions were incubated at 25°C overnight and treated the following day with 1 μL 2 μg/μL Proteinase K for 15 minutes at 37°C and 3 μL of the reaction was transformed with One Shot TOP10 Chemically Competent *E. coli* (Invitrogen C404010) and plated on Ampicillin selection plates. T4 ligations into multiple cloning site plasmids were performed by amplifying fragments via PCR with Roche Expand High Fidelity Polymerase (Sigma 11732641001) with primers containing 5’ restriction enzyme sequences for digestion following PCR purification. Digested backbones were treated with shrimp alkaline phosphatase (NEB M0371S) for 2 hours at 37 °C prior to gel extraction. T4 ligations were all performed with a 1:3 pmol ratio of backbone to insert at 16 °C overnight. 7.5 μL of T4 reactions were then transformed with One Shot TOP10 Chemically Competent *E. coli* (Invitrogen C404010) and Ampicillin selection. All assembled gateway reactions were tested by diagnostic digest and confirmed by sequencing using the primers below (Mosimann, 2022): *attB1 fw*: *5’-CAAGTTTGTACAAAAAAGCAGGCT-3’ attB1 rv*: *5’-AGCCTGCTTTTTTGTACAAACTTG-3’ attB2 fw*: *5’-ACCCAGCTTTCTTGTACAAAGTGG-3’ attB4 fw*: *5’-CAACTTTGTATAGAAAAGTTG-3’*

### Zebrafish exorh promoter element

The *exorh promoter:EGFP* was amplified from the *pCR2*.*1 TOPO* vector containing a 1055 bp upstream region and short coding sequence of the zebrafish *exorh* gene driving *EGFP* (a kind gift from Dr. Yoshitaka Fukuda and Dr. Daisuke Kojima) (Asaoka et al., 2002) with 5’ *Asp718l* (*Acc651*) restriction site containing primers. Of note, exorh lies on the edge of a genomic contig in the sequenced zebrafish genome, rendering the promoter isolation challenging without the pioneering work by Asaoka et al.. The *Asp718l*-flanked *exorh:EGFP* fragment was subcloned into *pENTR5’-TOPO* as stated in the Entry Clone section above to generate *pCK074 pE5’ exorh:EGFP*. The shorter 147 bp *exorh* promoter fragment driving EGFP was amplified from the same vector (Asaoka et al., 2002) and subcloned in the same way to generate *pCK072 pE5’ 147bp exorh:EGFP*. The *exorh* promoter alone was also subcloned into *pENTR5’-TOPO* to generate *pCK011 pE5’ exorh*. The vectors containing the *exorh* promoter driving mCerulean and mCherry ORFs were cloned using PCR fusion of *Asp718l*-flanked fragments and subcloned into *pENTR5’-TOPO* to generate *pCK051 pE5’ exorh:mCerulean* and *pCK052 pE5’ exorh:mCherry*, respectively. Of note, the *exorh* promoter sequence contains a *GT*-rich repeat region, resulting in challenging PCR amplification. Primer sequences are stated below:

*pCK011 p5E exorh:*

*oCK001 exorh fw 5’- GCTCAGCTGGCAGTACTACC-3’*

*oCK009 exorh rv 5’-GATGGAGAAGTGGACGATCG-3’*

*pCK072 p5E 147bp Exorh:EGFP*: *oCK018 ASP7181-147bpexorh:EGFP fw*

*5’- agatctggtaccGCGAGCGCTGCTGTGTCTCC-3’*

*oCK017 ASP7181-exorh:EGFP Rv*

*5’-agatctggtaccTTACTTGTACAGCTCGTCC-3’*

p*CK074 p5E Exorh:EGFP*:

*oCK016 ASP7181-exorh:EGFP fw*:

*5’-agatctggtaccGCTCAGCTGGCAGTACTACC-3’ oCK017 ASP7181-exorh:EGFP rv*:

*5’-agatctggtaccTTACTTGTACAGCTCGTCC-3’*

*pCK051 p5E Exorh:mCerulean*:

*oCK016 ASP7181-exorh:EGFP fw*:

*5’-agatctggtaccGCTCAGCTGGCAGTACTACC-3’ oCK054 mcerulean-STOP-Asp7181 rv*:

*5’-GGTACCtaCAGCTCGTCCATGCCGAGAG-3’*

*pCK052 p5E Exorh:mCherry*:

*oCK016 ASP7181-exorh:EGFP fw*:

*5’-agatctggtaccGCTCAGCTGGCAGTACTACC-3’ oCK056 mcherry-STOP-Asp7181 rv*:

*5’-GGTACCtaCTTGTACAGCTCGTCCATGC-3’*

### Destination Vectors

To generate destination vectors containing the *exorh* promoter driving various fluorophores as secondary transgenic markers, digested *Asp718l-exorh:fluorophore-Asp718l* fragments were T4-ligated as described above into the Asp718l-digested and dephosphorylated *pCM326 pDEST_cryaa:Venus* (Mosimann et al., 2015) following gel extraction of the digested backbone minus the *cryaa:Venus* cassette to generate *pCK017 pDEST exorh:EGFP, pCK053 pDEST exorh:mCherry*, and *pCK054 pDEST exorh:mCerulean*. Clones were verified by test digest and full plasmid sequencing. All destination vectors were transformed with One Shot *ccdB* Survival 2 T1R Chemically Competent Cells (Invitrogen A10460) with both Ampicillin and Chloramphenicol selection.

*pCB59_crybb1:mKate2* was amplified from *pDEST-Tol2_Fli1ep-mCherry-GM130* (a kind gift from Dr. Holger Gerhard) (Franco et al., 2015) including the zebrafish *crybb1* promoter, parts of the minimal *CMV* promoter, and the mKate2 ORF. The PCR product was cloned into the *Tol2*-based Multisite Gateway backbone *pDestTol2CG2* (Tol2kit *#395*) (Kwan et al., 2007) via the *BglII* restriction site. Of note, the reporter uses the terminal Tol2 repeat downstream of the insert as possible polyadenylation signal. *pCB54*_*crybb1:TagBFP*, containing the zebrafish *crybb1* promoter and parts of the minimal *CMV* promoter expressing a zebrafish codon-optimized *TagBFP* ORF flanked by *BglII* restriction sites, was synthesized (GENEWIZ). After sequence verification, the fragment was cloned into the *Tol2*-based Multisite Gateway backbone *pDestTol2CG2* (Tol2kit *#395*) (Kwan et al., 2007) via the *BglII* restriction site.

The construction of cavefish-tested *pDEST_exorh:mGreenLantern* and *pDEST_exorh:mCherry* was performed using the NEBuilder HiFi DNA Assembly Cloning Kit (NEB E5520S). The original vector *pDestTol2A2* (Tol2kit *#394*) (Kwan et al., 2007) was digested with BglII (NEB R0144S) and HpaI (NEB R0105S) restriction enzymes. The digested plasmid was purified using the Wizard SV Gel and PCR Clean-Up System (Promega A9281). The zebrafish *exorh* promoter region was amplified from *pCK029* (this manuscript) using the following primers *HpaIBgl2_NEB_exorhprom:*

*o fwd:*

*5’- GCCACAGGATCAAGAGCACCCGTGGCCGTATCTTCGCAGCTCAGCTGGCAGTACTACCGC-3’*

*o rev:*

*5’-*

*cccttgctcaccatggtggcGATGGAGAAGTGGACGATCGG-3’*.

The mGreenLantern and mCherry fluorescent protein-coding sequences were amplified from *pTol2_tg:mGreen Lantern/mCherry* (a kind gift from Dr. Sumeet Pal Singh, IRIBHM, Belgium) using the following designed primers:

*HpaIBgl2_NEB_mGL_ o fwd*:

*5’- CGATCGTCCACTTCTCCATCgccaccatggtgagc-3’ o rev*:

*5’- TATTTGTAACCATTATAAGCTGCAATAAACAAGTTttactt*

*gtacagctcgtccatgtca-3’ HpaIBgl2_NEB_mCherry o fwd*:

*5’-CGATCGTCCACTTCTCCATCgccaccatggtgagc-3’; o rev*:

*5’- TGCTTTATTTGTAACCATTATAAGCTGCAATAAACAAG*

*TTttacttgtacagctcgtccatgcc-3’*

Purified amplicons were ligated to the digested *pDestTol2A2* according to the NEB HiFi DNA assembly protocol.

### RNA Synthesis

Tol2 transposase-encoding capped mRNA was created by *in vitro* transcription using the SP6 mMessage mMachine Kit (Ambion, AM1340) from *pCS2+Tol2* plasmid (Kwan et al., 2007) linearized with NotI restriction digest. RNA was purified with Lithium Chloride precipitation followed by the Megaclear Kit (Ambion AM1908).

### Zebrafish Injections and Transgenesis

All plasmids were miniprepped with the ZymoPURE plasmid miniprep kit (ZYMO D4212). Embryos were injected into the early cell of one-cell-stage wildtype embryos (*AB* or *TU*) with 1 nL injection mix containing plasmid DNA concentrations of 25 ng/mL and Tol2 mRNA at 25 ng/mL. Adult F0 animals mosaic for the transgene were outcrossed to wildtype and their progeny analyzed for stable transgene expression. Tol2-based zebrafish transgenics were generated using standard experimental protocols as specified in Felker and Mosimann (2016) to achieve single-insertion transgenics and reproducible quality control.

### Imaging

Embryos or larvae were anesthetized with 0.016% Tricaine-S (MS-222, Pentair Aquatic Ecosystems, Apopka, Florida, NC0342409) in E3 embryo medium. Basic fluorescence imaging was performed on a Leica M205FA with a DFC450 C camera. Laser scanning confocal microscopy was performed on a Zeiss LSM880 following embedding in E3 with 1% low-melting-point agarose (Sigma Aldrich, A9045) on glass bottom culture dishes (Greiner Bio-One, Kremsmunster, Austria, 627861). Images were collected with a 10/0.8 air-objective lens with all channels captured sequentially with maximum speed in bidirectional mode, with the range of detection adjusted to avoid overlap between channels. Maximum projections of acquired Z-stacks were made using ImageJ/Fiji (Schindelin et al., 2012; Schneider et al., 2012) and cropped and rotated using Adobe Photoshop 2022.

## Supplemental Movie 1

Adult *exorh:EGFP*-transgenic zebrafish observed using a NightSEA hand-held UV flashlight and UV filter glasses (“goggles”), demonstrating the persistent pineal gland expression of the reporter and the simple detection.

## Acknowledgements

We thank Christine Archer, Molly Waters, and Nikki Tsuji for zebrafish husbandry support, and all members of the Mosimann, Burger, Kwan, and Lovely labs for critical input on tool design, cloning, and the manuscript. We thank Dr. Stephanie Woo, Dr. Sumeet Pal Singh, and Dr. Holger Gerhard for sharing plasmids. We are especially grateful to Dr. Yoshitaka Fukuda and Dr. Daisuke Kojima for sharing the isolated zebrafish *exorh* promoter sequence with us.

## Author contributions

All plasmid vectors generated by the Mosimann, Burger, Kwan, and Affolter labs use plasmid designations with initials of the original cloner followed by a plasmid identifier number (*pXY#*). C.M., A.B., K.M.K. conceptualized the manuscript and the overall experiments, assisted in experiments, acquired funding, supported data gathering and interpretation, coordinated the collaborators, and co-wrote and finalized the manuscript draft; C.L.K. and H.R.M. cloned plasmid vectors including *exorh*-based reporters, performed microinjections, generated transgenic lines, assembled plasmid maps, gathered data and performed imaging, assembled figures and figure panels, and co-wrote the manuscript draft; K.M.K., B.F.M. and A.S. cloned constructs, performed microinjections, acquired imaging data and expression patterns, generated vector maps, and assembled figure panels; J.R.K., R.L.R., E.L., and C.B.L. generated the *sox17*-based reporters, acquired imaging data, assembled figure panels, and provided manuscript text; S.B. and N.R. performed and documented all Astyanax experiments, assembled figure panels, and provided manuscript text; S.N, S.B., G.D., A.P., A.F., K. D., S. B. and A.B. generated individual vectors including quality control by sequencing, generated vector maps, and performed individual validations; C. B. and M. A. g nerated the Destination vectors with crybb1-based reporters, performed validations, and supported the manuscript draft.

