## Supplementary figures and images for "Next-generation plasmids for transgenesis in zebrafish and beyond"

### 457. p3E 2A-EGFPpA.pdf

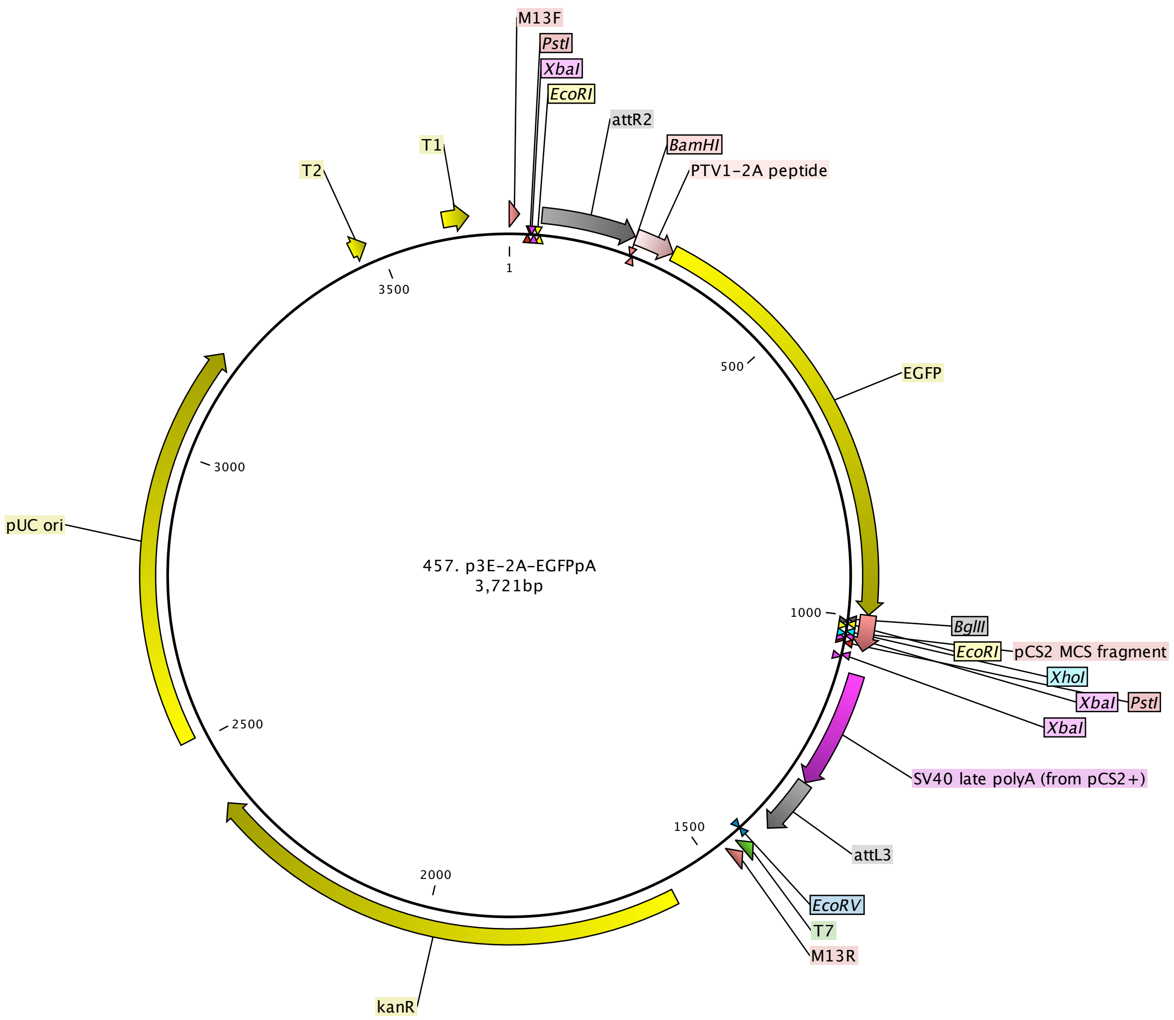

### 458 p3E 2A EGFPCAAXpA.pdf

Kemmler et al.

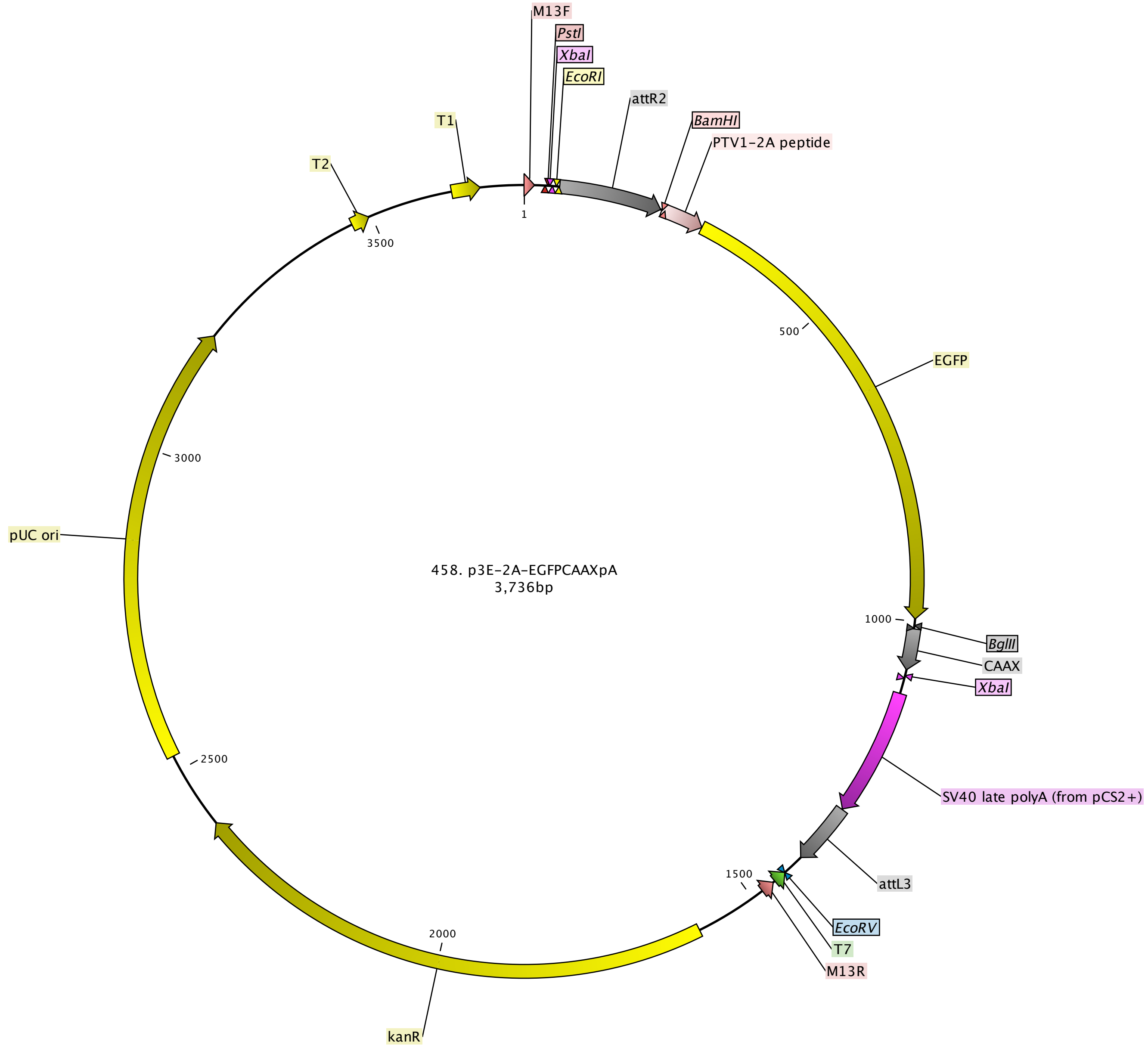

### 459. p3E 2A-nlsEGFPpA.pdf

Kemmler et al.

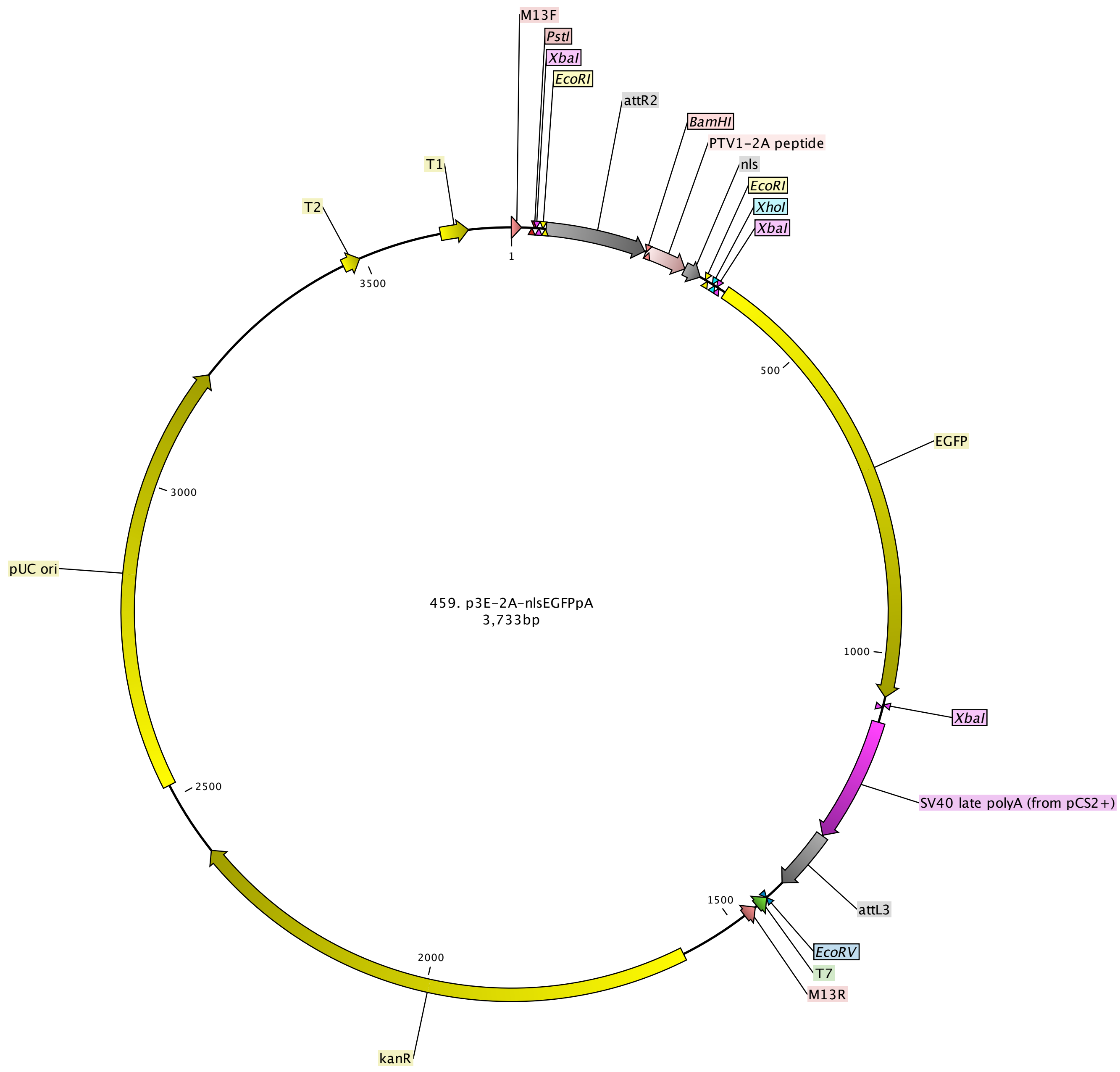

### 460 p3E 2A mcherrypA.pdf

Kemmler et al.

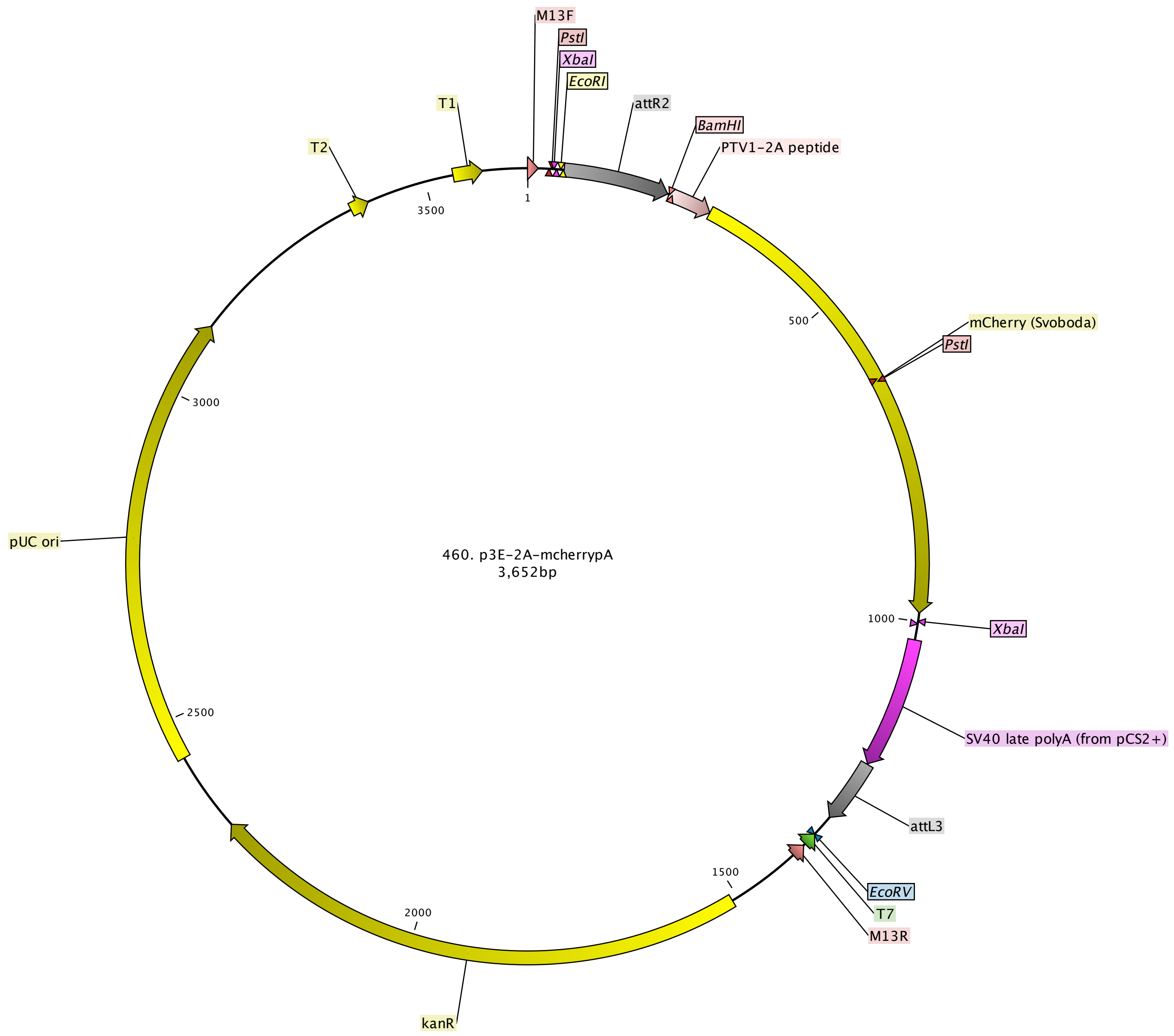

### 612. p3E 2A mCherryCAAXpA.pdf

Kemmler et al.

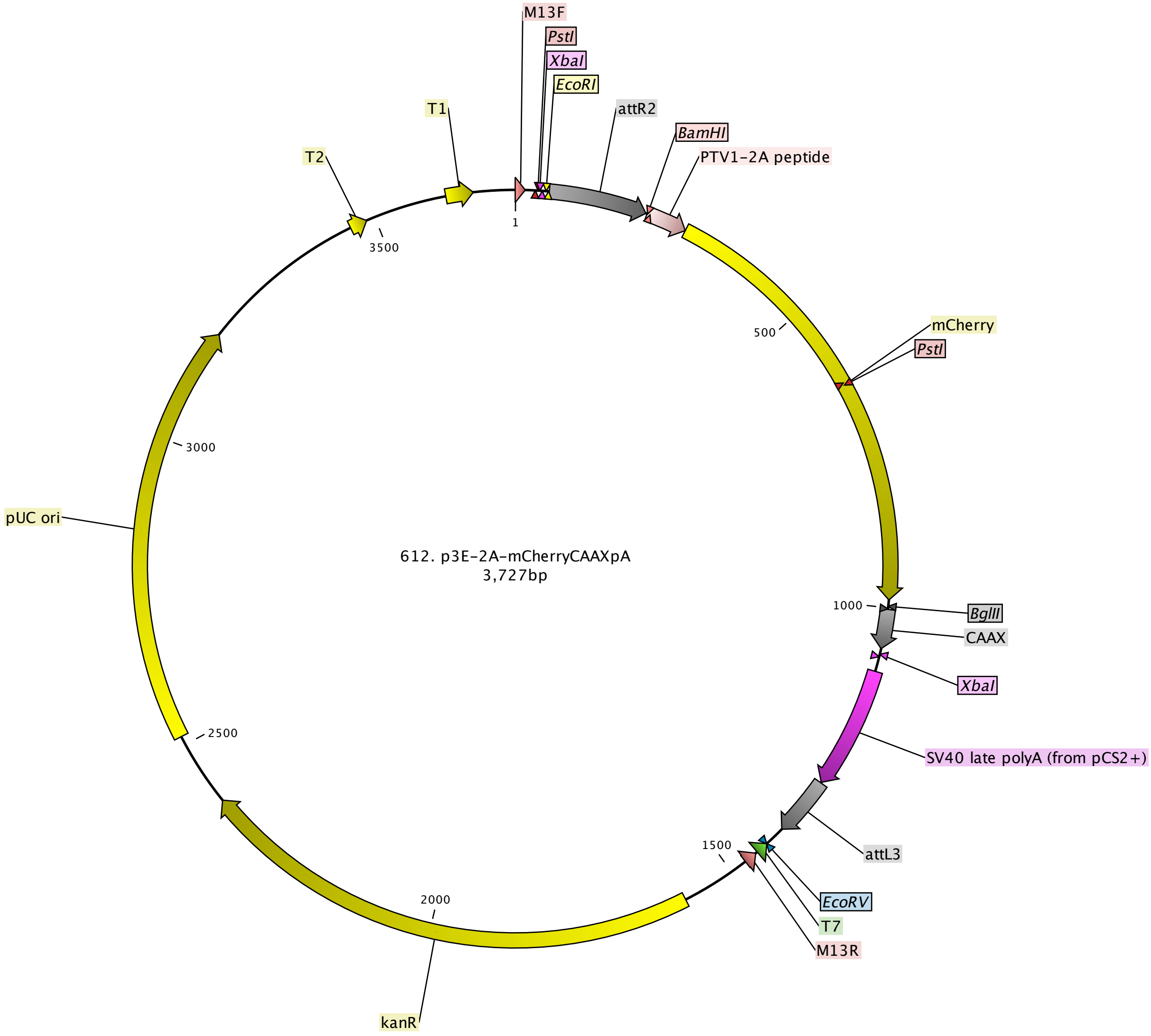

### 763 pME mApple.pdf

Kemmler et al.

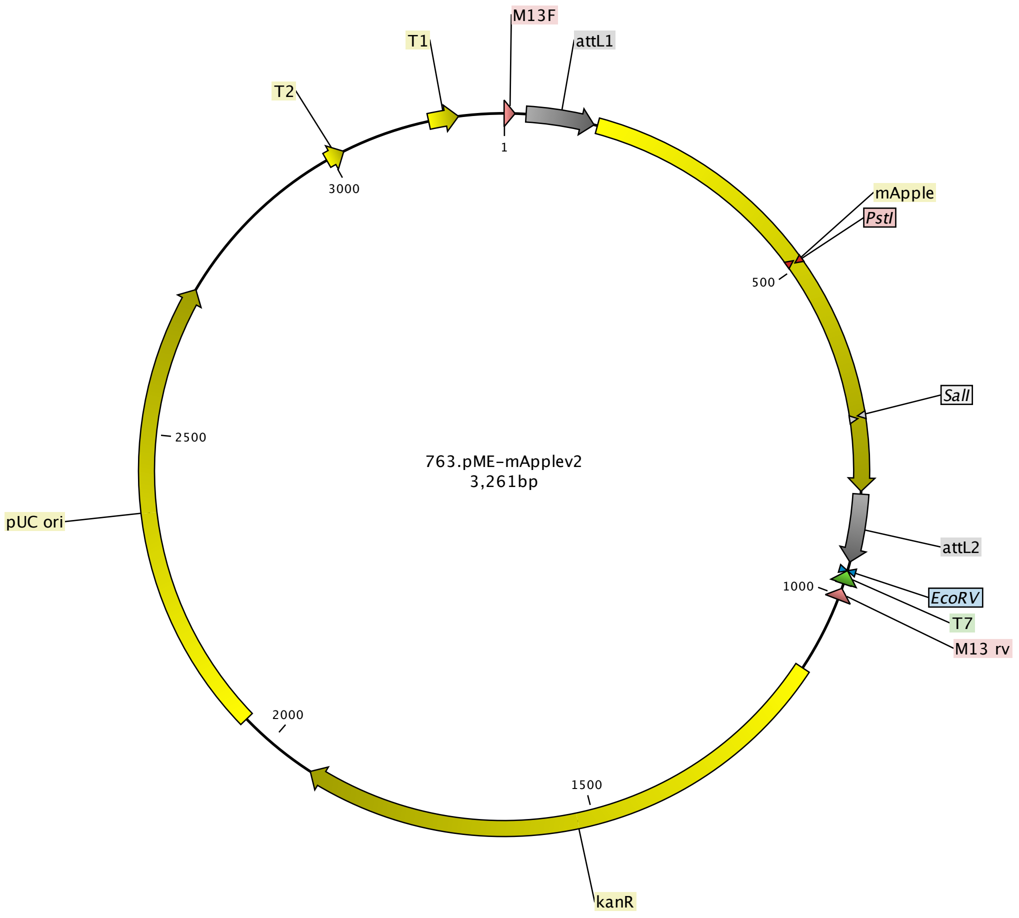

### 764 pME nls-mApple.pdf

Kemmler et al.

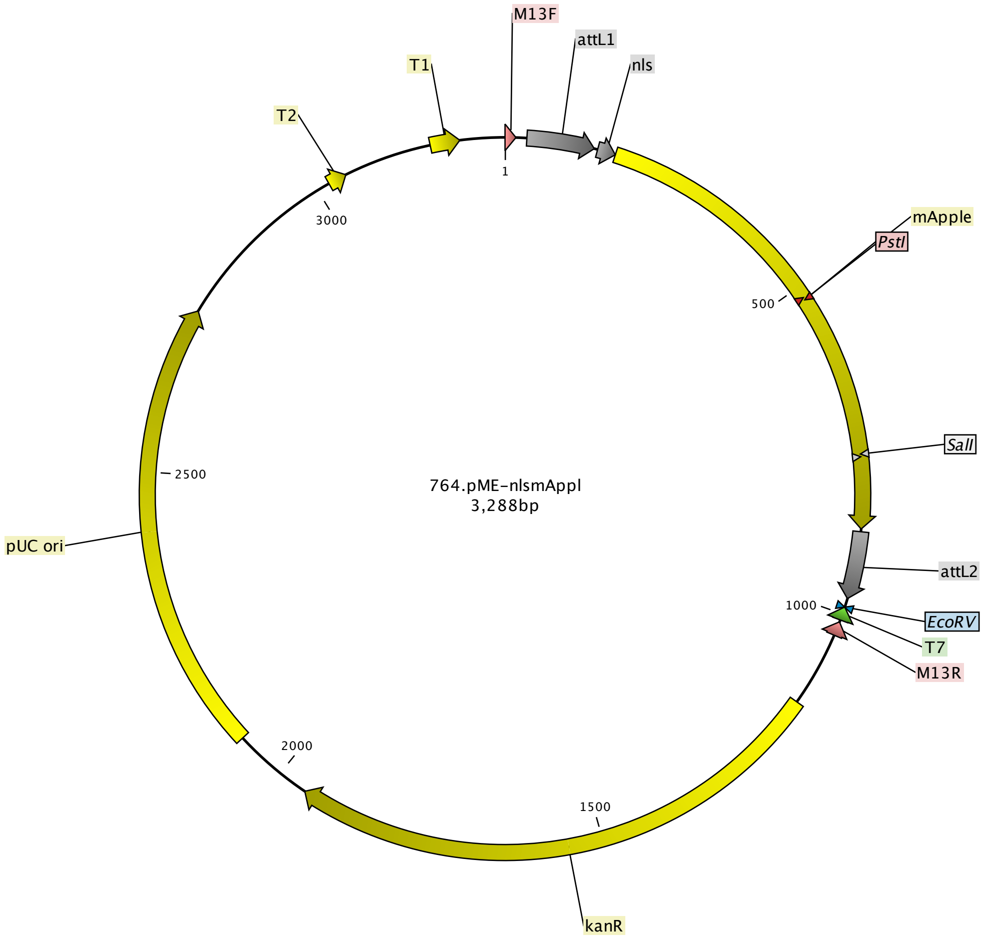

### 765 pME nls-mCerulean.pdf

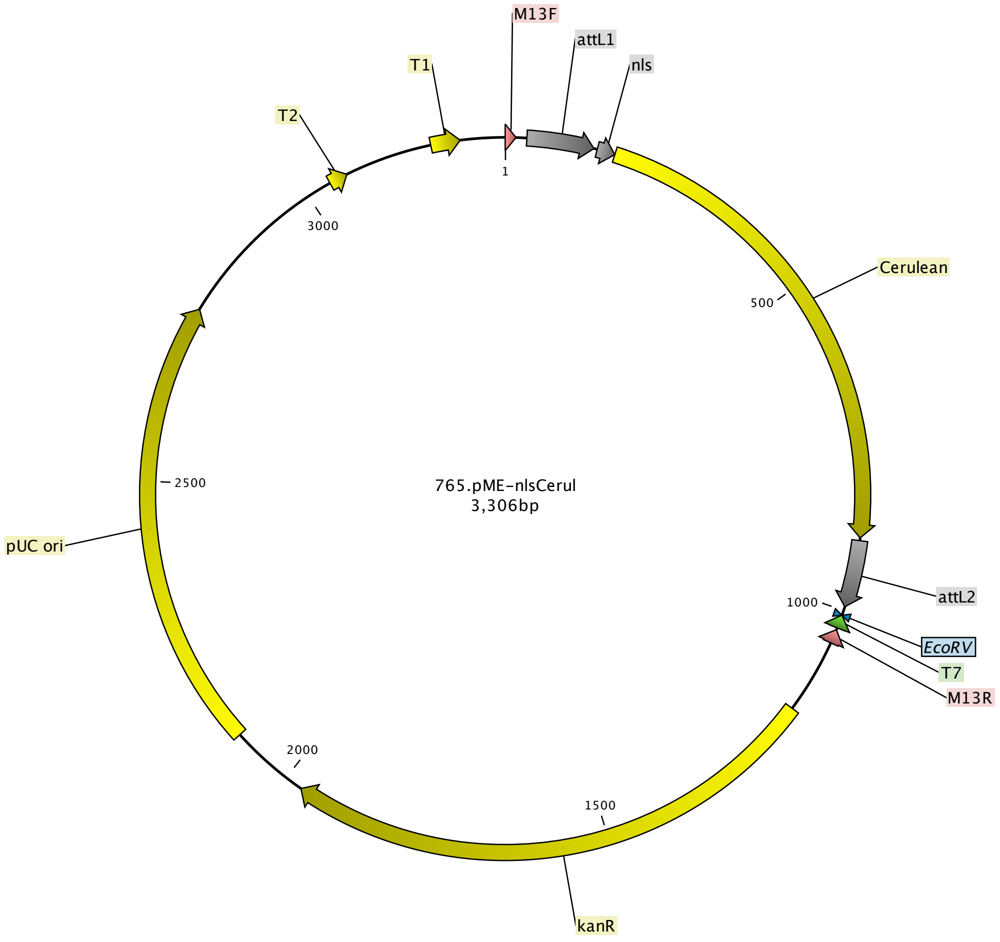

### 766. p3E 2A nlsmCherrypA.pdf

Kemmler et al.

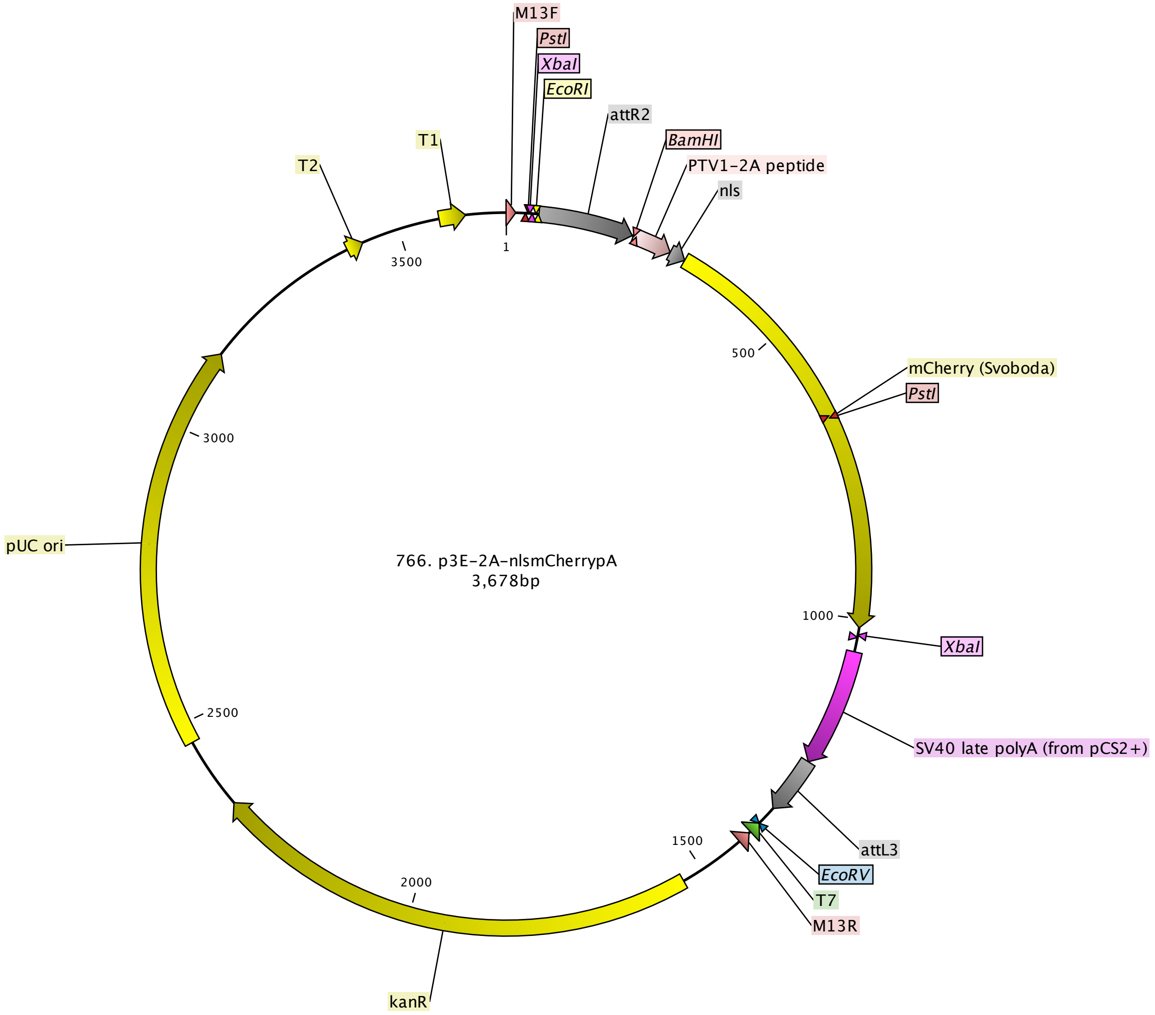

### 768 pME mAppleCAAX.pdf

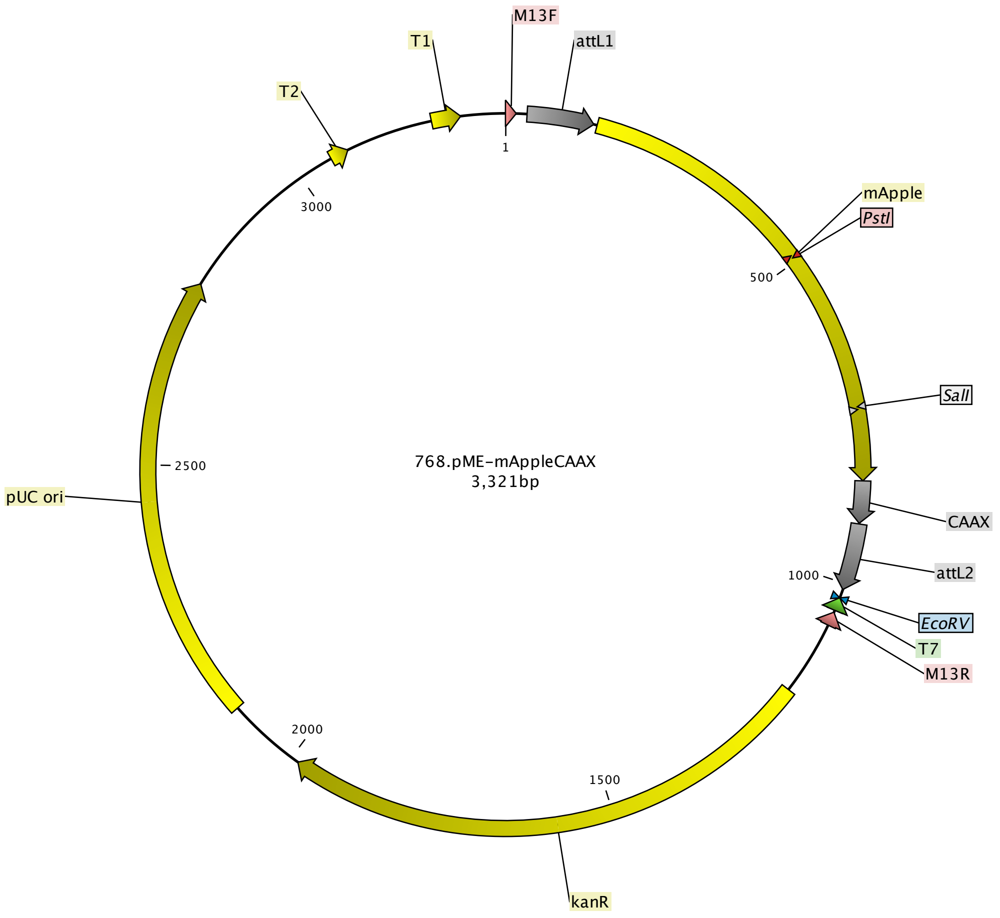

### 769 pME Cerulean CAAX.pdf

Kemmler et al.

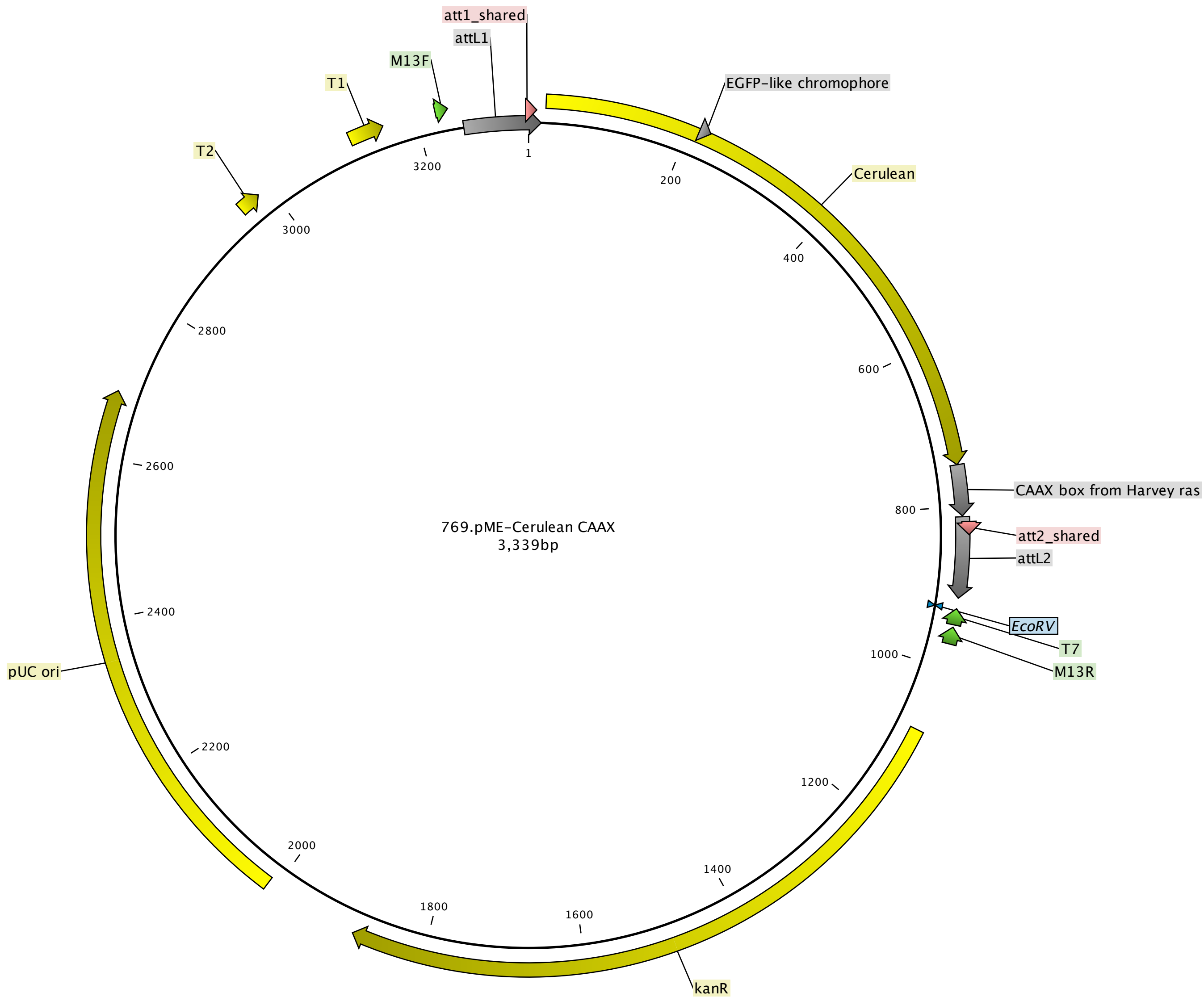

### 791. p3E 2a pCS2 pA.pdf

Kemmler et al.

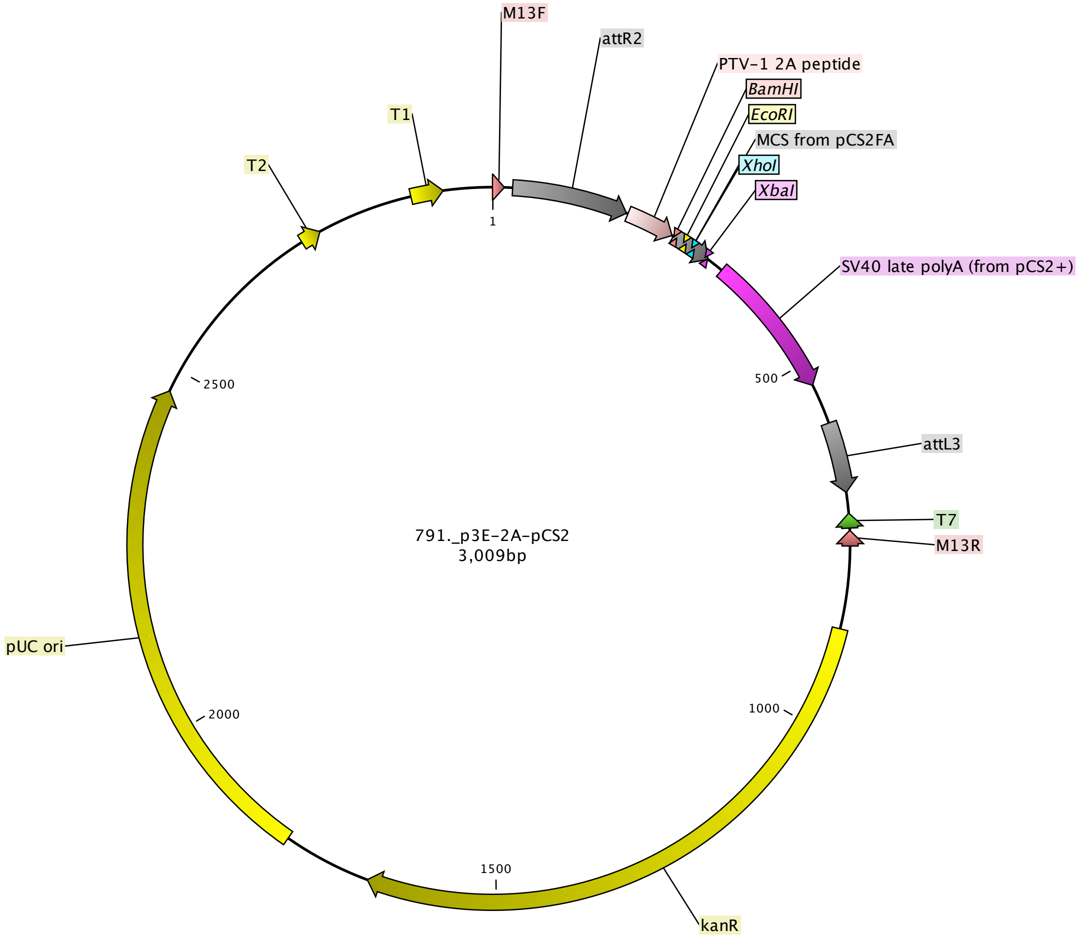

### p3E ubb polyA.pdf

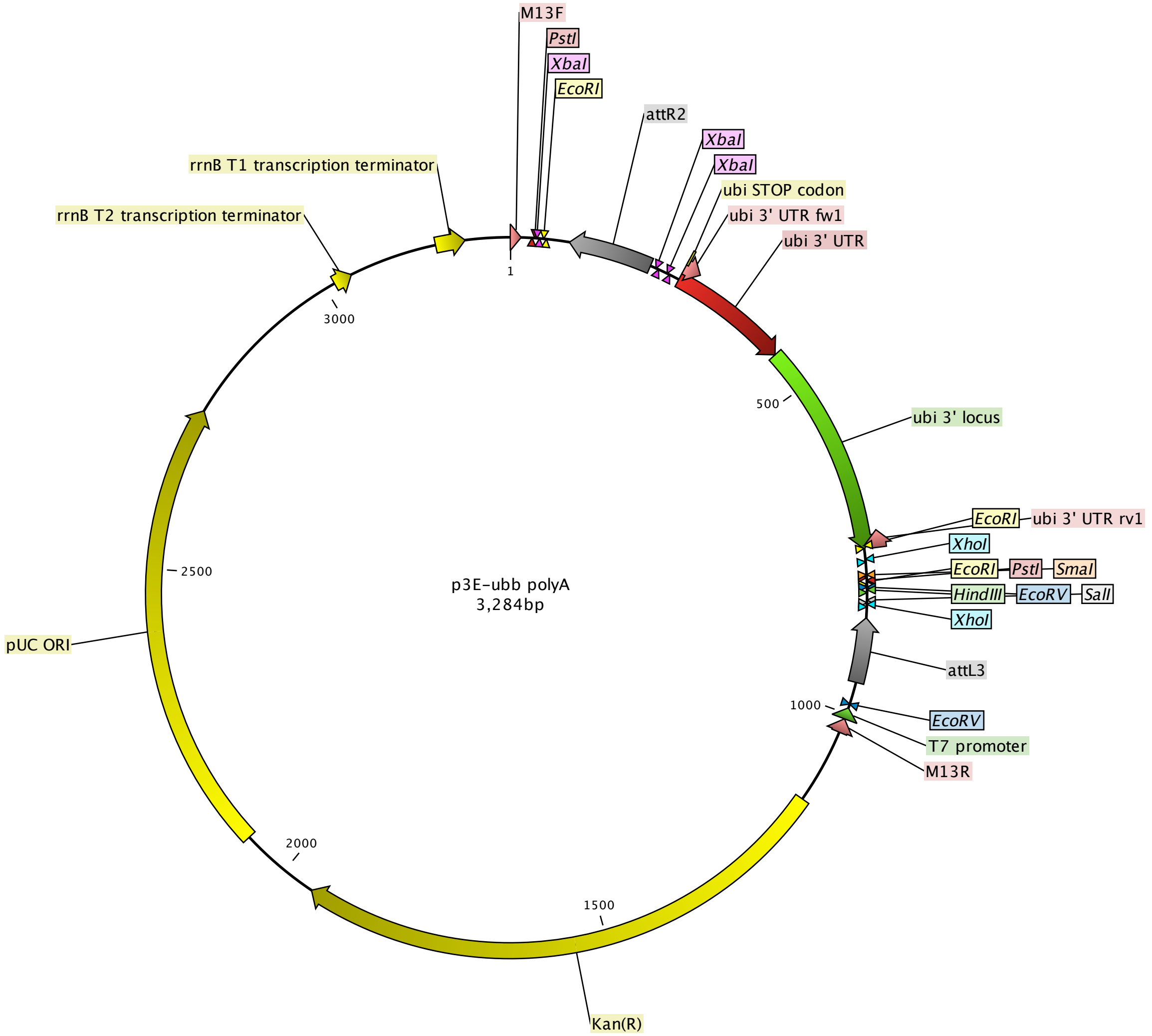

### pAF018 p3E 2A mCeruleanCAAXpA.pdf

Kemmler et al.

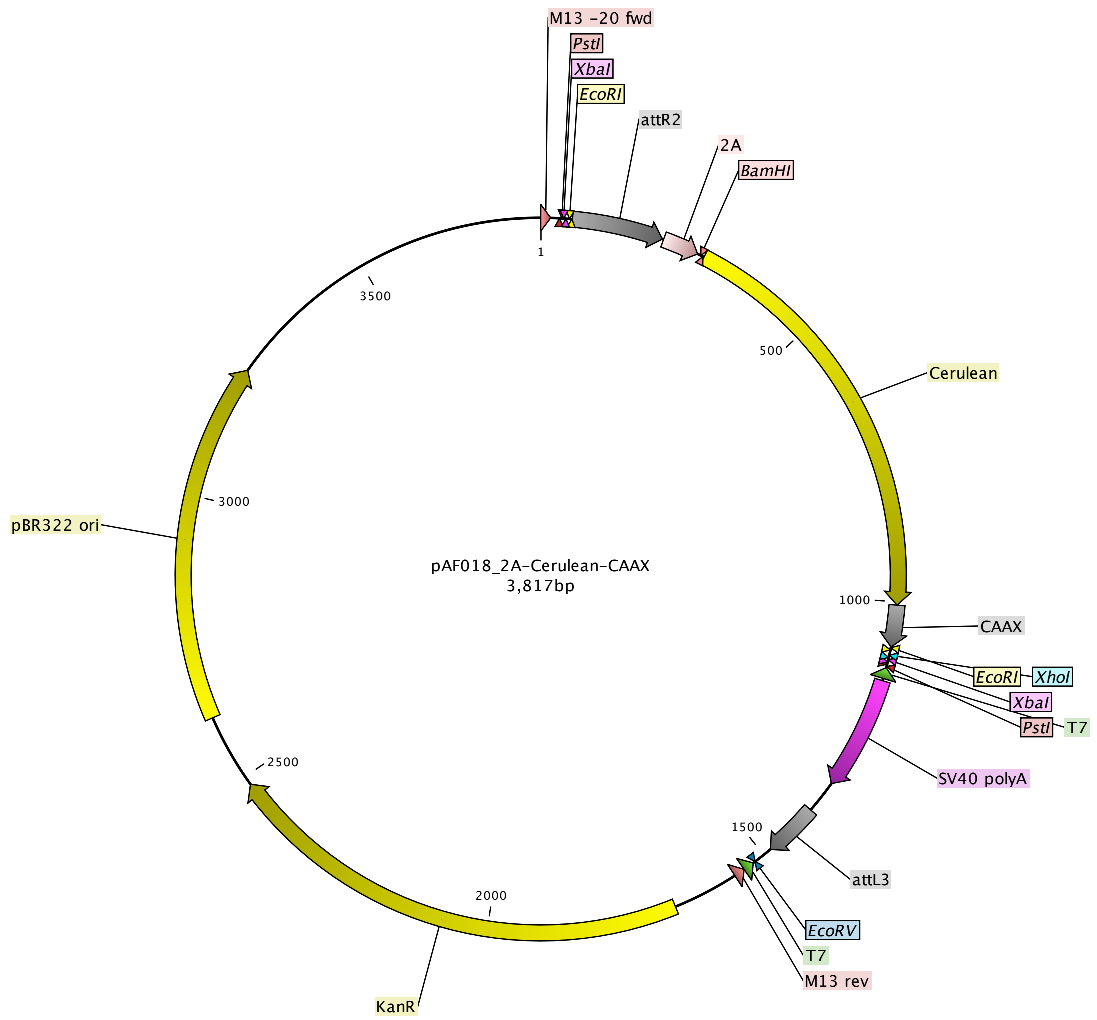

### pAF020 p3E 2A mCeruleanpA.pdf

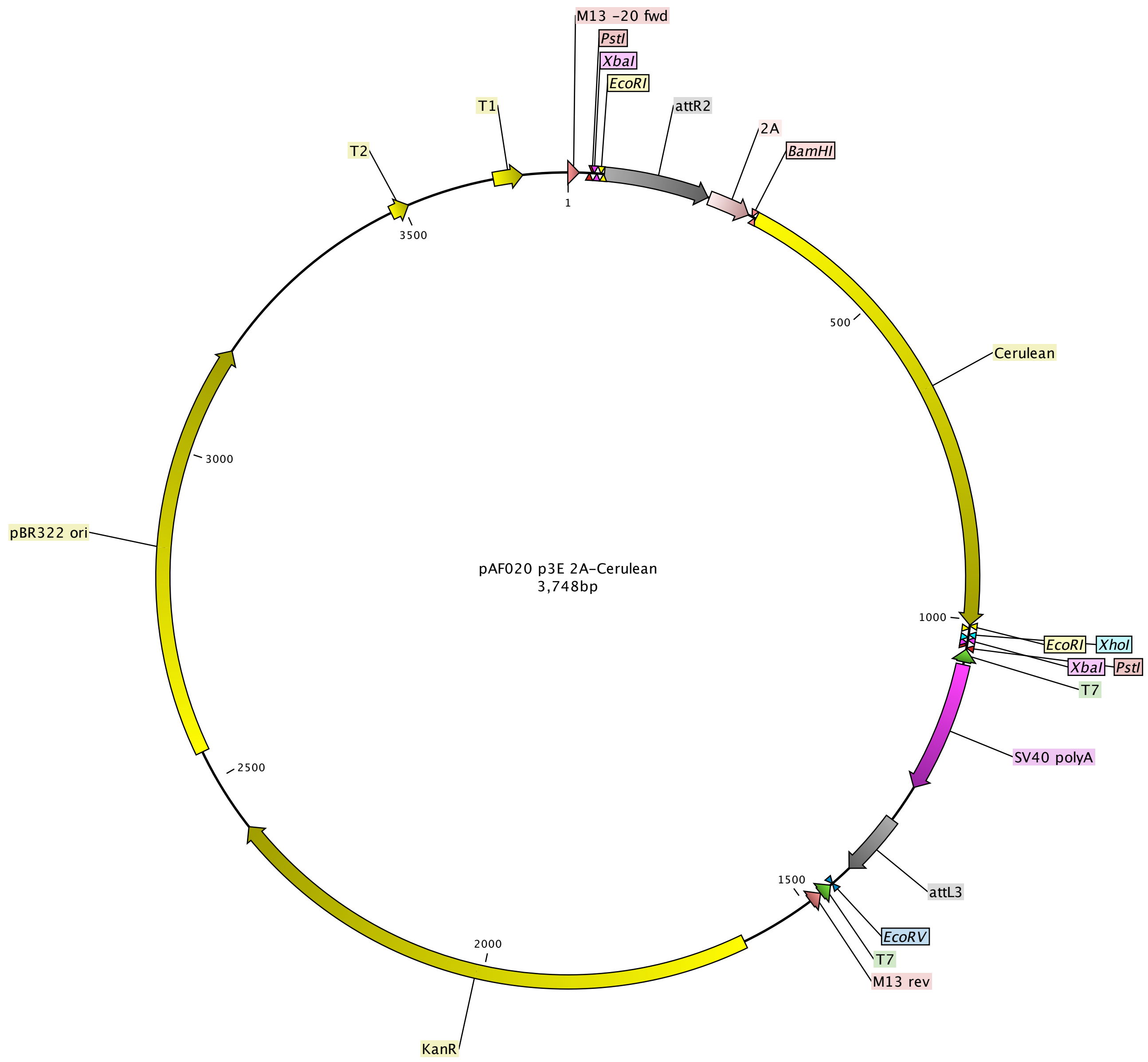

### pAP02 p3E MCS ubi 3' UTR.pdf

Kemmler et al.

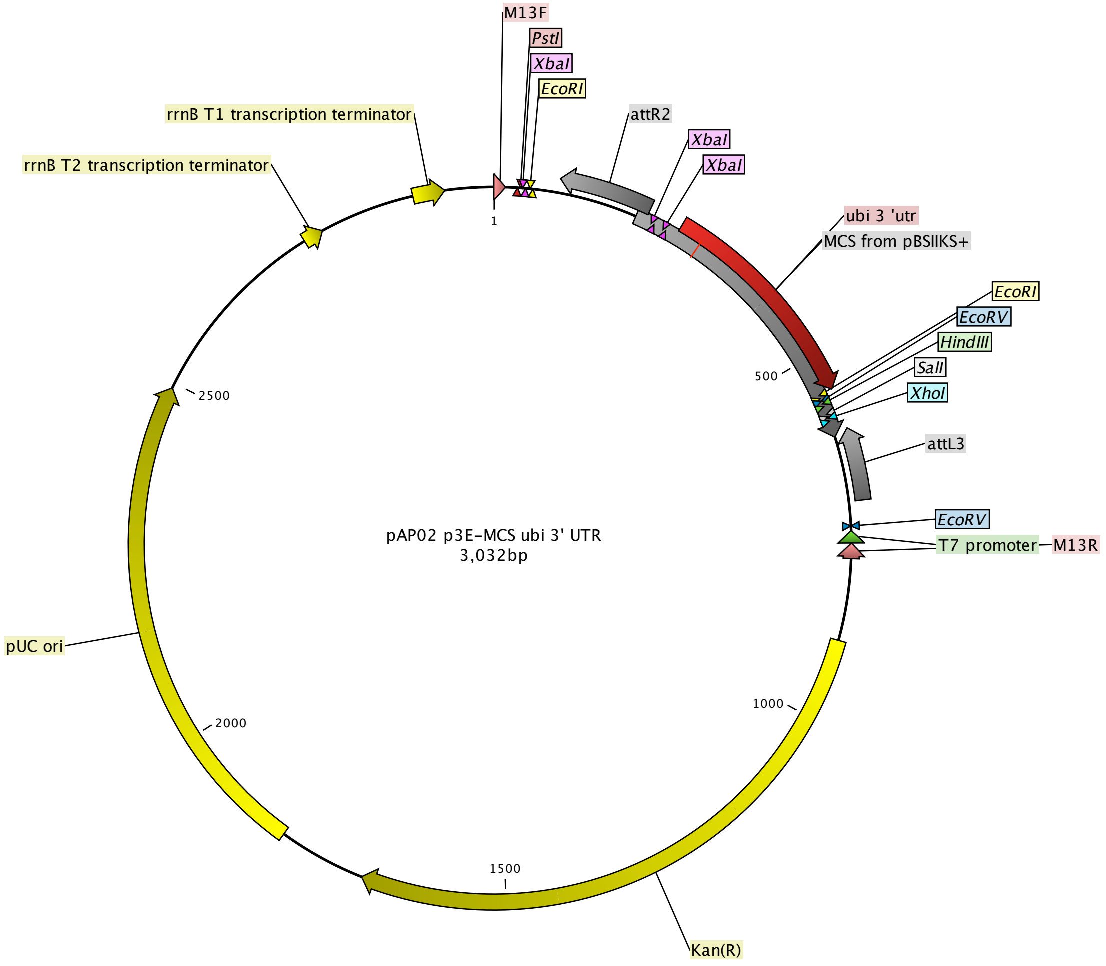

### pCB24 pDEST crybbTagBFP.pdf

Kemmler et al.

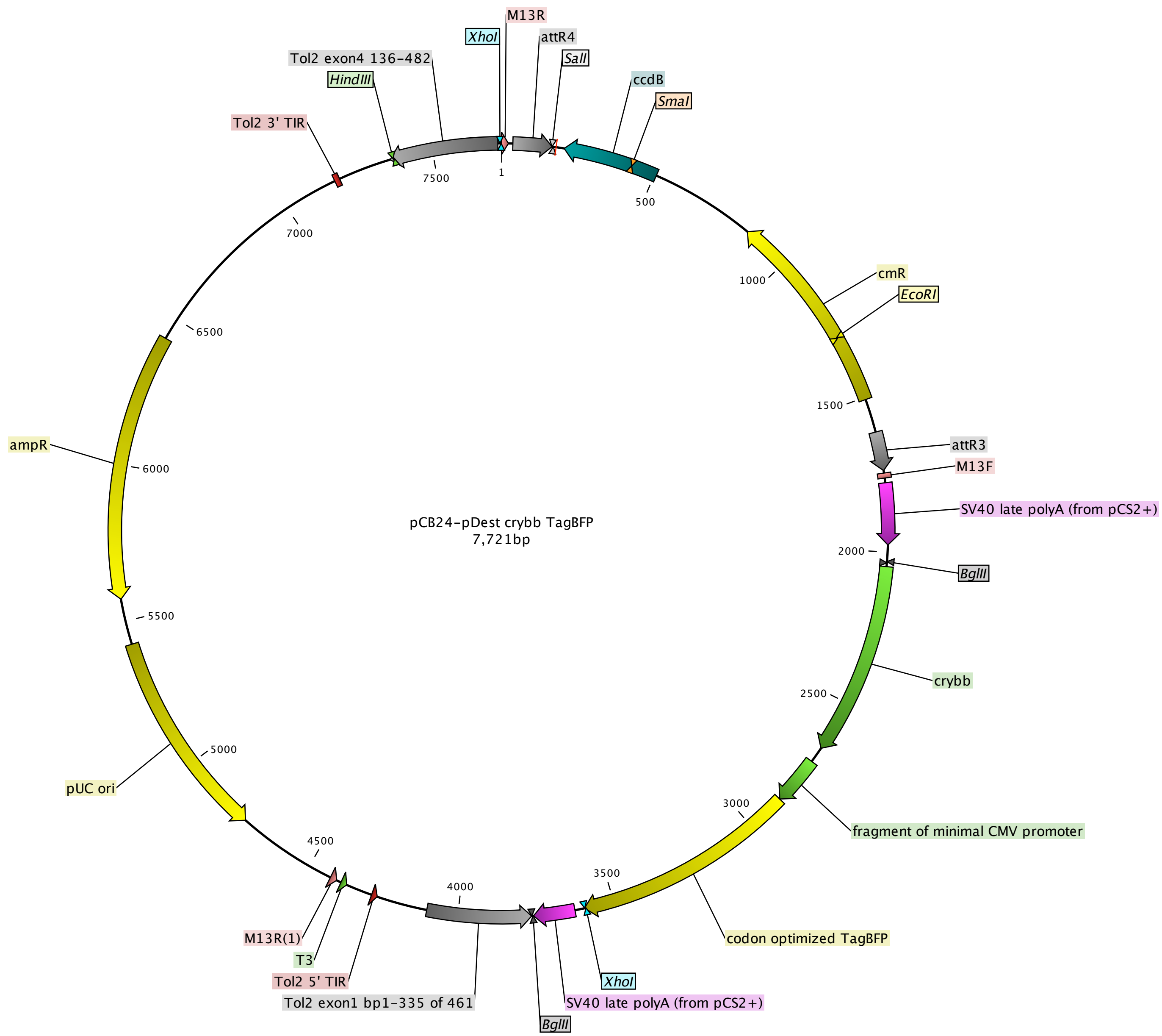

### pCB59 pDEST crybb mKate2.pdf

Kemmler et al.

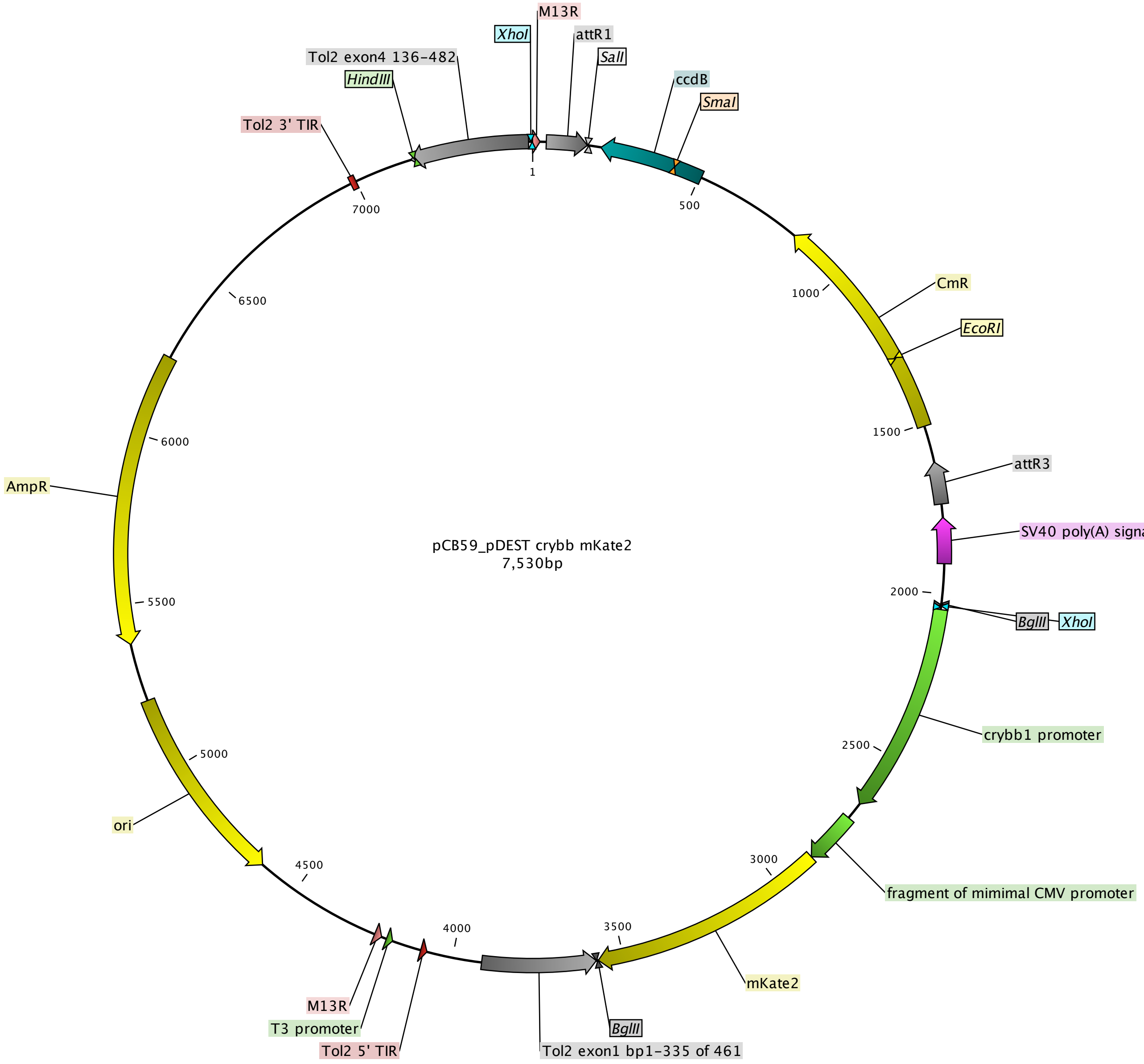

### pCK002 pME mCerulean no stop.pdf

Kemmler et al.

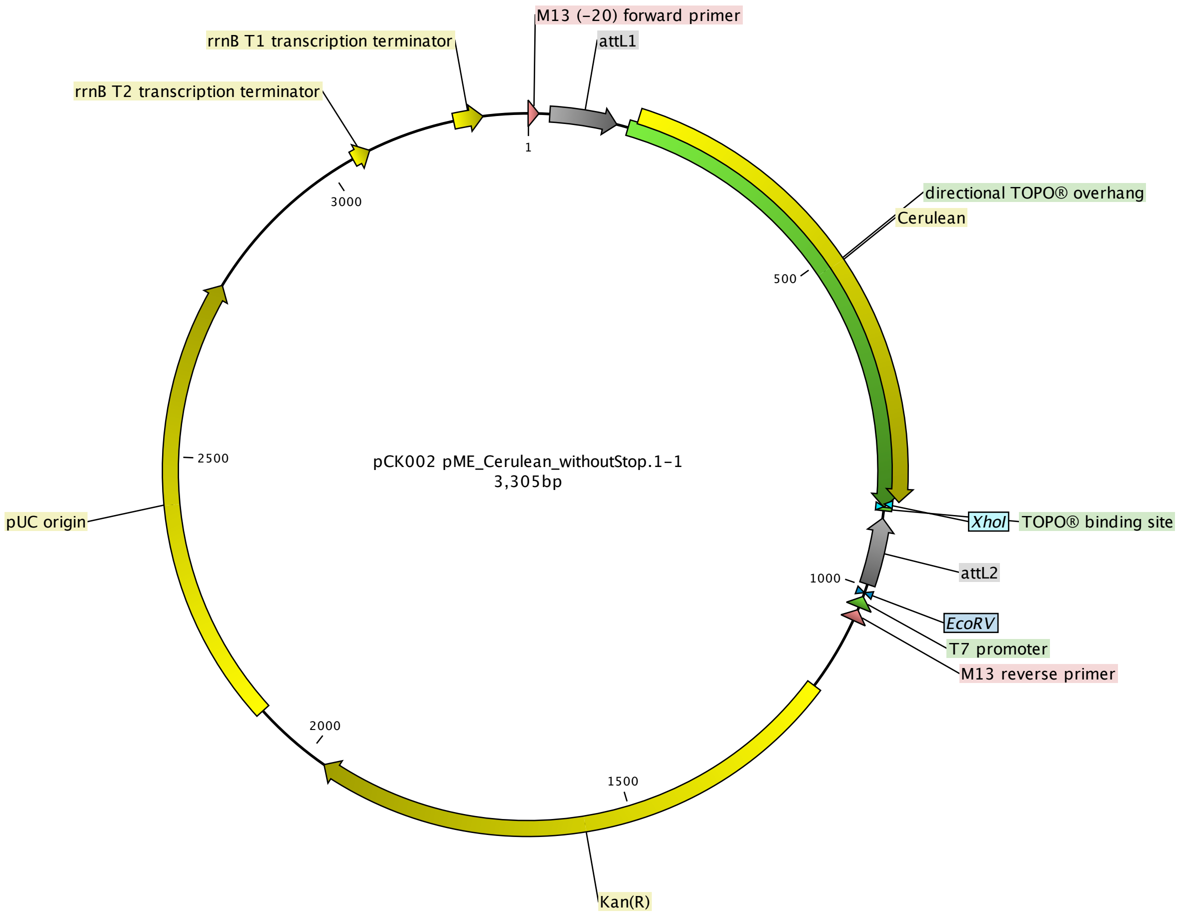

### pCK004 Ubi mCerulen 302pA.pdf

Kemmler et al.

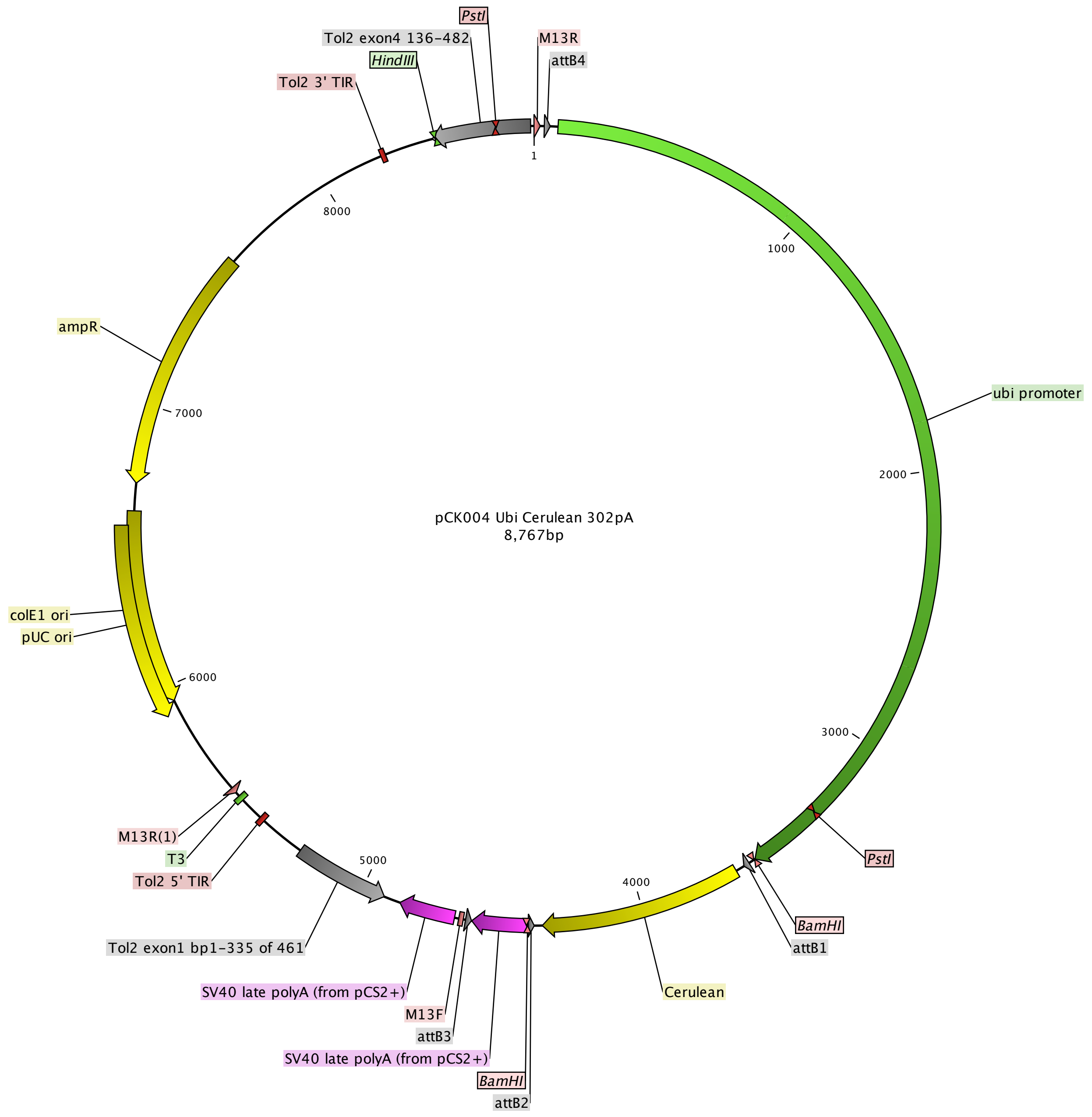

### pCK005 Ubi mCerulean ubbpA.pdf

Kemmler et al.

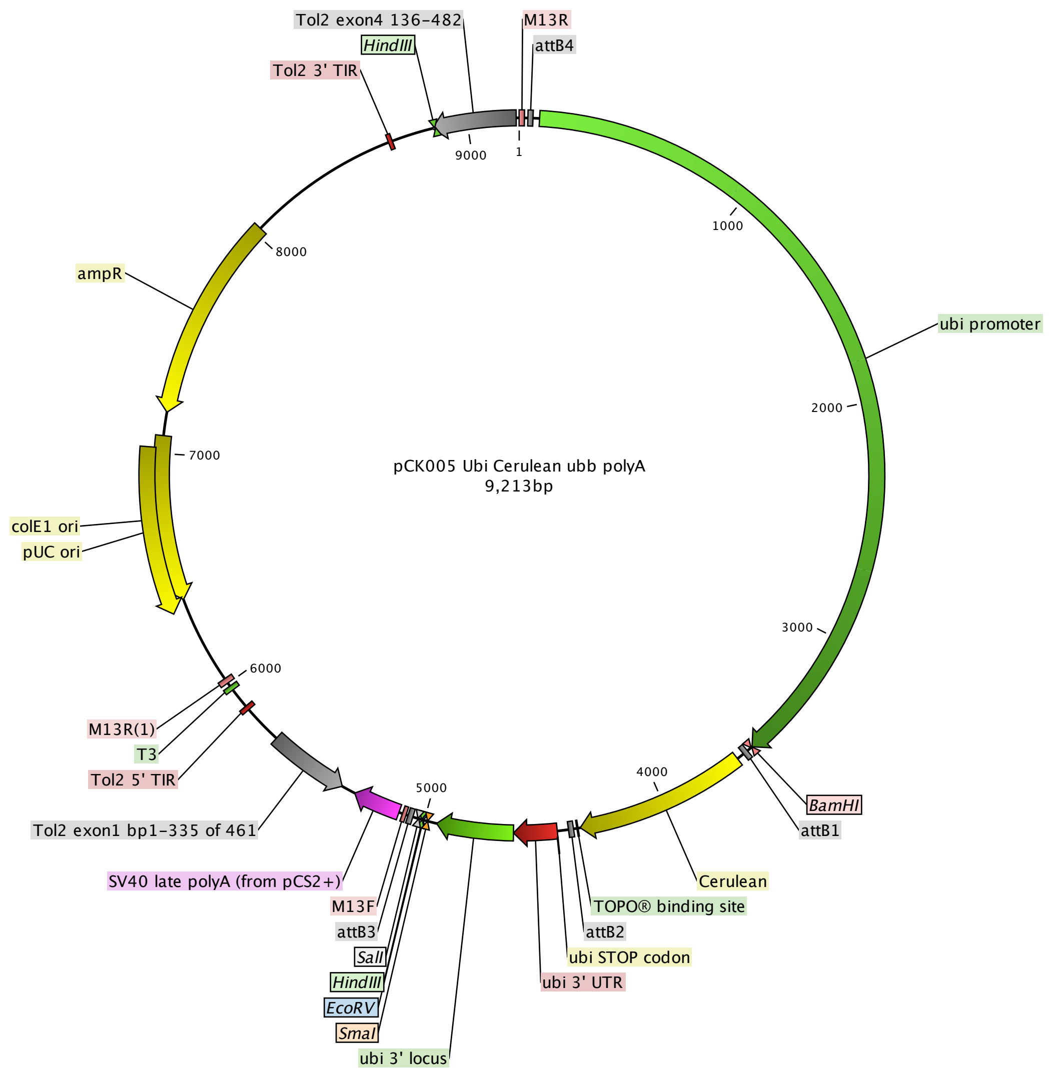

### pCK006 pME minmCherry no stop.pdf

Kemmler et al.

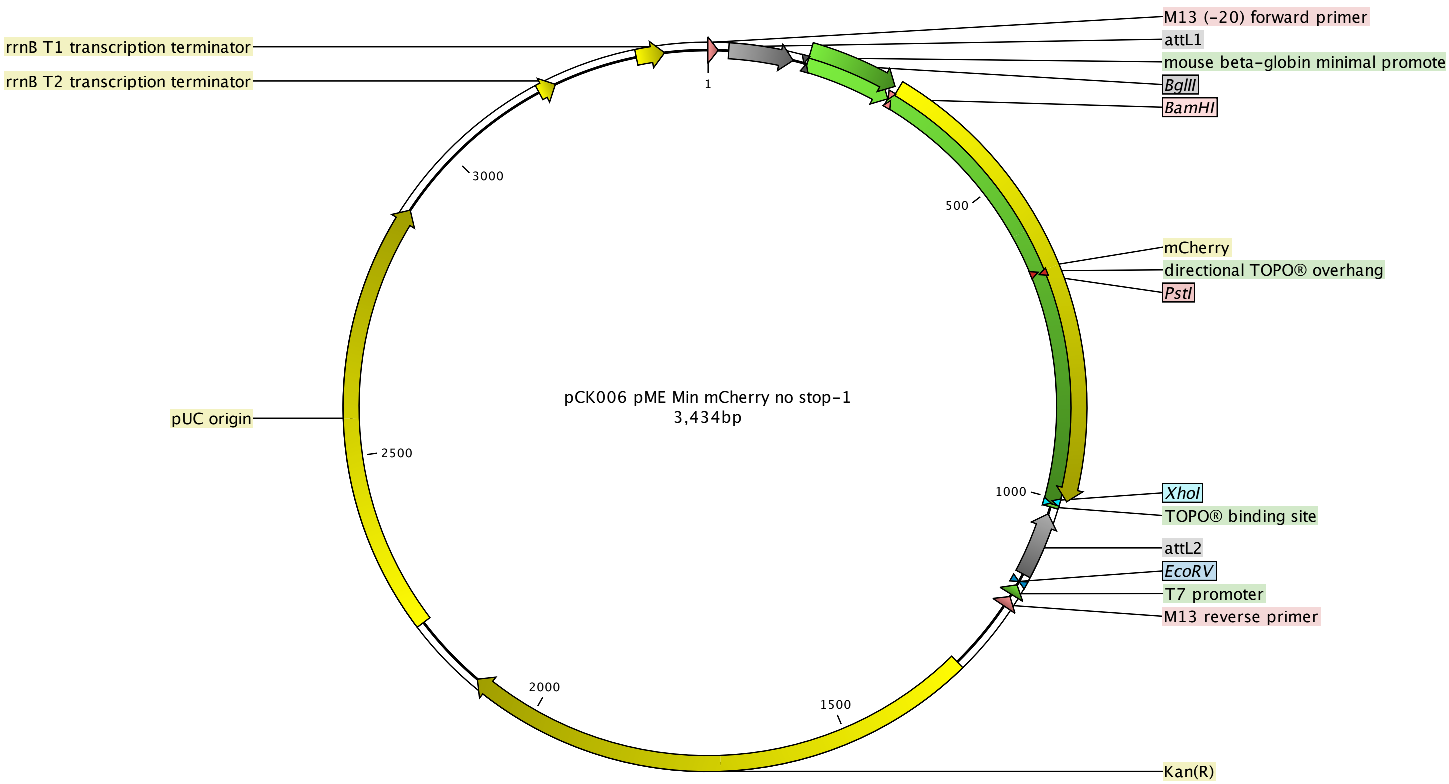

r

### pCK007 pME Kalta4.pdf

Kemmler et al.

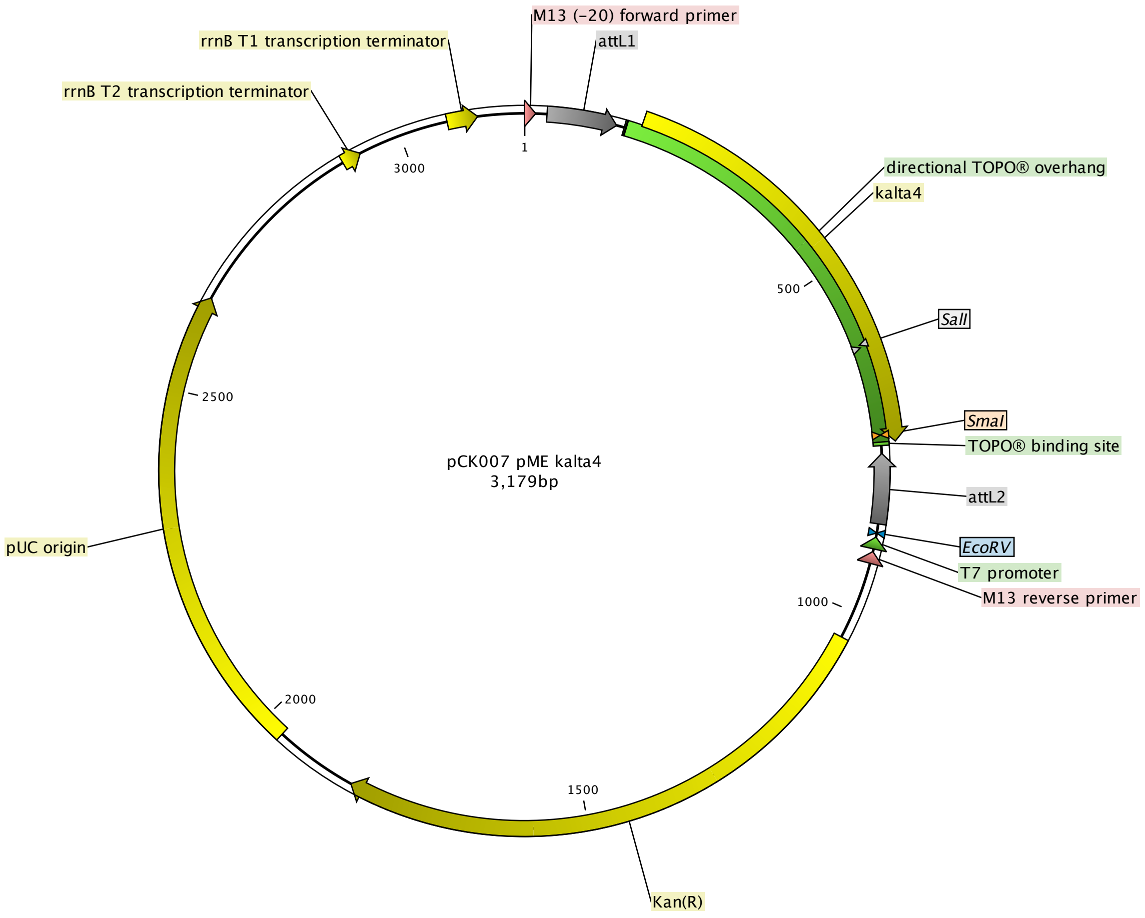

### pCK011 p5E 1055bpexorh.pdf

Kemmler et al.

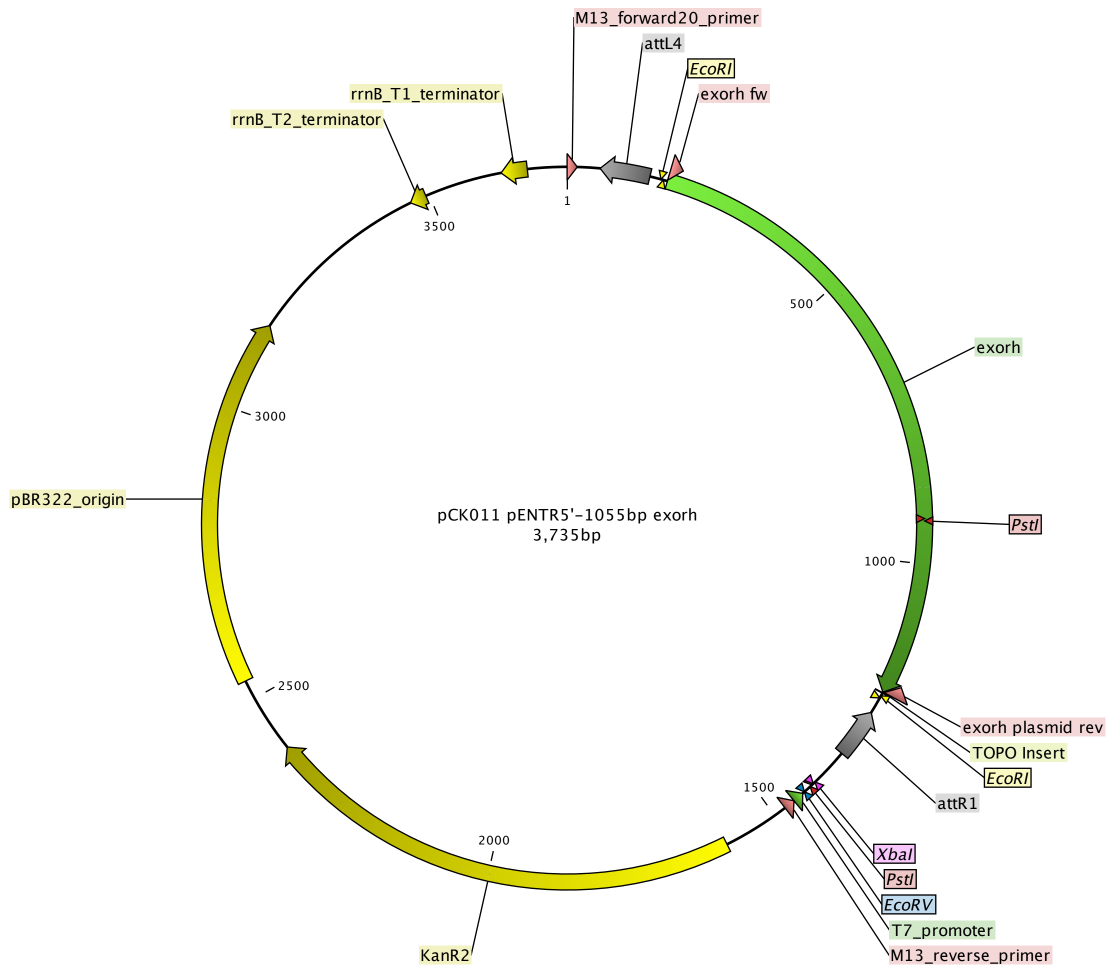

### pCK017 pDEST exorhEGFP.pdf

Kemmler et al.

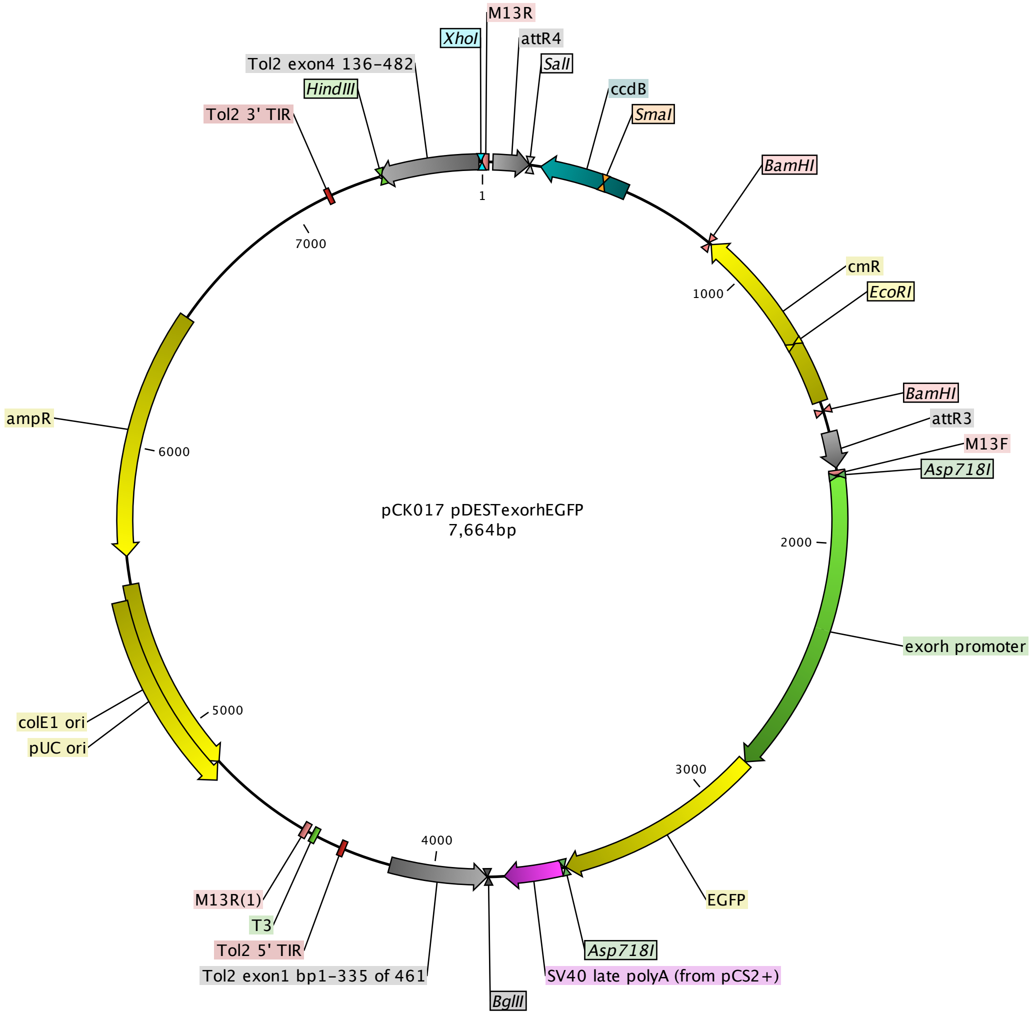

### pCK029 exorhEGFP.pdf

#### Kemmler et al.

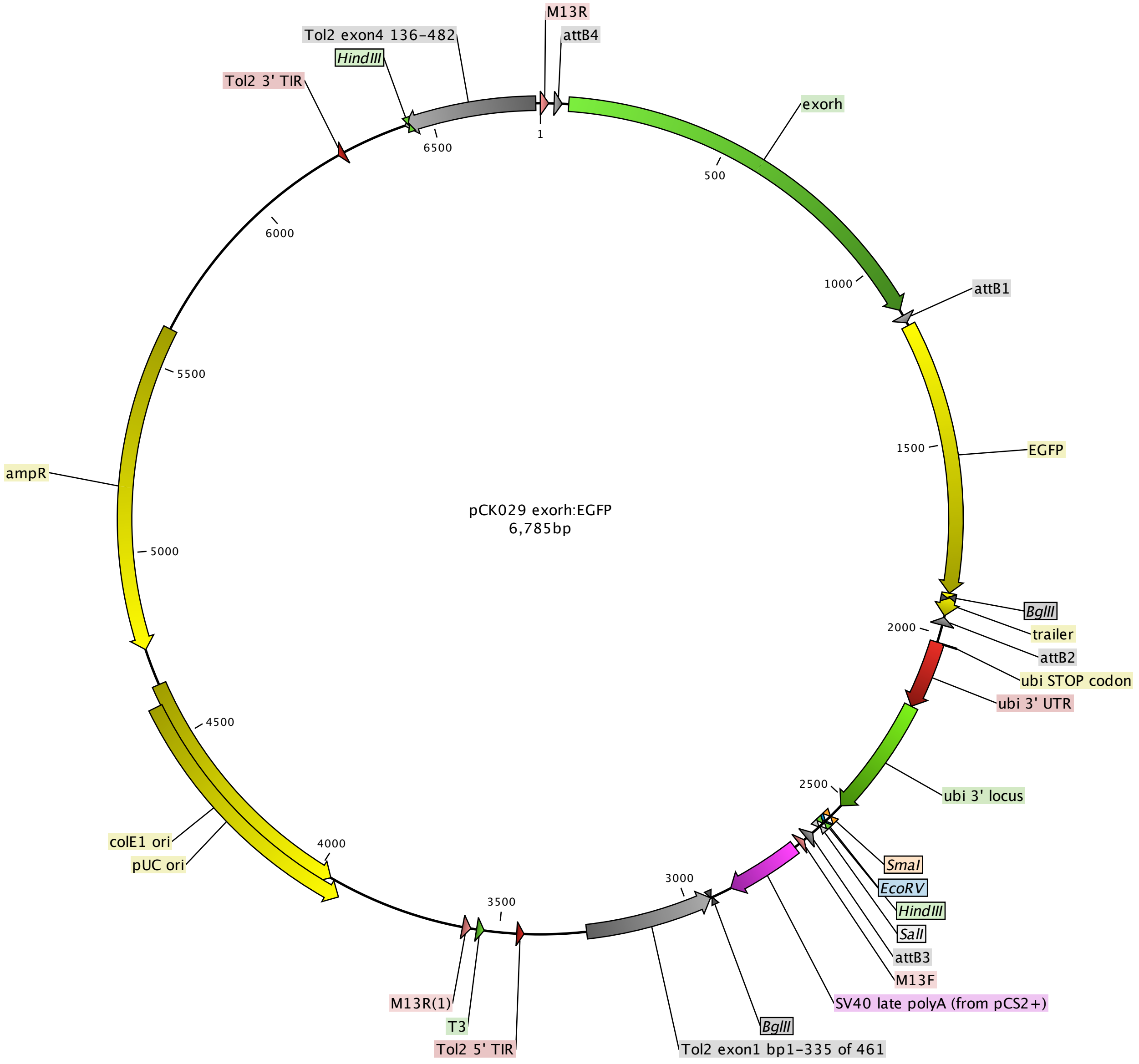

### pCK033 p5E desmin MCS.pdf

Kemmler et al.

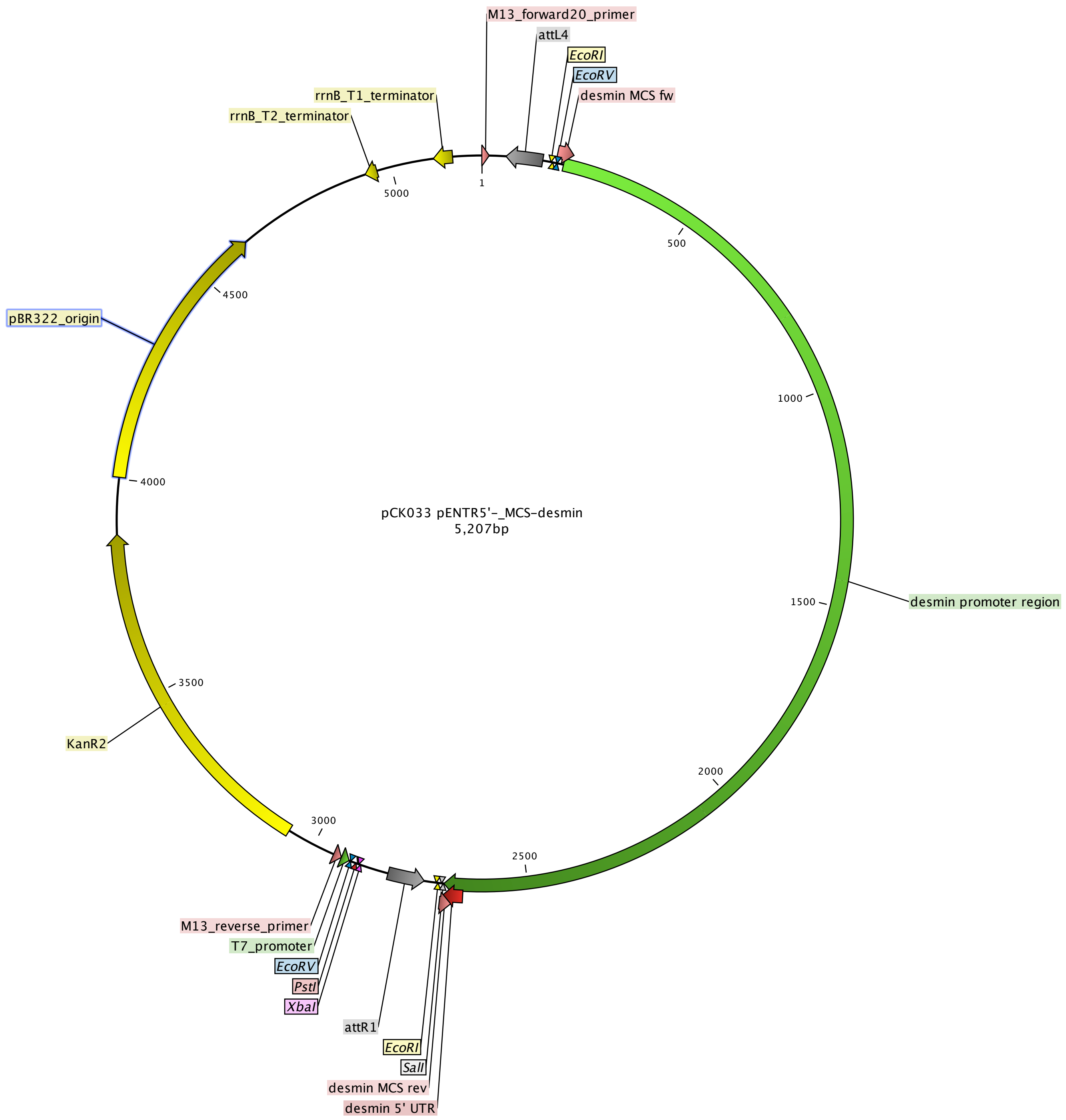

### pCK036 desminMCS-minCerulean-cryaaVenus.pdf

#### Kemmler et al.

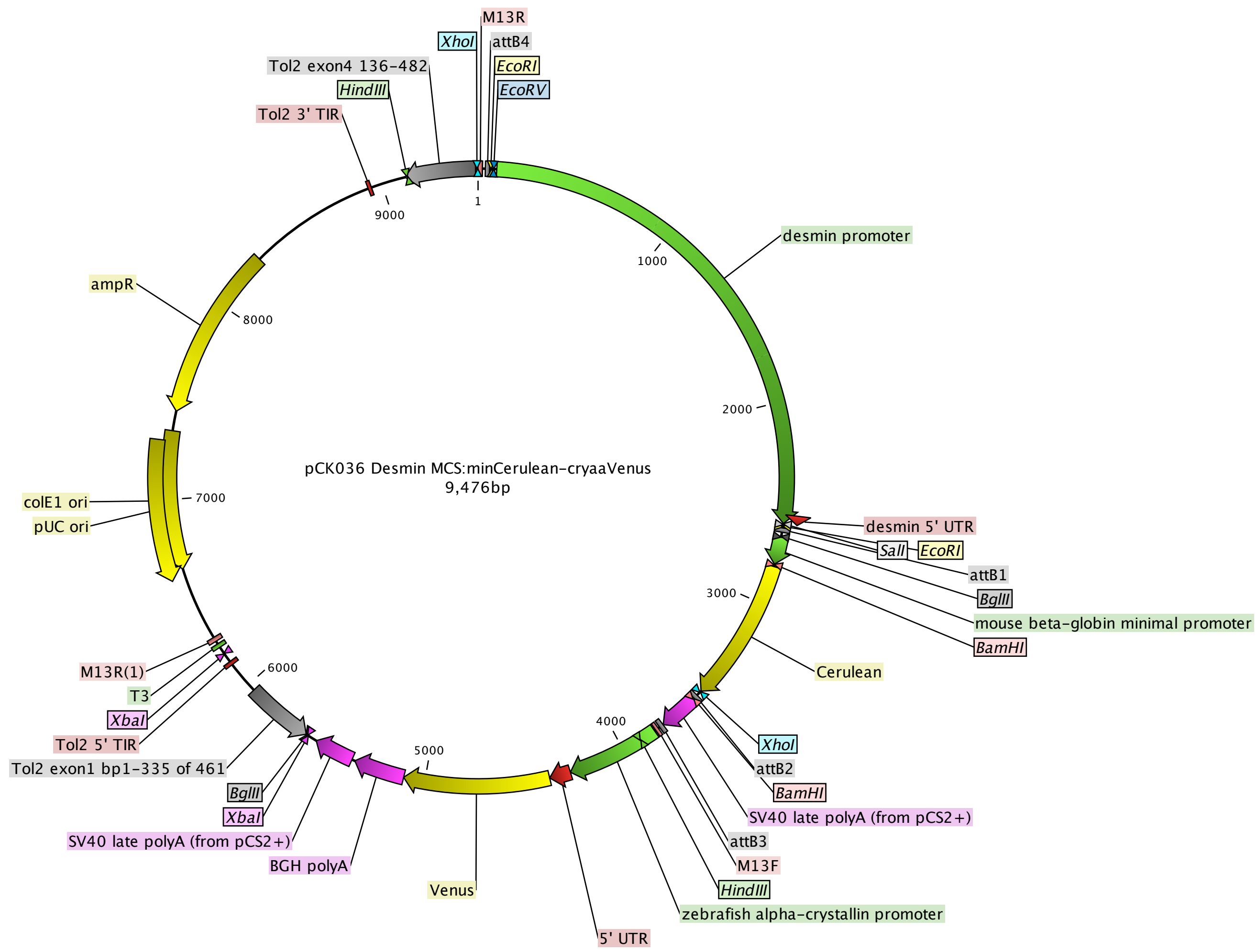

### pCK051 p5E exorh-mCerulean.pdf

Kemmler et al.

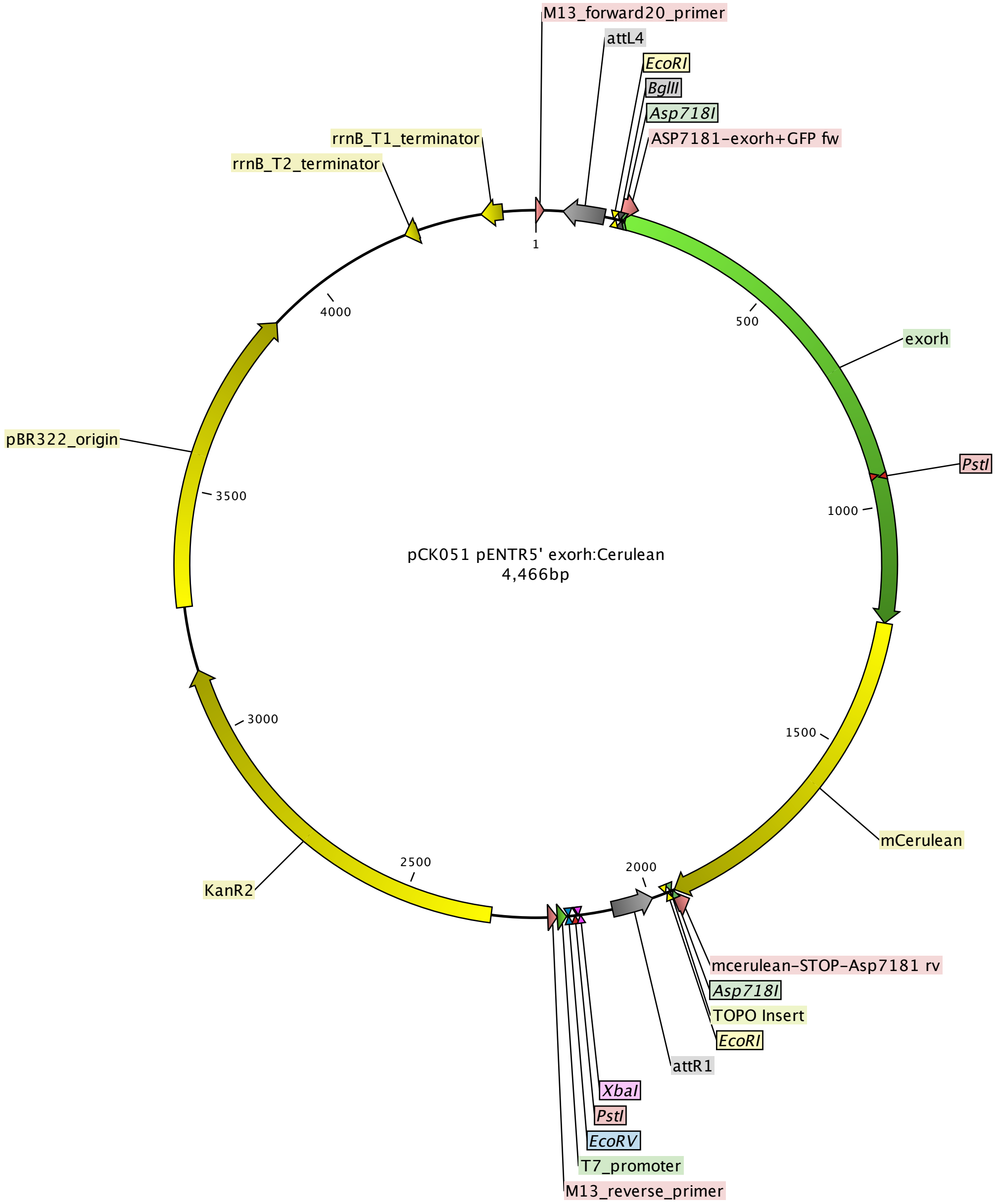

### pCK052 p5E exorhmCherry-.pdf

Kemmler et al.

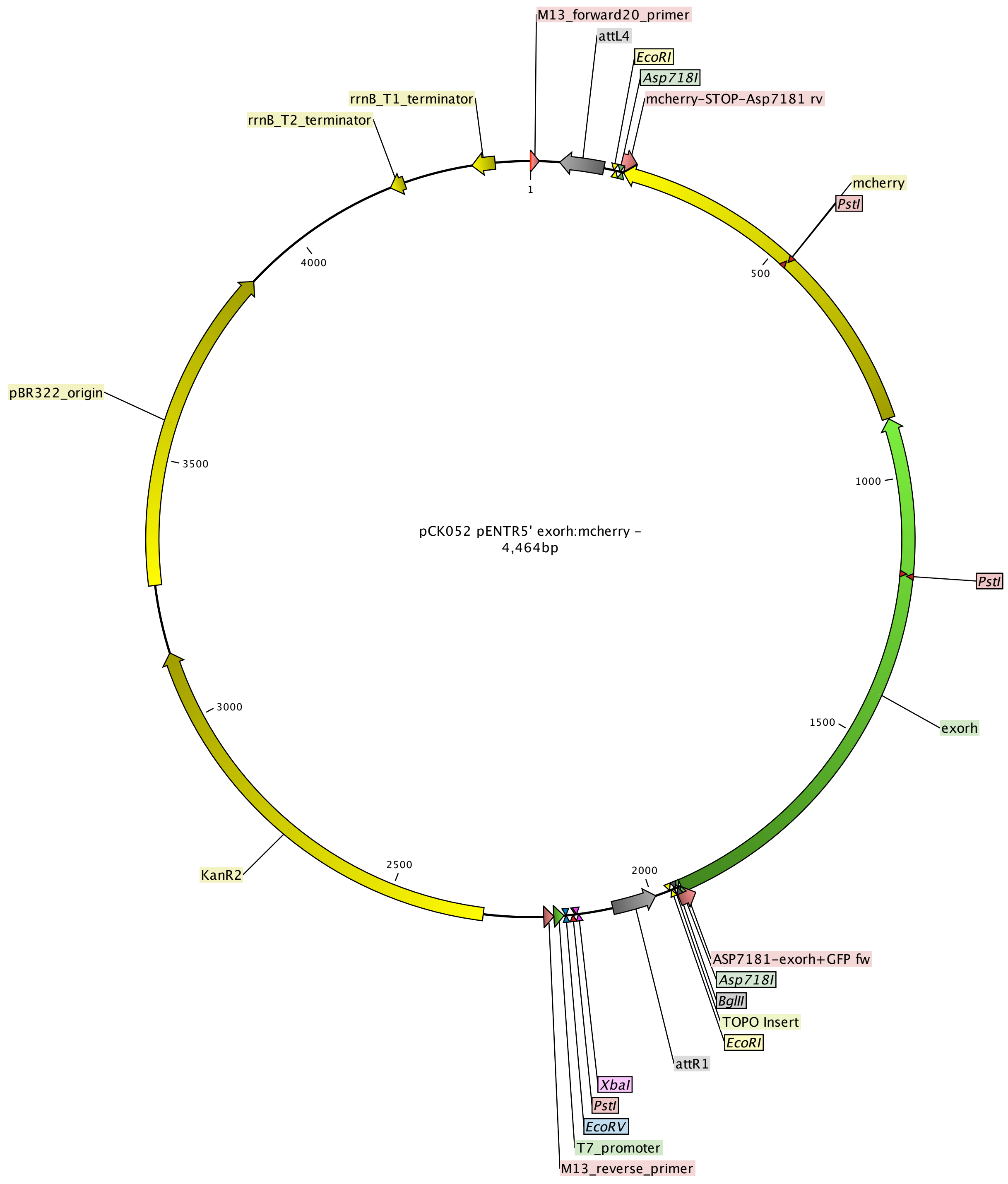

### pCK053 pDEST exorhmCherry-.pdf

Kemmler et al.

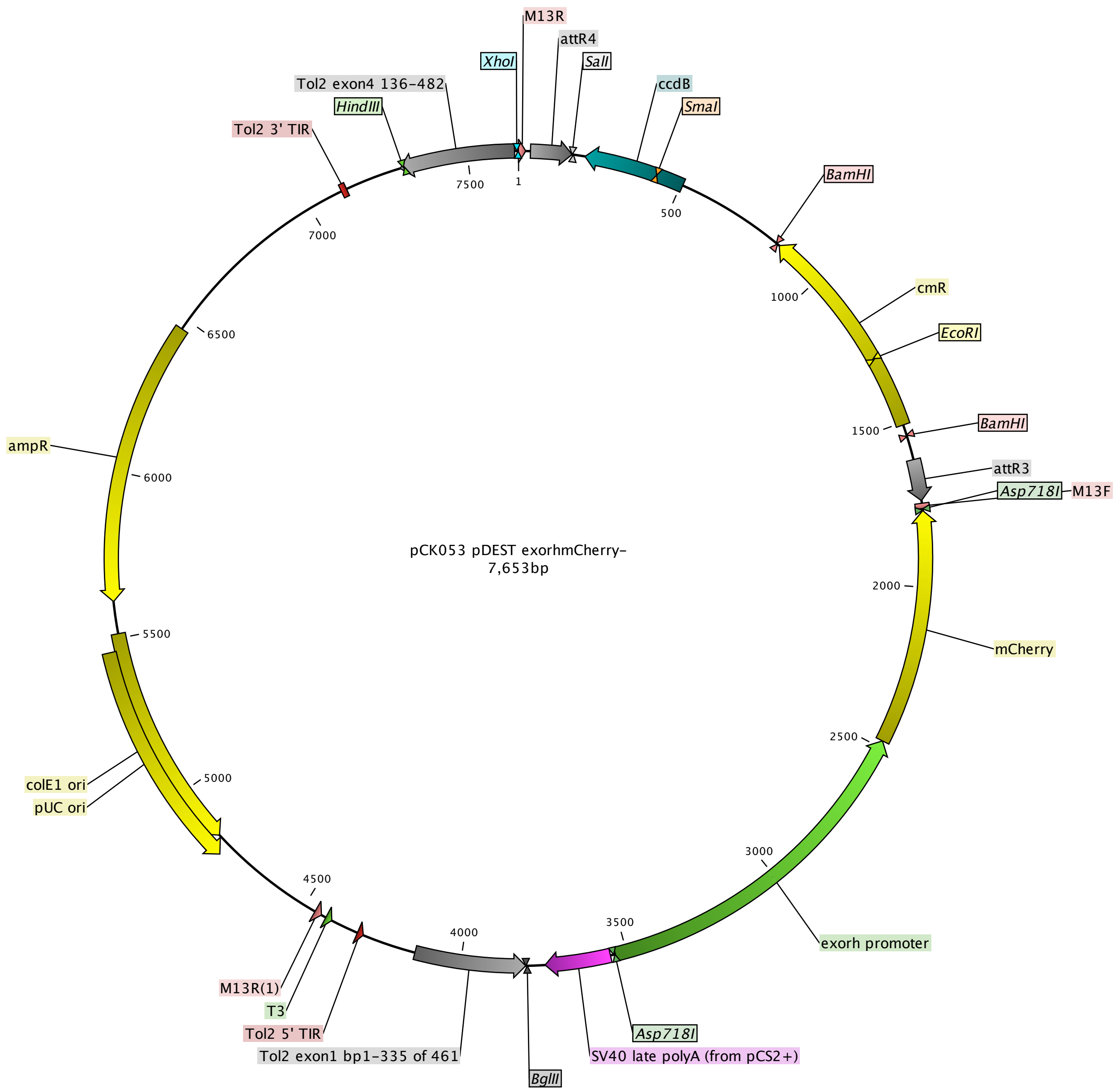

### pCK054 pDEST exorhmCerulean.pdf

Kemmler et al.

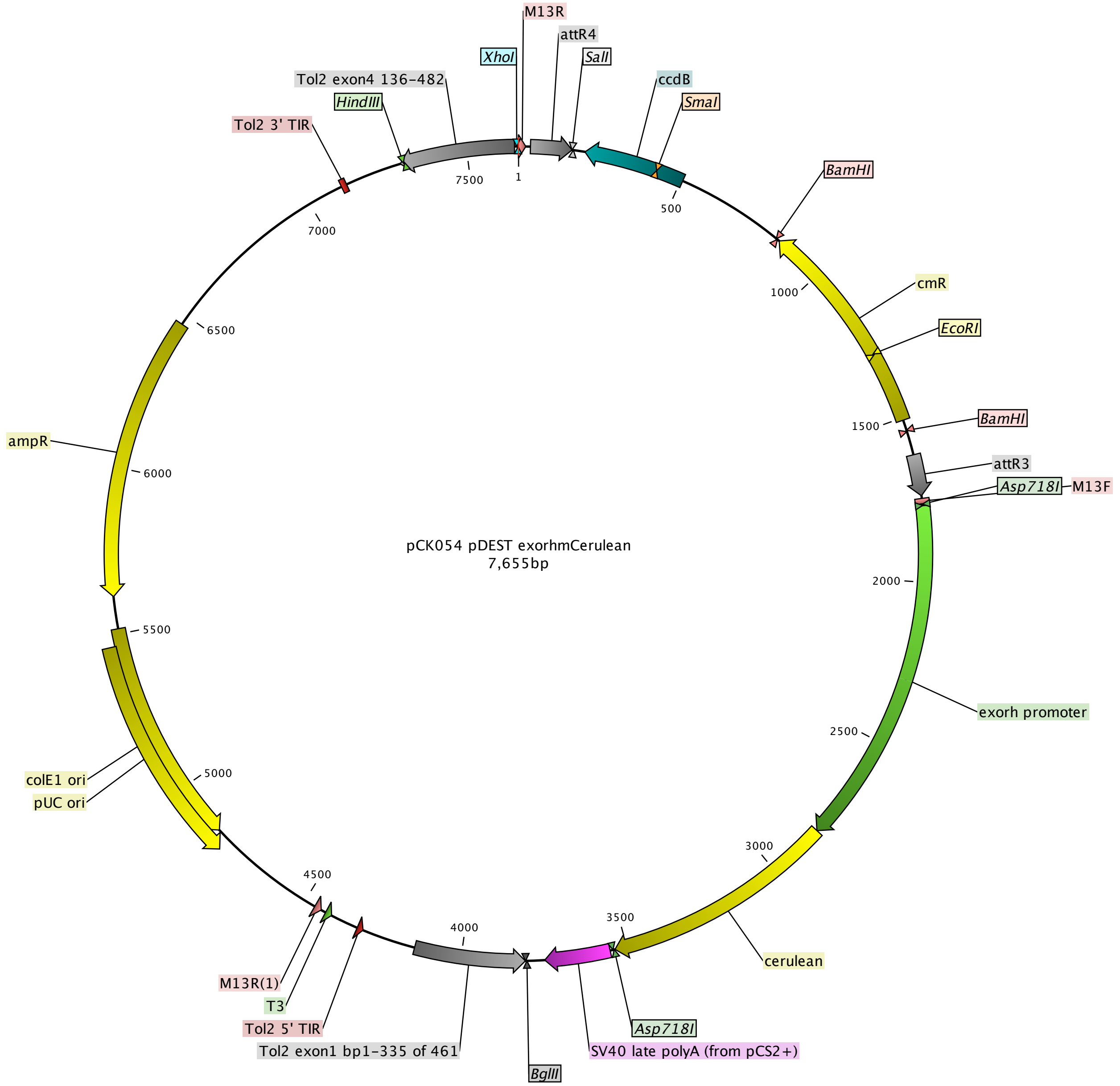

### pCK068 pME minmApple.pdf

Kemmler et al.

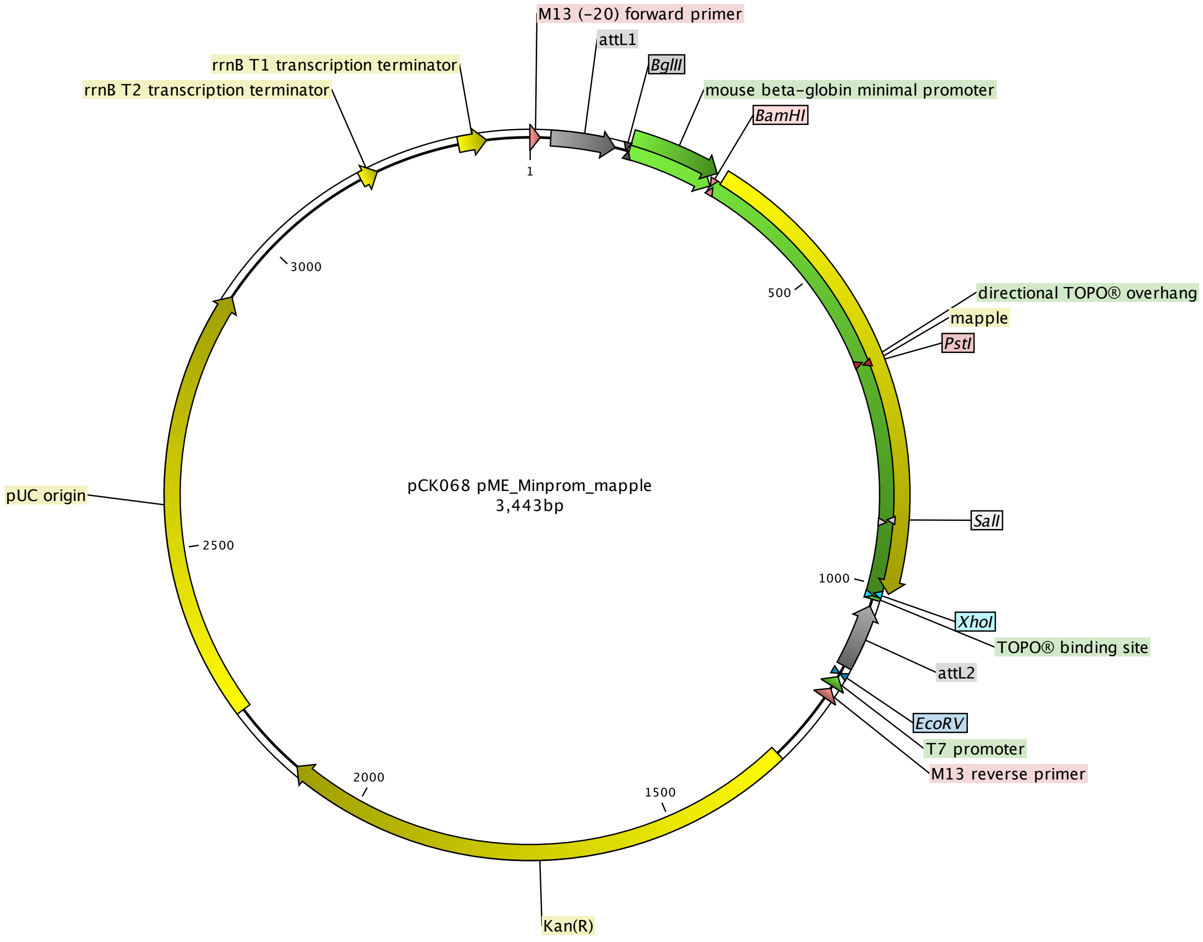

### pCK069 desminMCS-minmApple- exorhEGFP.pdf

Kemmler et al.

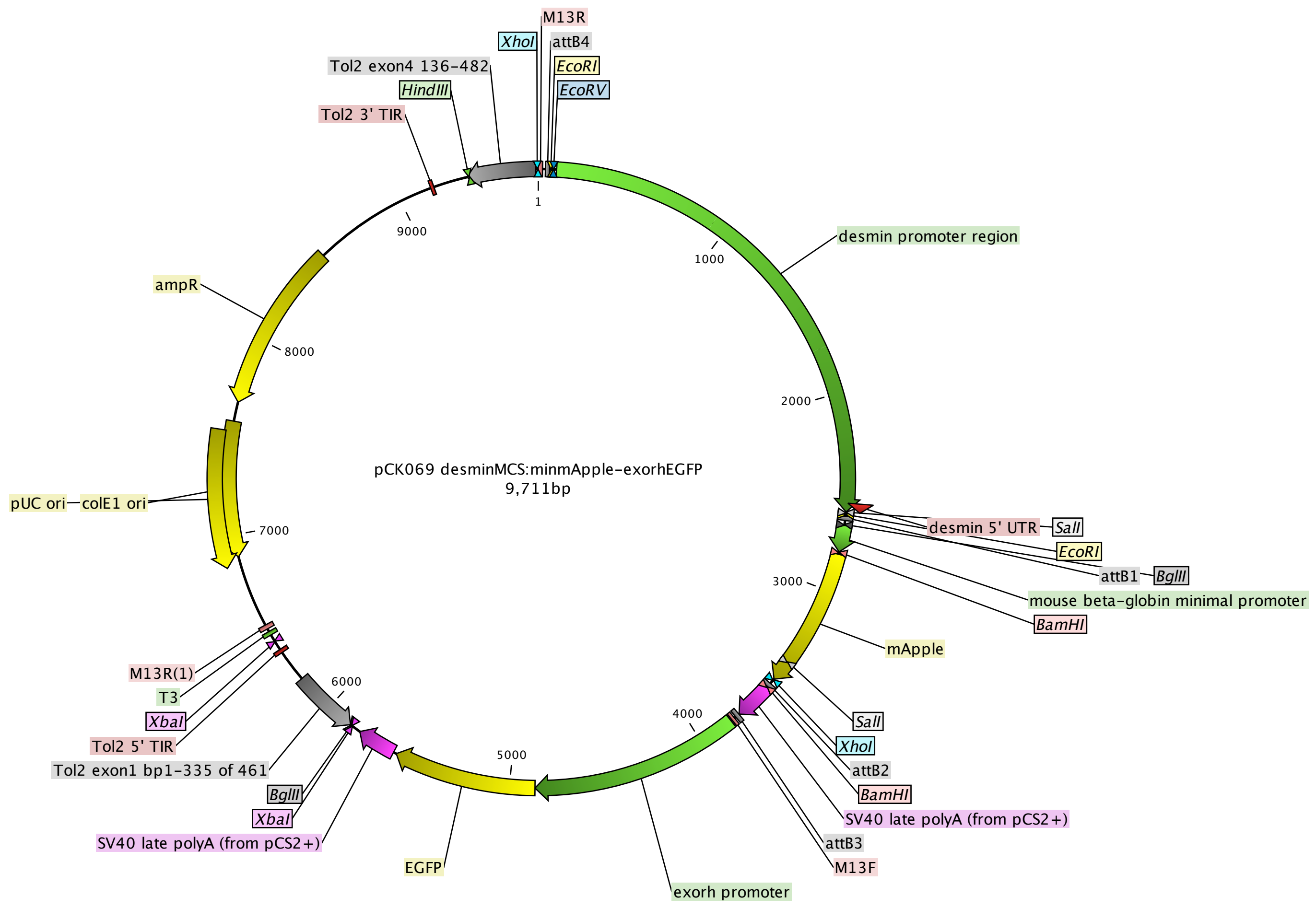

### pCK072 p5E 147bpexorhEGFP.pdf

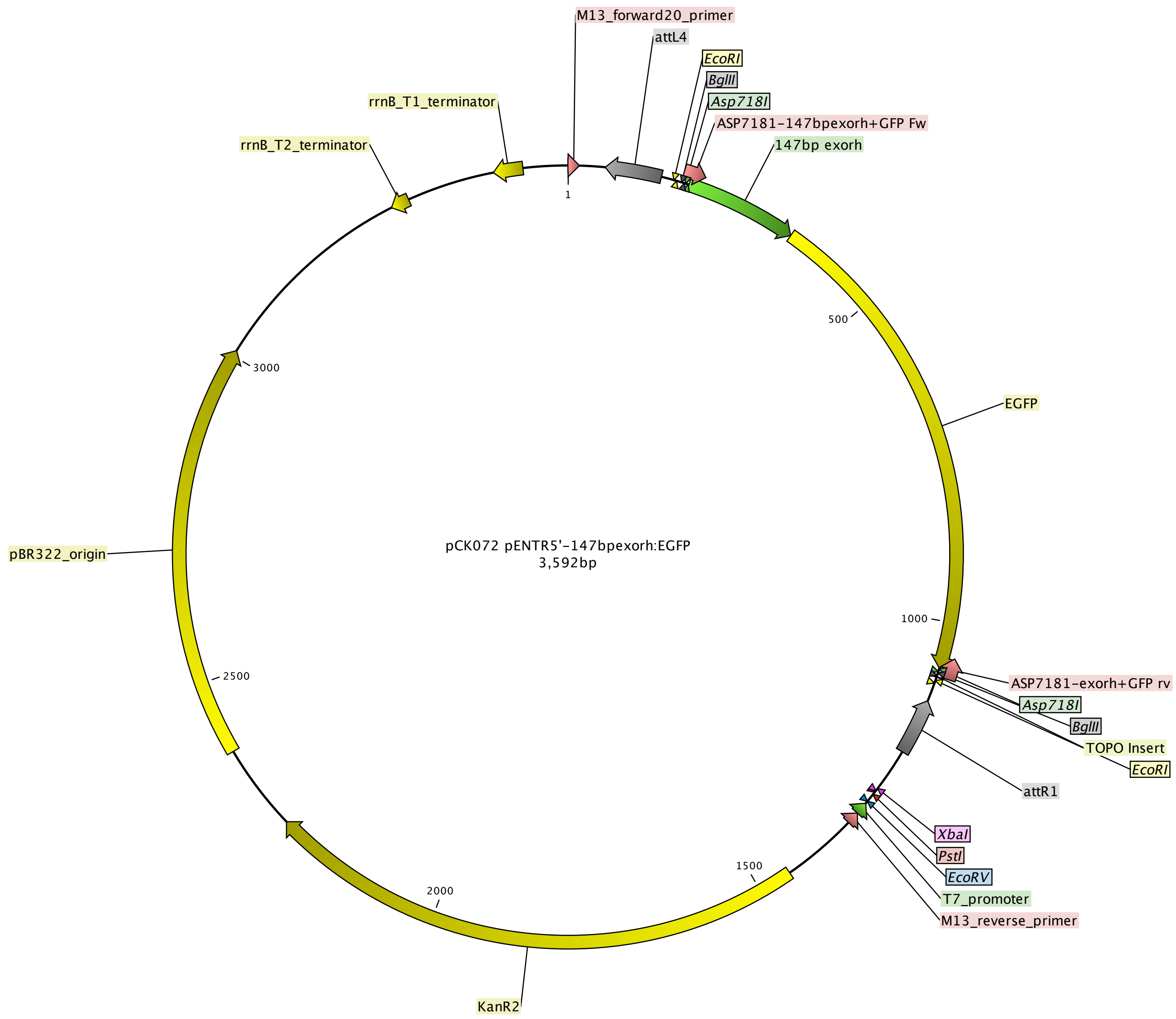

### pCK074 p5E exorhEGFP.pdf

Kemmler et al.

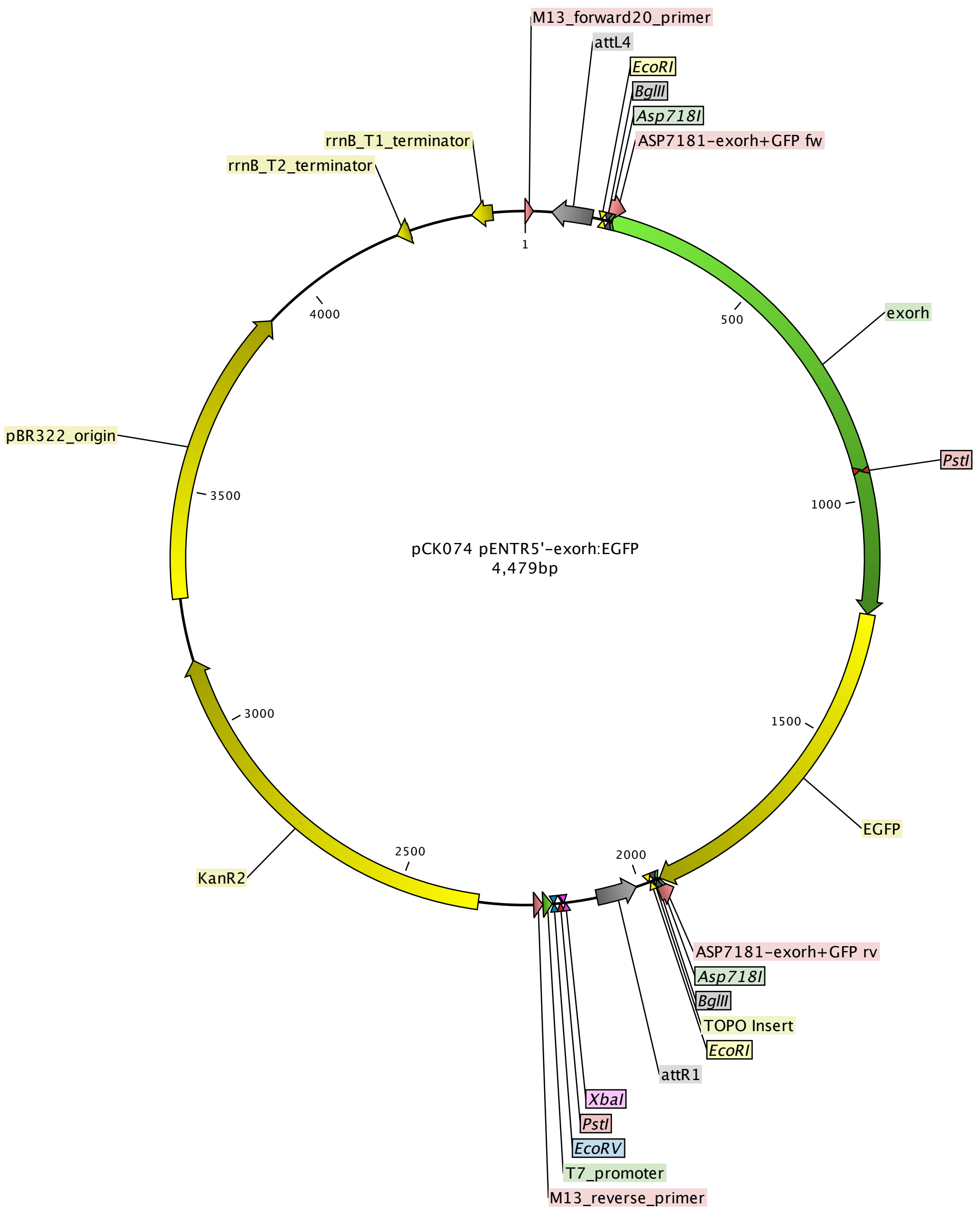

### pDEST_R4R3_exorhmCherry-cavefish.pdf

Kemmler et al.

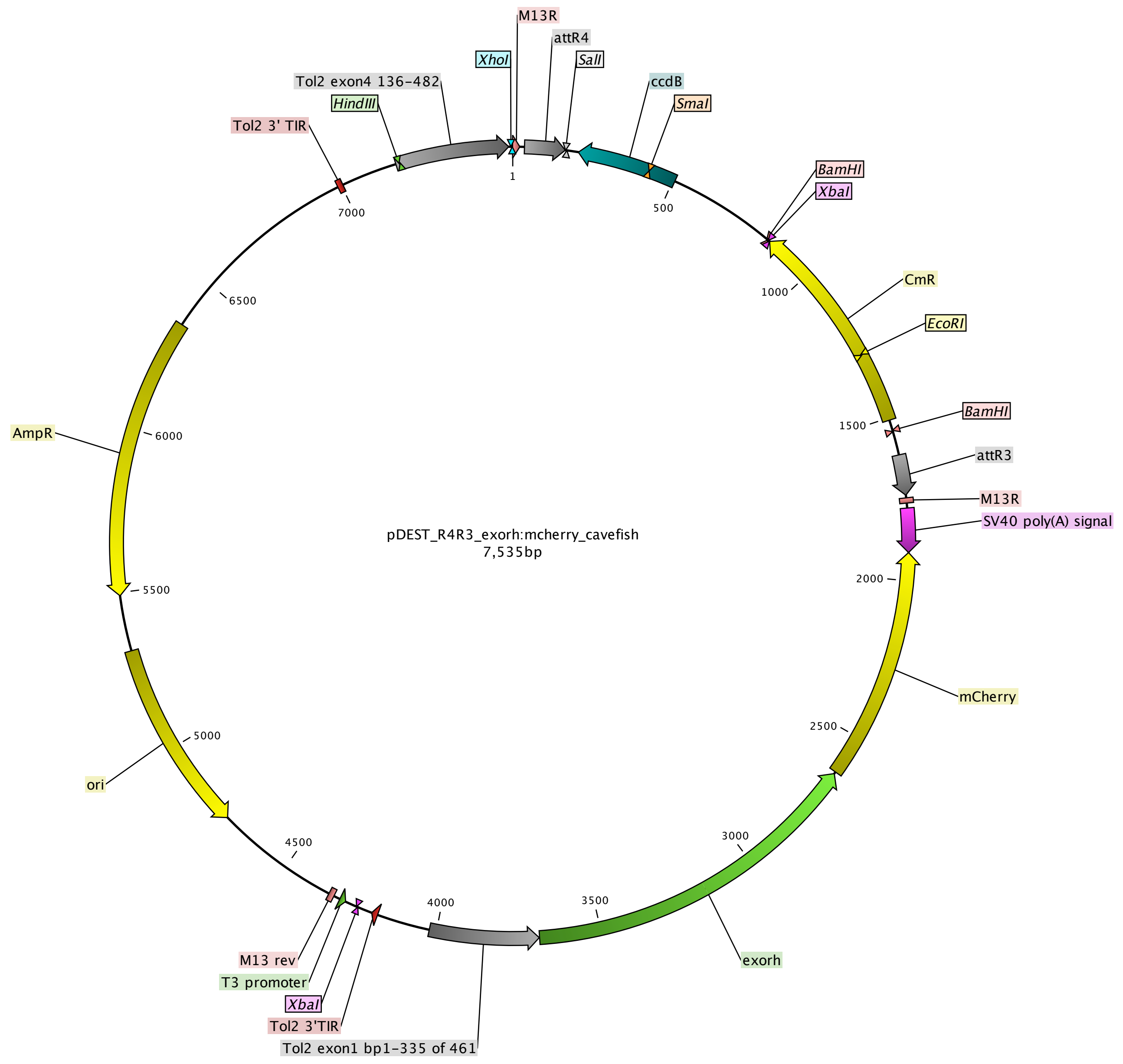

### pDEST_R4R3_exorhmGreenLantern_cavefish.pdf

Kemmler et al.

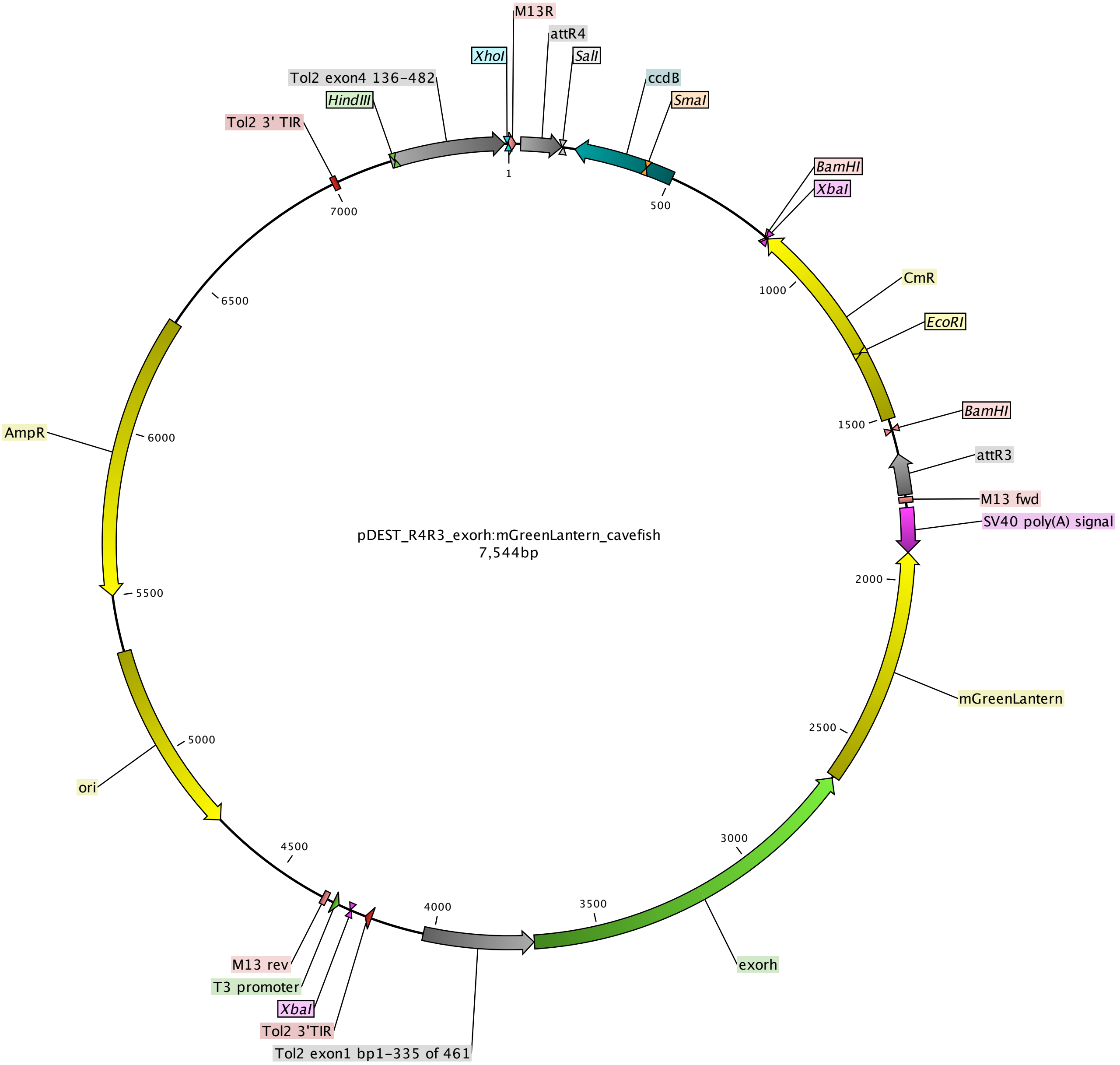

### pKD001 pME minEGFP.pdf

Kemmler et al.

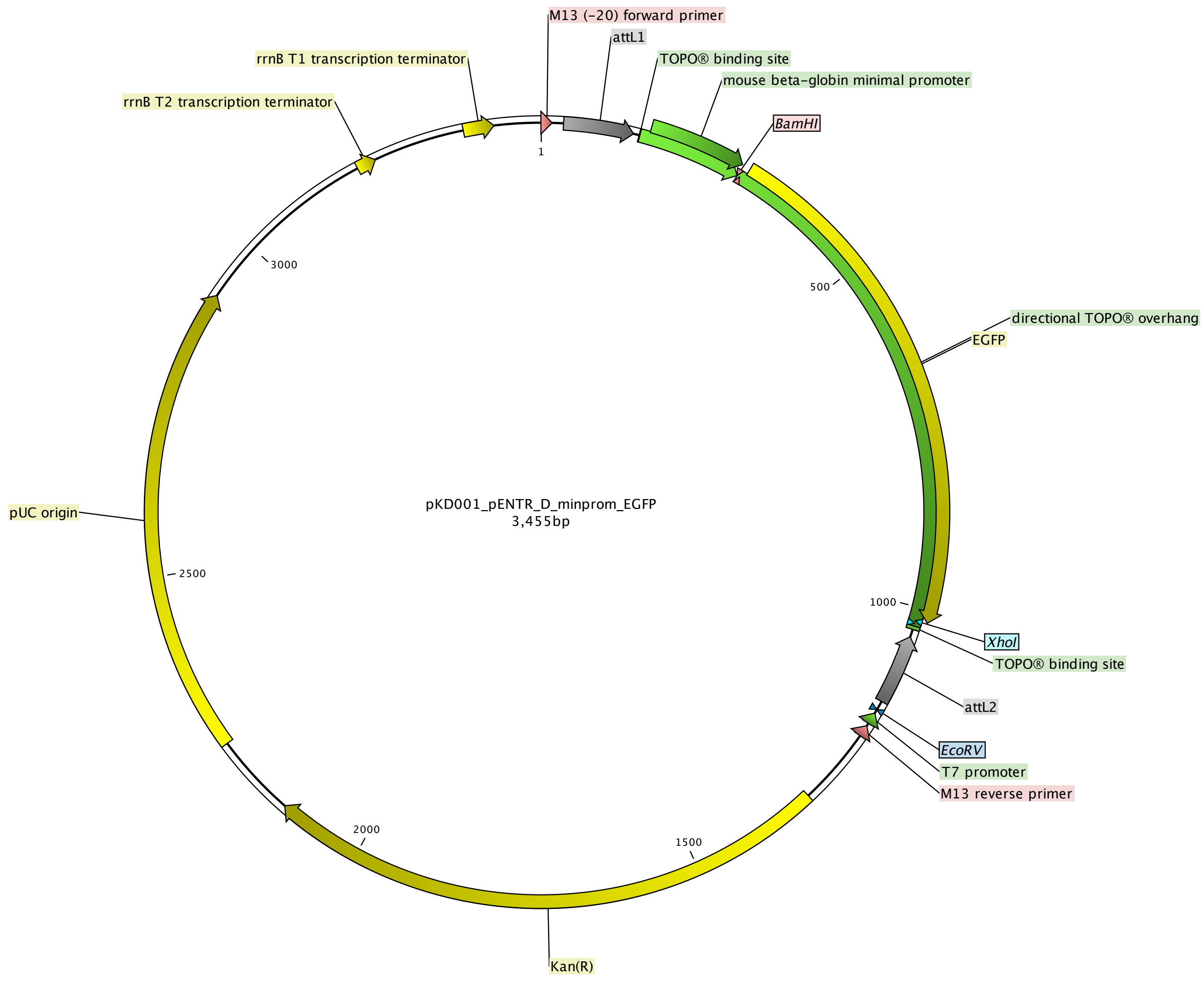

### pME min kalta4.pdf

Kemmler et al.

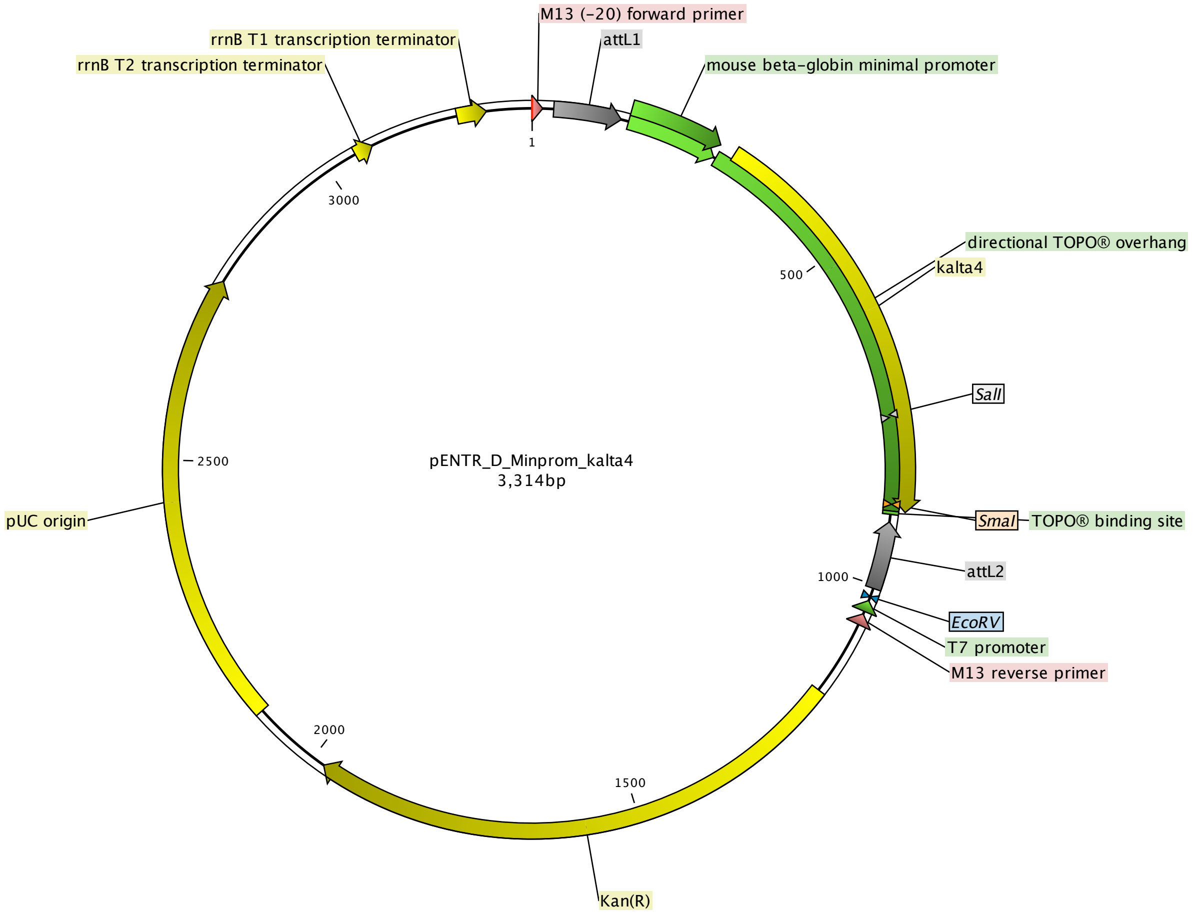

### pME minCreERT2.pdf

Kemmler et al.

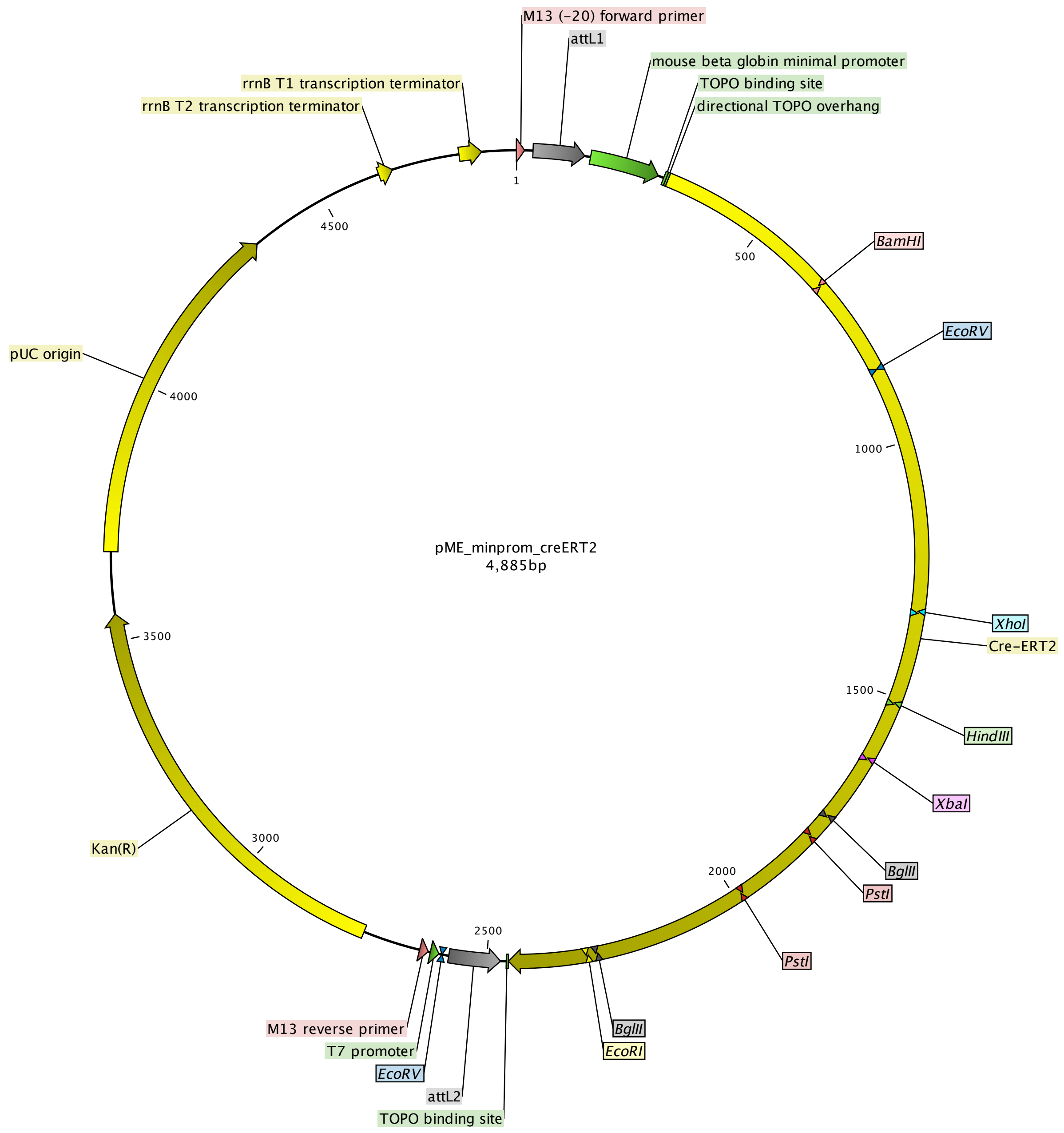

### pSN001 pME minmCerulean no stop.pdf

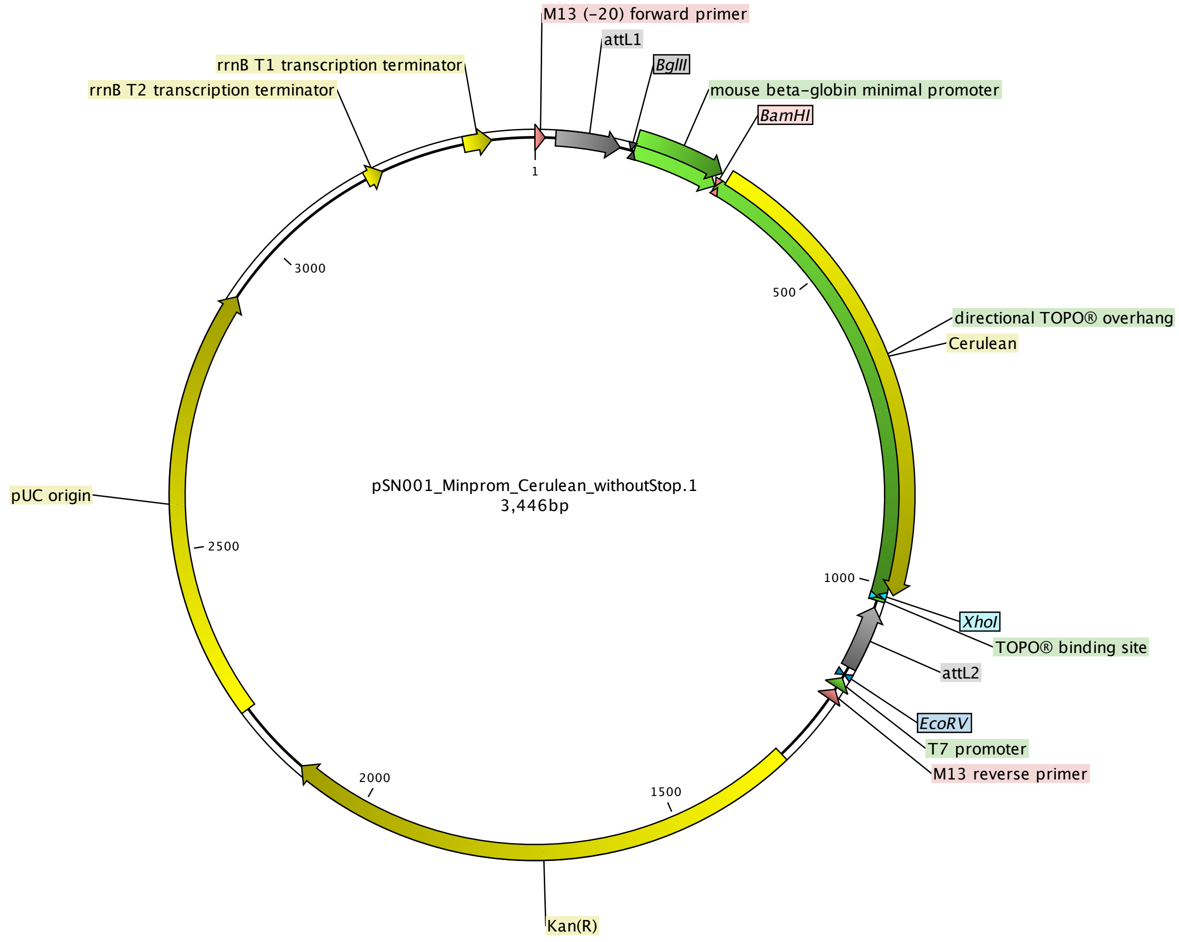
